# The mitotic spindle kinase MSK co-ordinates segregation of the nucleus and kinetoplast in *Leishmania mexicana*

**DOI:** 10.1101/2025.09.19.677349

**Authors:** Juliana B. T. Carnielli, Vincent Geoghegan, James A. Brannigan, Manuel Saldivia, Charlotte Hughes, Ana Paula C. A. Lima, Maria Cristina M. Motta, Adam Dowle, Anthony J. Wilkinson, Jeremy C. Mottram

## Abstract

Replication and segregation of the nucleus and kinetoplast, the mitochondrial DNA, are tightly coordinated in trypanosomatid parasites, but the signalling pathways that govern this process are unknown. Here, we characterise the mitotic spindle kinase (MSK), a key regulator of this coordination in *Leishmania*. Using chemical genetics, we engineered an analog-sensitive MSK to inhibit its activity. We show that inhibition of MSK impairs mitotic spindle elongation and blocks both nuclear and kinetoplast segregation, halting cell cycle progression and leading to cell death. We combined chemical genetics with proximity-based phosphoproteomics to identify four substrates: two GTPase-activating proteins, a nuclear segregation protein, and a hypothetical protein. We demonstrate that MSK co-localises with these four proteins in the nucleus, mitotic spindle, kinetoplast, and cytoplasm. Our findings establish MSK as a critical kinase that controls the co-ordinated segregation of the nucleus and kinetoplast, providing a new avenue for understanding cell cycle regulation in *Leishmania*.

## Introduction

In eukaryotes, the accurate inheritance of genetic material is fundamental for maintaining genome integrity and cellular viability. This process involves the precise replication and segregation of nuclear DNA (nDNA) as well as the faithful transmission of mitochondrial DNA (mtDNA) to daughter cells. These complex events are orchestrated by a dynamic interplay of molecular machinery, including the mitotic spindle and kinetochores for nuclear chromosome segregation^1^, and mitochondrial fission/fusion machinery for mtDNA inheritance^2,3^. Nuclear chromosome segregation requires the coordinated action of centromeres, kinetochores, and spindle microtubules, which are nucleated from centrosomes or other pathways in mammalian cells^4^. The kinetochore, a multiprotein complex, connects centromeric DNA to spindle microtubules, mediating chromosome movement^1,5^. A diverse array of motor and non-motor proteins, along with kinases and phosphatases like Aurora, Polo, and Wee1, dynamically regulate these events via phosphorylation, ensuring precise spatial and temporal control of cell division^6,7^. Unlike nuclear DNA, mtDNA exists as circular double-stranded molecules organized into nucleoids, which are distributed throughout the mitochondrial matrix^8^. In most eukaryotic cells, mtDNA replication is not synchronized with the cell cycle. Instead, replicated nucleoids are partitioned into daughter mitochondria through a dynamic process of mitochondrial fission and fusion, ensuring their proper inheritance during cytokinesis^2,3,9^.

The protozoan parasite *Leishmania* spp., a member of the early-branching kinetoplastid lineage, presents an exception to these general eukaryotic models. These parasites possess an unusual cell biology, including a single mitochondrion containing a massive and intricate kinetoplast DNA (kDNA) network. The kDNA is a catenated structure composed of thousands of minicircles and dozens of maxicircles, which must be accurately replicated and segregated along with the nuclear genome^10^. In *Leishmania*, nuclear and kinetoplast DNA duplication are tightly synchronized, with S-phase occupying approximately 40% of the cell cycle. This is followed by a brief G2 phase, leading to the near-simultaneous segregation of both organelles. While segregation of nuclear DNA is temporally coupled with kinetoplast segregation, whether one precedes the other remains unknown^11–15^.

Despite retaining conserved eukaryotic cell-cycle regulators such as CDK/cyclin complexes and Aurora B kinase, kinetoplastids lack several key components of the canonical segregation machinery. They undergo a closed mitosis, where the nuclear envelope remains intact, and they lack canonical kinetochores and the centromeric histone H3 variant, CENP-A. Instead, they rely on a lineage-specific set of proteins known as kinetoplastid kinetochore proteins (KKTs) and their interactors KKIPs to mediate chromosome-spindle attachment^16–22^. Furthermore, the mechanism of spindle nucleation is not fully understood, as centrioles and spindle pole bodies have not been identified. Kinetoplast DNA segregation, in turn, is a highly specialized process mediated by the tripartite attachment complex (TAC), a structural bridge that tethers the kDNA network to the basal body of the flagellum^23^.

Despite significant progress in identifying the unique proteins and structures involved in nuclear and kinetoplast segregation in trypanosomatids, the precise regulatory mechanisms governing these processes remain poorly understood. A deeper investigation into the protein kinases that coordinate this dual segregation is essential. Here, we investigate the function of a key mitotic kinase in *Leishmania*, the Mitotic Spindle Kinase (MSK). Using analogue-sensitive mutants, we demonstrate that MSK activity is crucial for spindle microtubule elongation and for the proper segregation of both nuclear and mitochondrial DNA. Furthermore, using spatial phosphoproteomics, we identify several novel spindle-associated proteins, including Rab3GAP, ARFGAP1, a nuclear segregation protein, and the hypothetical protein LmxM.36.2560, as substrates of MSK, providing new insights into the regulatory network controlling cell division in *Leishmania*.

## Results

### Mitotic spindle kinase, MSK, co-ordinates *Leishmania* mitochondrial and nuclear DNA segregation

In our previous analysis of the *L. mexicana* kinome, we identified four protein kinases – LmxM.24.0670, LmxM.09.0310 (CRK12), LmxM.08_29.2150 (CRK10) and LmxM.26.2440 (AUK3) – that exhibited primary localisation to the nucleus and secondary localisation to the kinetoplast^24^. For this study, we focussed on LmxM.24.0670 hereafter referred to as <u>m</u>itotic <u>s</u>pindle <u>k</u>inase (MSK) since the phenotype of CRK10 and AUK3 null mutants were reported in Baker et al. (2021)^24^ and CRK12 has been investigated elsewhere^25^. MSK is a CMGC family protein kinase conserved across trypanosomatids, including all *Leishmania* species, and both African and South American trypanosomes (Supplementary Fig. 1).

We were unable to generate MSK promastigote form gene deletion mutants, suggesting that the kinase is essential for cell-cycle progression. To test this hypothesis, we adopted an analog-sensitive (AS) approach to selectively inhibit MSK activity^26^. Using an AlphaFold 3 structural model (Fig. 1a), we predicted the gatekeeper residue and replaced the native methionine (M192) with glycine or alanine via CRISPR-Cas9-directed precision genome editing^27^ (Supplementary Fig. 2a). Editing was confirmed by PCR followed by restriction endonuclease digestion and sequencing (Supplementary Fig. 2b – d). Promastigotes expressing the glycine-substituted AS MSK display growth rates comparable to the parental T7/Cas9 line, indicating that gatekeeper mutation does not impair parasite fitness *in vitro* culture (Supplementary Fig. 3).

**Fig. 1.**
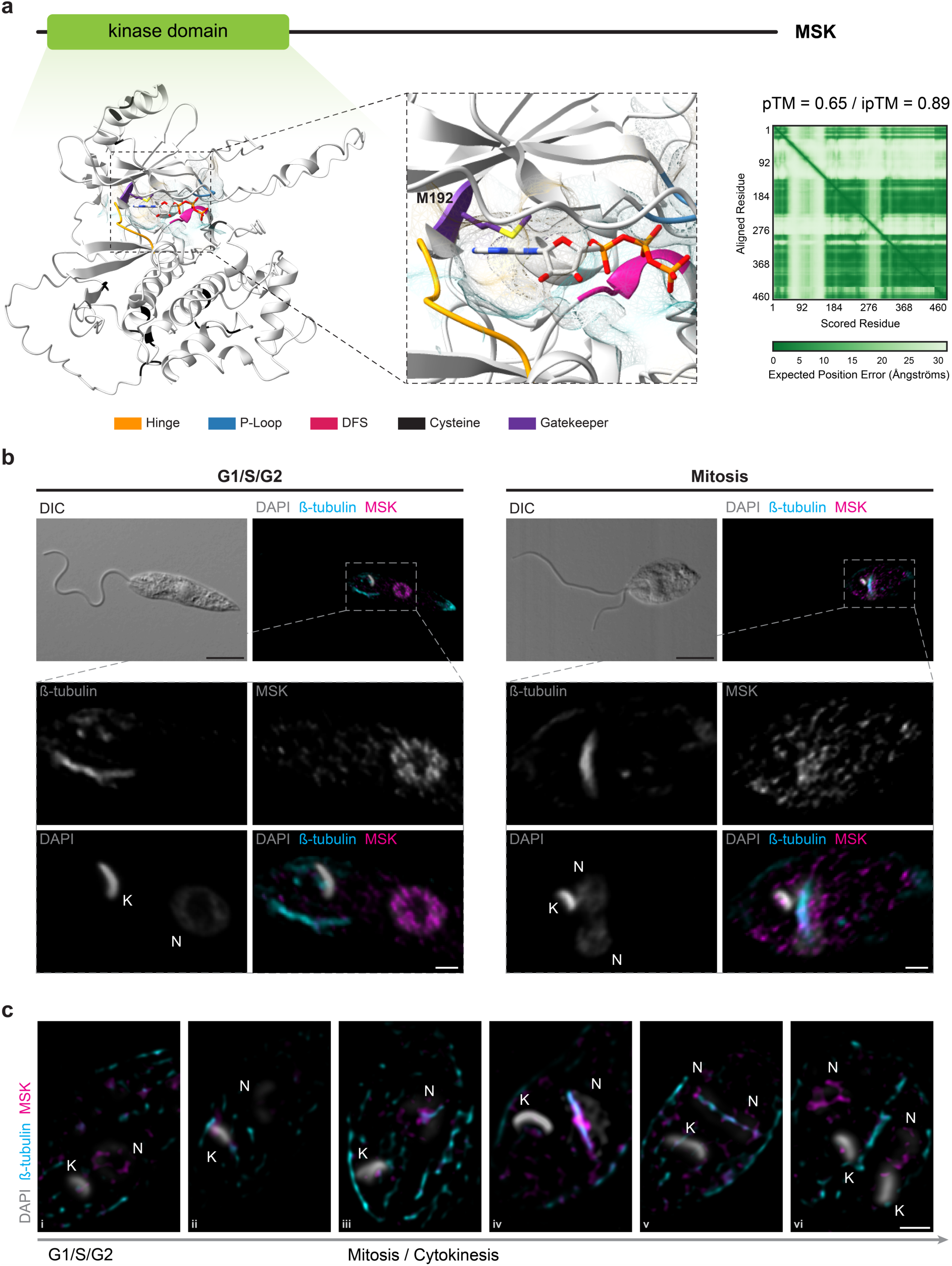
MSK kinase domain architecture and MSK localisation in *L. mexicana* promastigotes. **a** Schematic showing *Leishmania* MSK protein domain architecture predicted by online InterPro 103.0 tool^29^ and the predicted structural model of the MSK kinase domains with ATP ligand, generated by AlphaFold 3^30^ and visualized with ChimeraX v1.9. The ATP ligand is displayed as a stick model, with heteroatom-based colouring. A semi-transparent molecular lipophilicity potential surface is overlaid, with colour ranging from dark cyan (most hydrophilic) to dark golden (most lipophilic). Inset: detailed view of the gatekeeper residue (M192) in the hydrophobic region of the ATP-binding pocket. AlphaFold confidence metrics are displayed on the right: predicted template modelling score (pTM), the interface predicted template modelling score (ipTM) between the kinase and ATP, and the predicted aligned error (PAE) plot. Confocal (**b**) and high resolution (**c**) microscopy of *L. mexicana* promastigotes expressing 3xMyc::mNG::MSK_AS. Cells were stained with KMX-1 antibody to label β-tubulin, anti-Myc to label the endogenously tagged 3xMyc::mNG::MSK_AS, and counterstained with DAPI to visualise DNA. Representative fluorescence micrographs showing parasites in different cell cycle stages. The channel and the colours used for each marking are indicated on the top or the left. K, kinetoplast; N, nucleus. Individual channels and other merged channel patterns are shown in Supplementary Fig. 5 – 6. Scale bars, 5 μm.

To investigate MSK expression during the cell cycle, we generated an endogenous N-terminally tagged MSK fusion (3xMyc::mNeonGreen) in the MSK AS mutant background (Supplementary Fig. 4). Confocal microscopy confirmed MSK localisation to both the nucleus and the kinetoplast (Fig. 1b). High-resolution imaging of β-tubulin, MSK, and DNA across the cell cycle revealed that MSK is present in the cytoplasm, nucleus, and kinetoplast, consistently in close proximity to β-tubulin (Fig. 1c). Prior to mitosis, MSK exhibits a diffuse nuclear distribution (Fig 1c, i – ii). At mitotic onset, coinciding with spindle organisation, MSK accumulates within the nucleus. At this stage, β-tubulin forms a central ring-like structure, around which MSK is arranged (Fig. 1c, iii). During anaphase, MSK concentrates along the central portion of the elongating spindle, adopting a helical configuration. Interestingly, continuous points of connection are observed between MSK on the intranuclear spindle and β-tubulin on the subpellicular microtubules (Fig. 1c, iv), raising the possibility that the microtubule organizing centre (MTOC) in *Leishmania* is not fully intranuclear, in contrast to current models in *T. brucei*^28^. Following completion of DNA segregation, but prior to the end of cytokinesis, MSK redistributes to the periphery of the separated nuclei (Fig. 1c, v – vi).

The analog-sensitive kinase approach allows selective inhibition of an engineered kinase by cell-permeable bumped kinase inhibitors (BKIs) that do not bind to the wild-type enzyme^26,31,32^ (Fig. 2a). To determine whether MSK kinase activity is required for *L. mexicana* survival, we performed viability assays with BKIs 1NM-PP1 and 1NA-PP1 against both promastigote and intracellular amastigote stages, comparing the parental T7/Cas9 line with the MSK AS mutants. The two inhibitors differ by a single methylene group, with 1NM-PP1 exhibiting greater potency, particularly against the AS kinase harbouring glycine residue at the gatekeeper position (Fig. 2b). By contrast, 1NA-PP1 displays reduced specificity, with no significant difference in susceptibility between the glycine and alanine AS variants.

**Fig. 2.**
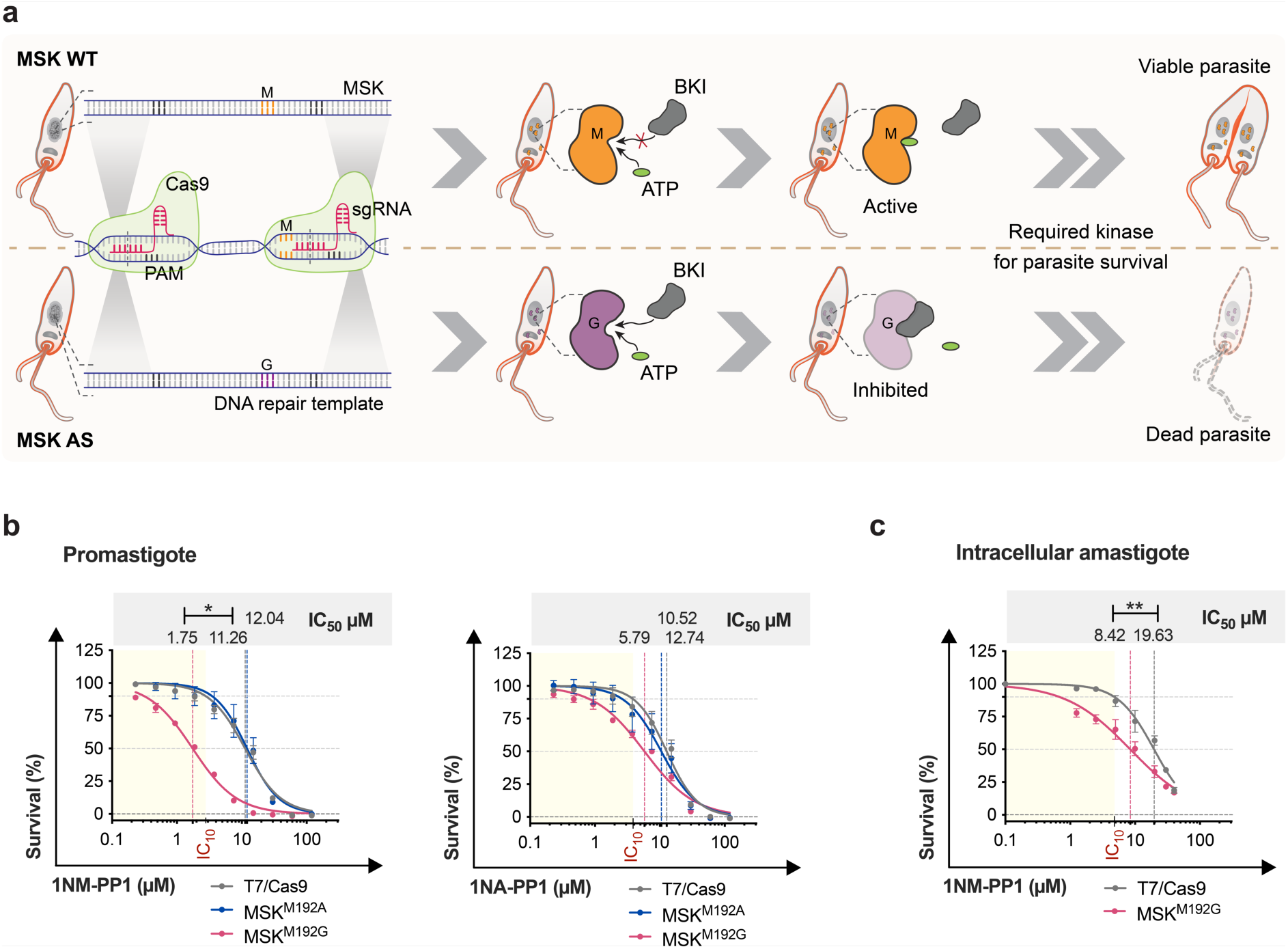
Cell viability of MSK analog-sensitive and progenitor line growing under bumped kinase inhibitors pressure. **a** Schematic representation of the viability assay performed on parasites expressing either the wild-type or the MSK AS variant, illustrating the expected outcome if the targeted protein kinase is essential for parasite survival following treatment with a bumped kinase inhibitor. The same experimental principle was applied to the amastigote viability assay. WT, wild type; AS, analog-sensitive; Cas9, endonuclease; sgRNA, single guide RNA; PAM, protospacer adjacent motif; M, methionine; G, glycine; BKI, bumped kinase inhibitor. **b** Parasites harbouring analog-sensitive kinases for MSK had their susceptibility to the1NM-PP1 and 1NA-PP1 BKI assessed by resazurin assay in promastigotes. **c** Parasites containing glycine analog-sensitive kinase for MSK had their susceptibility to 1NM-PP1 BKI assessed in intracellular amastigote stage by percentage of infected macrophage. Fitting of dose-response curves was carried out using GraphPad Prism v10.4.1, considering the untreated control for each cell line as 100% viability. BKI half maximal inhibitory concentration (IC_50_), calculated using the dose-response-inhibition model in GraphPad Prism v10.4.1, are shown above each graph in the grey area. Data represents mean ± SEM from three independent biological replicates. P values were calculated using unpaired two-tailed Student’s t-tests comparing each cell line with the T7/Cas9 parental cell line. * p-value <0.05; ** p-value <0.01. IC_10_ highlighted in red corresponds to the BKI concentration that impairs the T7/Cas9 progenitor viability by 10%. The yellow area covers the BKI concentration below the IC_10_.

Parasites expressing the glycine AS variant of MSK were significantly more sensitive to 1NM-PP1 treatment in both life stages. In the promastigotes, IC_50_ values are 1.8 µM for MSK^M192G^ versus 11.3 µM for the parental T7Cas9. In the amastigotes, IC_50_ values are 8.4 µM for MSK^M192G^ and 19.6 µM for T7Cas9. These findings indicate that MSK catalytic activity is required for the survival of *L. mexicana* in both insect and mammalian stages (Fig. 2b – c).

To assess if MSK kinase activity is required for cell-cycle progression, we analysed DNA content in promastigotes by flow cytometry following inhibition with 5µM 1NM-PP1. After six hours of treatment, MSK AS promastigotes accumulate in G2/mitosis, while untreated MSK AS and parental T7/Cas9 cells retain a normal cell cycle profile (Fig. 3a, left panel and Supplementary Fig. 7). This arrest persists for at least 24 h. Flow cytometry plots of DNA content versus forward scatter reveal that G2/mitosis arrested cells are larger than the control cells at the same stage (Fig. 3a, right panel and Supplementary Fig. 7). To investigate this arrest in more detail, we employed fluorescence microscopy to monitor cell cycle progression by staining for β-tubulin (marking the mitotic spindle) and DNA, as described previously^27^. Fluorescence microscopy shows that 6 h inhibition of MSK causes the accumulation of a distinct mitotic stage of the *Leishmania* cell cycle (hereafter termed sub-stage 3.1). At this stage, promastigotes display a fully elongated cell body with two long flagella and the morphology of two nascent daughter cells, yet lack nuclear or kinetoplast segregation (Fig. 3b – c). Quantification showed that 28.6% of cells are arrested in sub-stage 3.1 after 6 h of MSK inhibition, with 73% of these displaying β-tubulin concentrated in a short nascent spindle, consistent with a role for MSK in early mitosis, spindle elongation, and subsequent DNA segregation (Fig. 3c). After 24 h of inhibition, 16.1% of the cells remain in sub-stage 3.1, but one flagellum has shortened markedly (termed sub-stage 3.1*), and only 56.1% of these cells still exhibit a visible mitotic spindle. Moreover, the majority (62.4%) of cells display morphology resembling two daughter cells but, distinct from the sub-stage 3.1*, lack the mitotic spindle (termed sub-stage 3.2) (Fig. 3c). Importantly, none of these atypical stages are observed in untreated controls. Prolonged MSK inhibition also led to a significantly higher number of atypical cells in cytokinesis. After 24 h, 29.4% of cell display a cleavage furrow progressing from the posterior end towards the anterior end of the cell, generating a cell body without DNA-containing structures: 91.9% of these cells are found at sub-stage 3.2. Because 11% of untreated MSK AS cells also exhibit a posterior furrow, we assessed this in the parental T7/Cas9 line and found no significant difference from the untreated MSK AS (Fig. 3d – e).

**Fig. 3.**
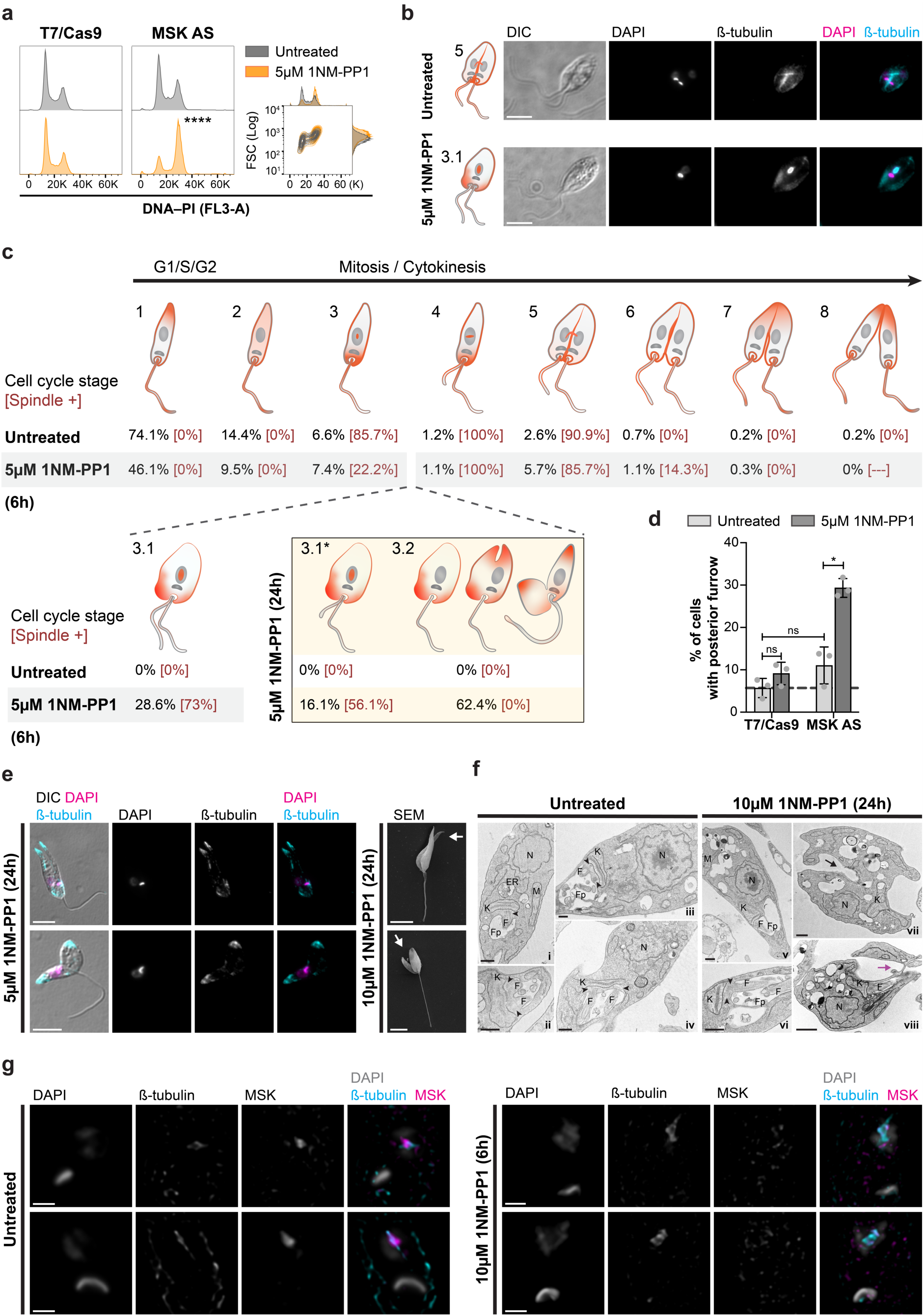
Mitotic spindle kinase regulates mitochondrial and nuclear DNA segregation during the *Leishmania* cell cycle. **a** Representative cell cycle-profile histogram of cells stained with propidium iodide (PI) after 6 h of treatment with 1NM-PP1. Right panel, adjunct cell cycle-profile and forward scatter (FSC). P values were calculated using two-tailed Student’s t-tests comparing the percentage of cells between MSK AS treated and untreated: ****, p = 0.0000007. **b** Representative fluorescence micrographs showing parasites in cell cycle stages 3.1 and 5 (see panel ’c’) after 6 h of treatment of MSK AS with 1NM-PP1. Scale bars, 5 μm. DIC, the Nomarsky differential interference contrast. **c** Schematic of kinetoplast/nucleus (grey) configuration and β-tubulin (orange) distribution through the cell cycle of *L. mexicana*. Percentage of cells in each stage [percentage of cell with mitotic spindle, Spindle +] were measured for MSK AS parasites growing in the presence or absence of 1NM-PP1 for 6 h (n > 425 cells) and 24 h (n > 200). **d** Percentage of cells with furrow formation progressing from the posterior end towards the anterior end, after 24 h incubation of MSK AS with 1NM-PP1. Data represents mean ± SEM. P values were calculated using two-tailed Student’s t-tests from three replicates (n > 80 in each replicate): *, p = 0.0058; ns, not statistically significant. **e** Fluorescence (on the left) and scanning electron microscopy (SEM, on the right) of promastigotes after 24 h incubation with 1NM-PP1. Representative micrographs showing parasites in the cell cycle stage 3.2 with division furrow forming at the posterior end of the cell (top panels) and progressing to form structure lacking DNA-containing organelles (bottom panels). Scale bars, 5 μm. **f** Representative micrographs using transmission electron microscopy (TEM) of MSK AS parasites after 24 h incubation with or without 1NM-PP1. N, nucleus; K, kinetoplast; M, mitochondrion; F, flagellum; Fp, flagellar pocket. Black arrow points to the division furrow at the posterior end of the cell and, purple arrow indicates an advanced stage of cytokinesis initiated at the posterior end. Arrowheads point to the basal bodies. Scale bars, 1 μm. **g** High resolution fluorescent microscopy of promastigotes with and without 6 h incubation with 1NM-PP1. Representative fluorescence micrographs showing parasites at the early stage of mitosis. The colours used for each marking are indicated on the top. Scale bars, 1 μm.

A further investigation, by transmission electron microscopy (TEM), after 24 h of incubation with 10 µM 1NM-PP1, revealed ultrastructural changes in MSK inhibited cells. In untreated cells, cells in G1-S have a nucleus, a kinetoplast and one flagellum contained in the flagellar pocket (Fig. 3f, i). As the cell cycle progresses, the kinetoplast elongates, a second basal body appears from which a new flagellum emerges and a duplicate flagellar pocket is formed (Fig. 3f, ii – iv). When MSK kinase activity was inhibited, cells had two flagella emerging from the same flagellar pocket, along with an additional structure resembling either a second, empty flagellar pocket or an expanded contractile vacuole adjacent to the original pocket. These cells also have a single nucleus and a single kinetoplast, which fail to elongate (Fig. 3f, v – vi). The TEM also corroborates the finding that MSK kinase inhibition leads to the generation of a partially formed cell without DNA containing structures (Fig. 3f, vii – viii).

Following MSK kinase inhibition, β-tubulin rearrangement in the mitotic spindle is blocked in its initial ring-like structure and MSK fails to position itself around it, remaining more dispersed in the nucleus (Fig. 3g and Supplementary Fig. 8). Taken together, our data reveal that inhibition of MSK activity impairs mitotic spindle elongation, as well as nuclear and mitochondrial DNA segregation, irreversibly compromising cell-cycle progression and leading to cell death.

### Combined chemical genetics and proximity phosphoproteomics to identify MSK kinase substrates

We have recently adapted cross-link BioID (XL-BioID) proximity labelling to map the proximal environment of target proteins in *Leishmania*^33^. To identify substrates of MSK, we combined chemical genetics with proximity labelling^34^ to compare the proximal proteome and phosphoproteome of MSK, in the presence and absence of 1NM-PP1. We used Cas9-mediated genome engineering to endogenously N-terminally tag both alleles of the MSK AS with the biotin ligase variant, miniTurbo (mT) (Supplementary Fig. 9), which enables faster proximity biotinylation than BirA^35^. A cell line expressing the KKT2 AS kinase fused to mT at its C-terminus was used as spatial reference control. Because both MSK and KKT2 are essential protein kinases in *Leishmania*^24^, successful recovery of mutants with both alleles endogenously tagged indicates that the tagged proteins retained functionality. Promastigotes expressing 3xMyc::mT::MSK_AS and KKT2_AS::mT::3xMyc were cultured in the presence or absence of 1NM-PP1 for 2 h, during which the biotinylation step was performed. Using our previous workflow that allows enrichment of proximal proteins and their phosphosites from the same sample^33^, proximal proteins and phosphosites were detected using label-free quantitation to calculate enrichment relative to control samples (Supplementary Data 1, 2). 78 and 117 significant MSK-proximal proteins and phosphopeptides, respectively, were identified. These 117 phosphopeptides were found in 103 different proteins. Based on Gene Ontology (GO) functional and biological process annotations available on TriTrypDB, the MSK proximal proteome and phosphoproteome comprise hypothetical proteins (17 in the proximal proteome and 36 in the phosphoproteome), RNA processing factors (10 in each dataset), microtubule/cell motility (8 and 10 proteins in the proximal proteome and phosphoproteome, respectively), protein kinases (4 in each dataset), and nuclear pore components (2 and 3 proteins in the proximal proteome and phosphoproteome, respectively). Additionally, four proteins linked to post-transcriptional regulation and two phosphatases are uniquely enriched in the phosphoproteome. As expected, the bait protein MSK is enriched in both the total and phosphoproteome datasets, with two phosphopeptides containing phosphorylation sites at S609 and the S722*–S724* highlighted (Fig. 4a and Supplementary Data 1, 2).

**Fig. 4.**
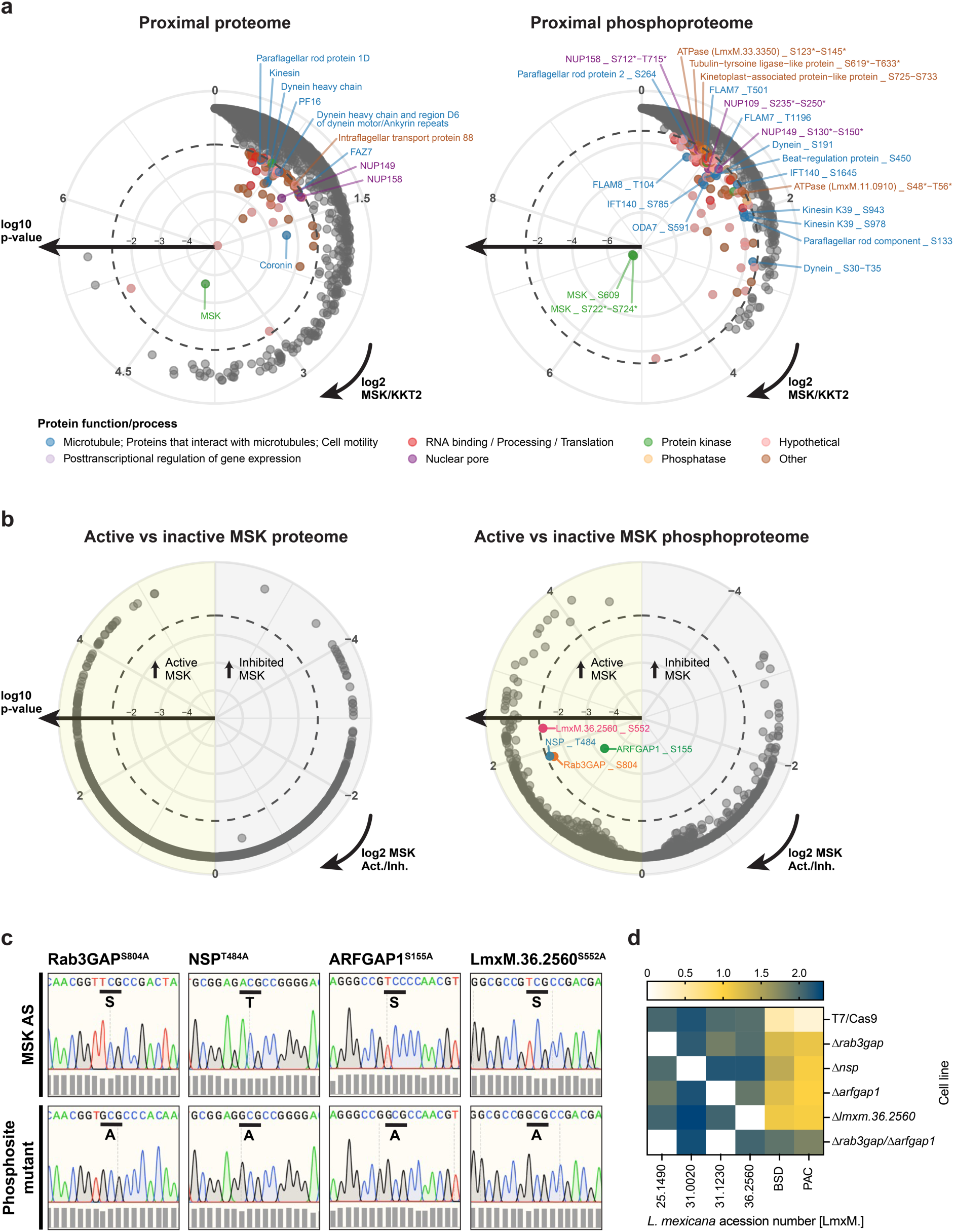
Mapping the proximal environment and substrates of MSK with combined chemical genetics and proximity labelling. **a** Radial plot of proteins (left panel) and phosphosites (right panel) enriched in MSK compared to the spatial reference KKT2, with phosphosites indicated. **b** Radial plot of protein (left panel) and phosphopeptides (right panel) enriched in MSK after inhibition with 1NM-PP1, with phosphosites indicated. The dashed line indicates a 5% false discovery rate (FDR) threshold for distinguishing enriched (coloured points) and non-enriched (grey points) total proteins and phosphosites. Log2 fold enrichment increases in a clockwise direction; Log10 p-value increases from the centre outward. *, where a phosphosite could not be confidently localised, the possible phosphorylation region is indicated. **c** Phosphosite mutants for MSK substrates. CRISPR-Cas9–mediated precision genome editing was used to mutate the target phosphosite to an alanine in the MSK AS line. Sanger sequencing of the phosphosite mutants and the wild-type region in the MSK AS line are shown. Sequencing files were visualised on SnapGene v.7.2 and the bars on the bottom indicate the sequence quality per base. Phosphosite mutant genotyping is shown in Supplementary Fig. 11. **d** Generation of *Rab3GAP*, *NSP*, *ARFGAP1* and *LmxM.36.2560* gene deletion mutants using CRISPR–Cas9 with two repair templates containing drug resistance selection markers (*BSD* and *PAC*). The heatmap indicates the copy number status of genes. Gene copy number was estimated by the ratio of the gene coverage by its chromosome coverage, multiplied by the chromosome ploidy.

Comparison of the MSK-enriched proteome with that obtained under MSK inhibition revealed no significant changes in the proximal proteome, suggesting that MSK inhibition does not disrupt MSK’s association with its binding partners (Fig 4b, left panel). Nevertheless, four phosphopeptides, each from a different protein, are significantly enriched in the proximal phosphoproteome of active MSK, highlighting them as likely substrates (Fig 4b, right panel). The corresponding phosphosites are S804 of Rab3GAP (Rab3 GTPase-activating protein catalytic subunit, LmxM.25.1490), T484 of NSP (nuclear segregation protein, LmxM.31.0020), S155 of ARFGAP1 (ADP-ribosylation factor GTPase activating protein 1, LmxM.31.1230), and S552 of a conserved hypothetical protein (LmxM.36.2560). Rab3GAP is not detected in the total proteome dataset, indicating that the phosphopeptide enrichment step increases the likelihood of phosphosite detection and identification. Furthermore, none of the other 3 proteins were assigned as MSK proximal in the proteome or in the phosphoproteome when KKT2 was used as the spatial reference control, suggesting a transient interaction between kinase and substrate (Supplementary Fig. 10).

To investigate the role of the four identified phosphosites, CRISPR-Cas9–mediated precision genome editing was used to engineer mutants with the serine or threonine phosphosites mutated to an alanine in both alleles of the targeted proteins (Fig 4c and Supplementary Fig.11). The success in generating these mutants shows that phosphorylation at these phosphosites is not essential for promastigote survival. To further explore the role of Rab3GAP, NSP, ARFGAP1 and LmxM.36.2560 we generated gene deletion mutants. Δ*rab3gap*, Δ*nsp*, Δ*arfgap1* and Δ*lmxm.36.2560* null mutants were isolated and confirmed by whole genome sequencing. This shows that these genes are not essential, but as the two GTPase-activating proteins (Rab3GAP and ARFGAP1) may have redundant roles, we attempted to delete both genes in the same cell line and created a Δ*rab3gap/arfgap1* double null mutant (Fig 4d and Supplementary Fig. 12). Together these results indicate a complex interaction network for MSK and its substrates with no single substrate or phosphosite of those identified in this study essential for promastigote cell viability.

We next investigated co-localization of MSK with its substrates. To this end, cell lines were engineered to simultaneously express 3xMyc::mNG::MSK_AS together with each substrate fused to a 3xHA tag at either the C- or N-terminus (Supplementary Fig. 13 – 14). Parasites were cultured in the presence or absence of 10 µM 1NM-PP1 for 6 h, and the distribution of MSK and its substrates during different stages of the promastigote cell cycle was visualised by fluorescent microscopy, with simultaneous labelling of β-tubulin, MSK, MSK substrate and DNA. Rab3GAP, NSP, ARFGAP1 and LmxM.36.2560 are constitutively expressed throughout the promastigote cell cycle. Moreover, MSK is found in close proximity to all four substrates within the nucleus, mitotic spindle, kinetoplast and cytoplasm. Beyond that, each substrate also showed proximity to β-tubulin (Fig. 5a and Supplementary Fig. 15).

**Fig. 5.**
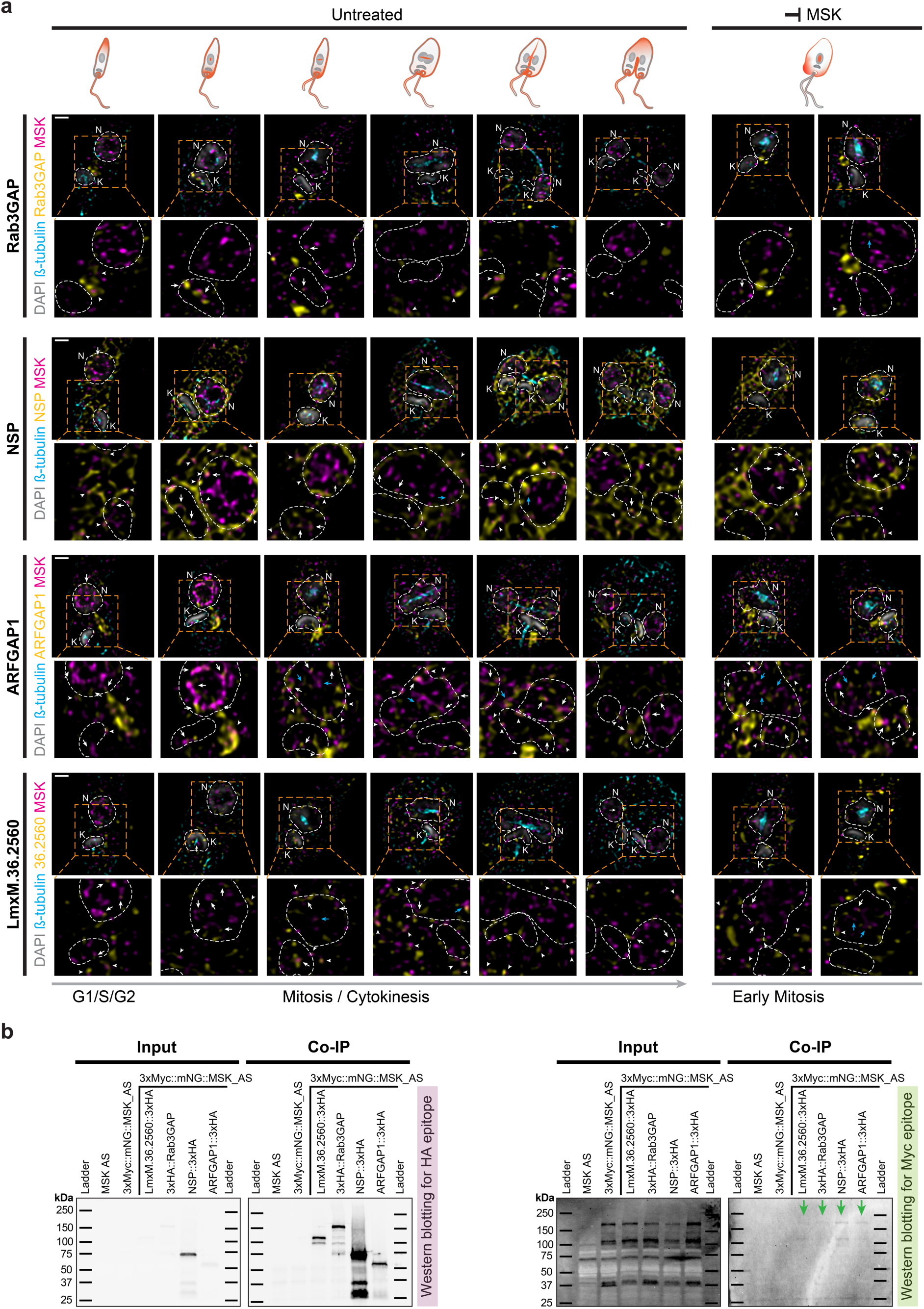
Rab3GAP, NSP, ARFGAP1 and LmxM.36.2650, revealed as new mitotic spindle components, interact with MSK. **a** Co-localization microscopy of MSK and its substrates (Rab3GAP, NSP, ARFGAP1 and LmxM.36.2650). High resolution fluorescent microscopy of promastigote parasites after 6 h of treatment with 10 µM 1NM-PP1 or untreated. Parasites were stained with KMX-1, anti-Myc, and anti-HA antibodies to, respectively, label β-tubulin, 3xMyc::mNG::MSK_AS, and the MSK substrates (Rab3GAP, NSP, ARFGAP1 or LmxM.36.2560) fused to 3xHA. Counterstaining with DAPI was used to visualise DNA. The colours used for each labelling are indicated on the left. Dashed line squares indicate areas magnified below each image. White arrows show MSK and its substrates co-localized in the nucleus or associated with the kinetoplast, while the blue arrows show co-localization of MSK and its substrates in the mitotic spindle. White arrowheads are, MSK and its substrate co-localized outside the nucleus/kinetoplast. Scale bars, 1 μm. Individual channels and other merged channels pattern are shown in the Supplementary Fig. 15. **b** Co-immunoprecipitation of MSK with its substrates. Epitope-tagged MSK substrate – 3xHA::Rab3GAP (138.82 kDa), NSP::3xHA (60.64 kDa), ARFGAP1::3xHA (47.09 kDa), and LmxM.36.2560::3xHA (104.96 kDa) – were used as bait in a crosslinking anti-HA co-immunoprecipitation. Enrichment of the bait proteins was confirmed by western blotting using an anti-HA antibody (left panel). Co-immunoprecipitation of the 3xMyc::mNG::MSK (predicted molecular weight: 171.24 kDa) protein was detected by western blotting using anti-Myc antibody (right panel). These blots represent a single experiment.

Although constitutively expressed, Rab3GAP is detected in the nucleus and kinetoplast only at specific stages of the cell cycle. Rab3GAP co-localises with MSK in the kinetoplast during mitochondrial DNA segregation and in the nucleus at late stages of DNA segregation – when nuclei are separated but the mitotic spindle is still present. At this stage, Rab3GAP also aligns with β-tubulin on the spindle, indicating it functions as a transient spindle component. Notably, inhibition of MSK kinase activity does not disrupt its co-localisation with Rab3GAP in either the nucleus or kinetoplast, suggesting that kinase activity is not required for correct localisation (Fig. 5a and Supplementary Fig. 15).

NSP is found throughout the promastigote and co-localises with MSK at multiple distinct sites. When β-tubulin moves to the nucleus to form the mitotic spindle, NSP is in proximity to MSK in the kinetoplast and in the nucleus, which was evident until the end of DNA segregation. However, like Rab3GAP, co-localization between NSP and β-tubulin in the mitotic spindle occurs only at the end of DNA segregation. Once DNA segregation is complete and the mitotic spindle is no longer present, points of co-localization between NSP and MSK occur only outside the nucleus and kinetoplast, more frequently close to the cytokinesis furrow. Notably, when the kinase activity of MSK was inhibited, NSP was found in proximity to MSK in the mitotic spindle. This interaction was not detected under physiological conditions, suggesting that it could be a transient interaction that is potentially important for elongation of the mitotic spindle (Fig. 5a and Supplementary Fig. 15).

ARFGAP1 is also distributed through the promastigote, but with multiple points of co-localization between MSK and ARFGAP1 detected close to the kinetoplast. In contrast to Rab3GAP and NSP, co-localization of MSK and ARFGAP1 in the nucleus and kinetoplast were sustained throughout the cell cycle. Furthermore, ARFGAP1 was detected in proximity to MSK and β-tubulin in the mitotic spindle from the beginning to the end of the DNA segregation process, indicating that ARFGAP1 is also a component of the mitotic spindle. In addition, co-localization of MSK and ARFGAP1 in the nucleus, kinetoplast, mitotic spindle, and cytoplasm was evident after treatment with 1NM-PP1 (Fig. 5a and Supplementary Fig. 15).

LmxM.36.2560 is found in the nucleus and kinetoplast in all promastigote cell cycle stages. Co-localization of LmxM.36.2560 protein and MSK in the nucleus was sustained throughout the cell cycle, however the two proteins were not found in proximity in the kinetoplast in the final steps of mitosis. Similarly to the other MSK substrates, LmxM.36.2560 co-localises with β-tubulin in the mitotic spindle, indicating that this protein is also a component of the spindle. In addition, co-localization of MSK and LmxM.36.2560 in the nucleus and mitotic spindle was evident after treatment with 1NM-PP1, but not in the kinetoplast (Fig. 5a and Supplementary Fig. 15).

The close association of MSK and Rab3GAP, NSP, ARFGAP1 and LmxM.36.2560 protein was confirmed by co-immunoprecipitation, using the epitope-tagged cell lines. Co-immunoprecipitation using anti-HA-coated beads successfully enriched all 3xHA-tagged MSK substrates. The small amount of MSK precipitated in these immunoprecipitations is consistent with the transient nature of MSK interactions found in our co-localization microscopy screening (Fig. 5b).

## Discussion

Segregation of the kinetoplast and nucleus in trypanosomatids is a tightly regulated process that ensures faithful inheritance of genetic material during cell division. Despite its importance, the kinase signalling pathways orchestrating this process remain poorly understood. Here we have taken an analogue-sensitive kinase approach to investigate the biological function of a previously uncharacterised kinase in *L. mexicana*, which is found in both the nucleus and the kinetoplast^24^. We called this kinase <u>m</u>itotic <u>s</u>pindle <u>k</u>inase, MSK, based on its predominant association with the mitotic spindle. Using precision genome editing, we successfully generated endogenous AS mutants of MSK and identified the glycine gatekeeper variant as responsive to inhibition by the BKI 1NM-PP1. Functional analysis of this mutant revealed that MSK activity is essential for parasite viability in both the promastigote and the clinically relevant intracellular amastigote stages, indicating that MSK is a promising candidate for the development of targeted antileishmanial kinase inhibitors.

In *L. mexicana* promastigotes, inhibition of MSK resulted in an unusual phenotype for trypanosomatids; cells with duplicated DNA arrested in early mitosis, characterised by unsegregated nuclei and kinetoplast, mitotic spindle blocked in its initial short ring-like conformation, and two fully elongated flagella (1K1N2F). These findings suggest that, in contrast to what has been described in *T. brucei*^36–38^, kinetoplast and nuclear segregation in *Leishmania* are not completely uncoupled, and that MSK activity co-ordinates *Leishmania* mitochondrial and nuclear DNA segregation. In *T. brucei*, kDNA segregation occurs during G2-phase, prior to nuclear division^39^. RNAi-mediated knockdown of the mitotic cyclin gene CYC6 in *T. brucei* provided the first genetic evidence of cell cycle dysregulation and revealed fundamental differences in cell cycle control between the parasite’s life cycle stages. In the procyclic form, CYC6 depletion resulted in the accumulation of anucleated zoid cells (1K0N), indicating that cytokinesis was not inhibited. In contrast, CYC6 knockdown in the bloodstream form inhibited both cytokinesis and nuclear segregation, while nuclear DNA synthesis, as well as kinetoplast replication and segregation, remained unaffected, leading to the accumulation of cells with abnormal DNA content (>2K1N)^40^. Besides, RNAi-mediated depletion of subunits of the chromosomal passenger complex (CPC) – including TbAUK1; TbKIN-A; TbKIN-B; TbCPC1; and TbCPC2 – in the procyclic form of *T. brucei* disrupted spindle assembly and nuclear segregation, while kinetoplast segregation remained unaffected. This led to an accumulation of cells with 2K1N configuration^41,42^. In the bloodstream form, depletion of TbAUK3 resulted in a significant reduction of 1K1N cells and a concomitant increase in multinucleated cells, zoids (1K0N), and aberrant cells lacking DNA (0K0N), or displaying other atypical configurations such as 0K1N and 1K2N^43^. Similarly, in *T. cruzi*, AUK1 is essential for symmetric mitosis, as its depletion led to the accumulation of 2K1N cells and zoids^44^. Conversely, disruption of kDNA segregation by knockdown of the casein kinase TbCK1.2 did not impact mitosis which progressed normally, resulting in cells with a single kinetoplast and two nuclei (1K2N)^45–47^. Interestingly, RNAi-mediated depletion of the MSK ortholog in *T. brucei* (Tb927.11.5340) resulted in a progressive decrease in 1K1N cells, accompanied by an increase in 2K1N cells, zoids (1K0N), and cells containing three or more kinetoplasts and one or two nuclei (≥3K1N or ≥3K2N)^45^.

We also observed that inhibition of *L. mexicana* MSK activity results in aberrant cytokinesis in mitotically arrested cells. In these parasites, cleavage initiates at the posterior end and progresses anteriorly, generating partially formed cells devoid of DNA-containing structures. These “cells” are distinct from previously described zoids, which originate from cells that have completed kinetoplast segregation but are compromised in mitosis, and which retain the new flagellum along with its associated basal body-kinetoplast complex^36^. Together, these observations suggest that *Leishmania* possesses alternative mechanisms for co-ordinating mitosis/cytokinesis and organelle segregation to those of *T. brucei*, with MSK activity likely contributing to this regulation.

Whilst we don’t have evidence that MSK inhibition disrupts the bipolar organization of the mitotic spindle or the attachment of microtubules to kinetochores, our data demonstrate that MSK inhibition impairs mitotic spindle elongation, consistent with a spindle assembly defect. This phenotype contrasts with that observed in *L. donovani* overexpressing the twinfilin-like protein (TWF), which exhibit abnormally elongated mitotic spindles, leading to chromosome mis-segregation^48^. The *L. donovani* heterozygous gene deletion mutant *TWF*^+/-^ exhibited a phenotype more closely resembling that observed upon MSK inhibition, characterized by the accumulation of cells with short mitotic spindles 6 to 10 h after release from hydroxyurea-induced synchronization. However, the increased proportion of cells with elongated spindles at 10 h post-release indicates that these cells eventually progress through mitosis, albeit with a delayed cell cycle compared to the wild-type *TWF^+/+^*line^48^. In addition, *TWF*^+/-^ mutants arrest in anaphase, displaying incomplete nuclear separation, despite kinetoplast segregation proceeding normally, indicating a mechanistic distinction from the MSK-inhibited phenotype. MSK and TWF also differ in their location during mitosis: MSK is enriched in the central region of the mitotic spindle, while TWF localizes at the ends of the spindle, where it is predicted to mediate microtubule elongation^48^. Taken together, these observations support a model in which MSK functions upstream of TWF in regulating mitotic spindle dynamics.

Previous studies have indicated that trypanosomes lack a canonical spindle assembly checkpoint (SAC), as they appear unable to arrest cell cycle progression in response to spindle defects^36–38^. However, we found that after 24 h of MSK inhibition, a timeframe encompassing approximately three cell cycles, 91.4% of the parasites had a DNA content less than or equal to 4n, indicating that cell cycle progression was either arrested or substantially delayed, possibly due to a checkpoint. Indeed, partial depletion of the TWF in *L. donovani*, which disrupts spindle elongation, was also associated with a delay in nuclear DNA synthesis^48^. In higher eukaryotes, the SAC serves as a surveillance system that delays anaphase onset until all chromosomes have achieved proper kinetochore-microtubule attachments and bi-orientation. Core SAC components include Mad1, Mad2, Mad3 (BubR1), Bub1, and Bub3^49^. Of these, only Mad2 has a recognized ortholog in trypanosomatids^50^, including *Leishmania*. In *T. brucei*, MAD2 localizes to the basal body rather than to kinetochores or the nucleus during mitosis, suggesting that in these flagellated eukaryotes, an unconventional checkpoint control could be linked to basal body state^37,51^. The presence of cells with two well-separated basal bodies but unsegregated kinetoplasts (1K2N) in *T. brucei*, however, indicates that basal body separation and kDNA segregation can be uncoupled^47^.

Intriguingly, we observed that during anaphase, MSK maintains continuous connection between the intranuclear mitotic spindle and the extranuclear subpellicular microtubules, possibly passing through a pore in the nuclear envelope. Consistent with this observation, our phosphoproteomic analysis identified the nuclear pore proteins NUP109, NUP149, and NUP158 as proximal to MSK, with NUP149 and NUP158 also found to be phosphorylated. These findings suggest a potential association between MSK and nuclear envelope substructures involved in microtubule organization. This organization may share features with the spindle pole body (SPB) – the functional analogue of the metazoan centrosome – found in the budding yeast *Saccharomyces cerevisiae*, which remains permanently embedded in specialized pores of the nuclear envelope throughout closed mitosis^52^. A different model has been proposed for the closely related kinetoplastid *T. brucei*, in which a ring-like structure located entirely within the nucleus and adjacent to the nuclear envelope, appears to function as the microtubule organizing centre (MTOC) during mitosis^28^. Moreover, trypanosomatids lack orthologs of Brr6, a nuclear envelope protein required for SPB insertion during mitosis in *Schizosaccharomyces pombe*^53^, supporting the idea that fenestration-dependent SPB insertion may not occur in these organisms. Supporting a role for MSK in spindle microtubule dynamics, and potentially in microtubule nucleation, in *Leishmania*, our proximity phosphoproteomic analysis also identified 18 proteins associated with the Gene Ontology category “microtubule/cell motility”. Of the known SPB components in *S. cerevisiae*, only the tubulins Tub4, Spc97, and Spc98, the calmodulin CMD1, and the centrin Cdc31 have identifiable orthologs in *Leishmania*. Our phosphoproteomic analysis identified the *L. mexicana* ortholog of Cdc31 (LmxM.01.0620) as being proximal to MSK. In yeast, Cdc31 has been characterized as a key regulator of SPB duplication^54^. In *L. donovani*, deletion of the centrin gene resulted in defective basal body duplication and cytokinesis, although this phenotype was restricted to the axenic amastigote stage^55^. Despite these insights, canonical spindle pole-specific components have not yet been identified in trypanosomatids, and the precise mechanisms underlying bipolar spindle assembly in these parasites remain largely unknown.

By using the analog-sensitive kinase approach combined with proximity phosphoproteomics, we identified four MSK substrates: Rab3GAP, ARFGAP1, NSP, and the hypothetical protein LmxM.36.2560. All phosphorylation sites attributed to MSK exhibited a proline residue immediately adjacent to the phosphoacceptor site, which is a common feature of substrate recognition motifs by CMGC family kinases. Our data suggest that these proteins interact transiently with MSK in a manner that appears to be independent of its kinase activity. All four proteins were found to be constitutively expressed during the cell cycle and to co-localize with MSK in multiple subcellular compartments, including the nucleus, kinetoplast, cytoplasm, and the mitotic spindle. These proteins were also found near β-tubulin on the mitotic spindle, supporting their identification as previously uncharacterized spindle-associated proteins (SAP). While many SAPs identified in yeast and animals function as microtubule-associated proteins^56^, InterPro domain analysis revealed that none of the SAPs identified in this study possess canonical microtubule-binding motifs (data not shown). Despite the central roles of Rab and Arf GTPases in mitosis and cytokinesis in other eukaryotes^57,58^, our functional analyses in *L. mexicana* showed that individual and combined Rab3GAP and ARFGAP1 gene deletions did not result in any obvious growth or morphological defects. Similarly, strains harbouring mutations in their predicted MSK phosphorylation sites did not exhibit alterations in their phenotype. This lack of phenotype may be explained by functional redundancy among GAP proteins^59^. Moreover, GAP proteins can act on multiple GTPase targets, adding to the complexity of their functional roles^60^. OrthoMCL analysis of ARFGAP1 identified five ArfGAP-domain-containing proteins in *L. mexicana*, suggesting redundancy. In contrast, no additional Rab3GAP orthologs were identified in *L. mexicana*, and comparative analysis indicates that this protein is conserved only within the *Trypanosomatida* lineage.

Similar to Rab3GAP, NSP lacks additional orthologs in *L. mexicana* and appears to be conserved in the *Trypanosomatida*. InterPro protein sequence analysis predicts that NSP belongs to the nuclear segregation protein Bfr1 family, originally described in *S. cerevisiae*. Bfr1 is a non-canonical RNA-binding protein localized to the cytosolic face of the endoplasmic reticulum (ER) membrane, where it plays a role in ER-Golgi transport and in directing mRNAs encoding secretory pathway proteins to the ER for processing^61,62^. Moreover, Bfr1 has been implicated in nuclear segregation and has been shown to interact with Bbp1, a protein involved in nuclear spindle assembly^63,64^. Notably, Bfr1 deletion in yeast resulted in a phenotype reminiscent of MSK inhibition in *Leishmania*, characterized by large-budded cells with a single nucleus and a short mitotic spindle^63^. The NSP ortholog in *T. brucei* (ERBP1, Tb927.10.14150) has also been characterized as a non-canonical RNA-binding protein that localizes to the ER and associates with mRNAs encoding ribosomal proteins. Consistent with findings in *L. mexicana*, ERBP1 is not essential in bloodstream-form *T. brucei*^65^. None of the MSK substrates identified in this study have been previously linked to cell cycle regulation in trypanosomatids.

Similar to other MSK substrates like Rab3GAP and ARFGAP1, the hypothetical protein LmxM.36.2560 showed structural homology to importin-9 (Supplementary Fig. 16). Importin-9 is a protein known to interact with small GTP-binding proteins^66^, and is implicated in chromosome segregation in *Drosophila*^67^.

Overall, our study revealed MSK as an essential protein kinase required for mitotic spindle elongation and the co-ordinated segregation of the nucleus and kinetoplast in *Leishmania*. We identified four MSK substrates and characterized them as previously unrecognized mitotic spindle-associated proteins. These findings provide new insights into the unique mechanisms of cell division in trypanosomatids.

## Methods

### Cell culture

The promastigote stage of *Leishmania mexicana* (MNYC/BZ/62/M379 T7/Cas9)^68,69^ were maintained in culture at 25 °C in HOMEM medium (modified Eagle’s medium – Gibco, ThermoFisher Scientific) supplemented with 10% heat-inactivated fetal calf serum (hi-FCS) (Gibco, ThermoFisher Scientific) and 100 U penicillin – 100 μg mL^-1^ streptomycin (Sigma-Aldrich), pH 7.2. Where required, parasites were grown with selective antibiotics at the following concentrations: Hygromycin B (InvivoGen, ant-hg) at 50 μg mL^-1^; Nourseothricin (Jena Bioscience, AB-101) at 50 µg mL^-1^; Blasticidin (InvivoGen, ant-bl) at 10 μg mL^-1^; Puromycin (InvivoGen, ant-pr) at 30 μg mL^-1^. Bone marrow-derived macrophages (BMDM) isolated from BALB/c mice were differentiated in DMEM medium supplemented with 10% hi-FCS and macrophage colony-stimulating factor secreted by L929 cells^70^. BMDM was maintained in culture in DMEM medium supplemented with 10% hi-FBS and 10 mM L-glutamine (Gibco, ThermoFisher Scientific) at 37°C, in an atmosphere of 5% CO_2_.

### Growth curve

The promastigote form of *L. mexicana* was set up at 4 × 10^4^ parasites mL^-1^ in HOMEM medium supplemented with 10% hi-FBS, and the cumulative cell growth was measured daily by cell counting in a Neubauer chamber. The growth rate was calculated in the logarithmic area of the growth curve (0 – 96 h).

### Structural prediction of the MSK kinase domain and identification of the gatekeeper residue

The MSK kinase domain was identified through protein domain analysis using the InterPro 103.0^29^ tool. The amino acid sequence corresponding to the kinase domain was used to generate a structural model of the MSK kinase domain bound to ATP, predicted with AlphaFold 3^30^ and visualized using ChimeraX v1.9. The gatekeeper residue, located in the hydrophobic region of the ATP-binding pocket, was identified through sequence alignment and manually inspected within the predicted structural model.

### *L. mexicana* genome editing by CRISPR-Cas9

Genome editing was performed in *L. mexicana* strain MNYC/BZ/62/M379 constitutively expressing Cas9 and T7 RNA polymerase, as previously described^68,69^. To introduce single-codon mutations without selection markers we used our previous adapted CRISPR-Cas9-mediated precise genome editing strategy^27^. Briefly, two sgRNAs targeting sequences flanking the codon of interest were designed using the Eukaryotic Pathogen CRISPR guide RNA/DNA Design Tool (http://grna.ctegd.uga.edu). sgRNA templates were generated using primers adapted from Beneke et al.^69^. Single-stranded 120-nt DNA repair templates (ssDRT) were synthesised to include the desired mutation, silent shield mutations to prevent Cas9 re-cutting, and 23 – 25 nt homology arms flanking the target site.

For phosphosite mutagenesis, 200 bp double-stranded DRT (dsDRT) were generated by PCR using oligonucleotides with 18 – 20 bp complementary overhangs. These templates were designed to encode the intended codon change, silent shield mutations, and 48 – 72 bp homology arms flanking the Cas9-induced double-strand break. PCR products were purified using the QIAquick PCR Purification Kit (Qiagen) and eluted in nuclease-free water at ∼1µg µL^-1^. For transfection, ∼5 µg of sgRNA and either 11 µg of ssDRT or ∼5 µg of dsDRT were co-transfected into 5 × 10⁶ promastigotes using the P3 Primary Cell 4D-Nucleofector Kit (Lonza) with program FI-115. After electroporation, cells were transferred to pre-warmed HOMEM medium with 20% hi-FCS and 10 µM 6-biopterin. After a 16-hour recovery period, transfected cells were cloned into 96-well plates at a ratio of one cell per two wells. Gene knockouts and endogenous tags were generated using the CRISPR-Cas9 toolkit for kinetoplastids^69^. sgRNA and repair template primers were designed using the LeishGEdit platform (http://leishgedit.net/). For knockout mutants, two repair templates carrying different drug resistance markers were used. Parasites were co-transfected with purified sgRNA and repair templates as described above. Sixteen hours post-transfection, selection with blasticidin and/or puromycin was applied, and clones were obtained by limiting dilution. Genomic DNA from recovered clones was extracted for diagnostic PCR and, where necessary, followed by restriction enzyme digestion. Digestions were incubated for 16 h using the appropriate restriction enzymes and buffers recommended by New England Biolabs. Oligonucleotide sequences used to generate and validate CRISPR-Cas9-edited lines are provided in Supplementary Tables 1 – 3.

Null knockout mutants were validated by Illumina paired-end sequencing (150 bp reads). Reads were aligned to the *L. mexicana* MNYC/BZ/62/M379 Cas9/T7 reference genome^68^ using Minimap2 v2.26-r1175. Alignments were visualized in IGV v2.16.1. Genome coverage was assessed using Mosdepth v0.3.3, which calculated read depth in 500 bp windows and mean coverage across all chromosomes. Gene coverage was also determined by Mosdepth based on annotations from the MNYC/BZ/62/M379 Cas9/T7 reference. Sample coverage was normalised per chromosome, and the coverage change was calculated from the relative difference between the comparison and reference. 0.5 was used as a cut off, which equates to one copy change. The Illumina sequencing data were deposited under the Bioproject accession numbers PRJNA1311012 and PRJNA1303394.

### *In vitro* susceptibility assay of *Leishmania* promastigotes

A dose response curve was set in a 96-well plate with 1x10^4^ parasites mL^-1^ treated with two-fold increasing concentrations of the inhibitor (1NA-PP1 or 1NM-PP1). The viability of treated and untreated control was assessed after 96 h by addition of 50 µL of 0.0125% (w/v) resazurin (Alamar Blue) prepared in PBS. Cells were incubated for an additional 2 – 4 h at 37°C. Fluorescence emission of the reduced resazurin was detected using a CLARIOstar^®^ reader (BMG LABTECH; excitation filter at 540 nm and emissions filter at 590 nm). Fitting of dose-response curves and IC_50_ calculation were carried out using GraphPad Prism v10.4.1, considering the untreated control for each cell line as 100% viability.

### *In vitro* susceptibility assay of intracellular *Leishmania* amastigotes

Promastigotes were grown in HOMEM medium supplemented with 10% hi-FCS until reaching late-log phase. Differentiated BMDM were plated on 16-well Labtek tissue culture slides (Nunc, NY, USA) and then infected with promastigote-form *Leishmania* at ratio of 10 parasites per macrophage. After 18 h incubation in DMEM supplemented with 5% hi-FCS in 5% CO_2_ at 37 °C, free promastigotes were removed and the cultures were then exposed to different concentrations of the inhibitors (1NA-PP1 or 1NM-PP1) prepared in DMEM supplemented with 2% heat-inactivated horse serum. After 96 h additional incubation, the slides were fixed with methanol and then stained with Giemsa. The slides were examined using Zeiss Axiolab-5 microscope and 100 cells in each well were counted to determine the percentage of infected macrophages. The parasite viability was determined from the percentage of infected cells in treated cultures relative to untreated controls. Fitting of dose-response curves and IC_50_ calculation were carried out using GraphPad Prism v10.4.1, considering the untreated control for each cell line as 100% viability.

### Cell cycle analysis

Promastigote cells growing in the presence or absence of the inhibitor (10 µM 1NM-PP1) for 6 or 24 h were washed in PBS-EDTA (PBS supplemented with 5 mM of EDTA) and resuspended in 70% methanol. After overnight incubation at 4°C, cells were washed once with PBS-EDTA and then resuspended in 1ml PBS-EDTA containing 10 µg mL^-1^ propidium iodide and 10 µg mL^-1^ RNase A. Cells were incubated for 45 min at 37°C in the dark and then analysed for FACS using a CyAn^TM^ ADP cytometer (Beckman Coulter). FlowJo^TM^ 10.6.2 cell cycle algorithm Watson model was used to determine cell cycle distribution.

### Immunofluorescence Microscopy

Promastigote cells growing in the presence or absence of the inhibitor (10 µM 1NM-PP1) for 6 or 24 h were washed twice with PBS (1,400 g for 10 min at room temperature). About 10^6^ cells in PBS were left to adhere on a poly-L-lysine-treated high precision coverslip (thickness No. 1.5H [0.170 mm ± 0.005 mm], MARIENFELD: cat. 0107222) for 15 min at 37°C. *In vivo* cross-linking was then performed incubating the adhered cell with 1 mM disuccinimidyl suberate (DSS) in PBS for 10 min at 37°C. Attached parasites were fixed at room temperature with 4% paraformaldehyde in PBS for 15 min, followed by quenching with 0.1 M glycine in PBS (pH 7.6) for 5 min. After two washes with PBS, cells were permeabilized with 0.5% Triton X-100 in PBS for 15 min. Blocking was performed by incubating cells in blocking buffer (5% BSA, 0.01% saponin in PBS) for 1 h at room temperature. Primary immunostaining was carried out for 1 h at room temperature using mouse anti-ϕ3-tubulin KMX-1 antibody (Sigma-Aldrich, MAB3408) diluted 1:1000 in blocking buffer. After three washes with 0.1% Triton X-100 in PBS, cells were incubated for 1 h with Alexa Fluor^TM^ 647-conjugated goat anti-mouse IgG secondary antibody (Abcam ab150119) diluted 1:1000 in blocking buffer. Following three washes with 0.1% Triton X-100/PBS, cells were counterstained with 20 µg mL^-1^ DAPI in PBS for 30 min, followed by a final PBS wash. Coverslips were mounted on glass slides using ProLong diamond antifade mountant (Invitrogen™), according to the manufacturer’s instructions. For simultaneous detection of endogenously tagged MSK and its substrates, the above protocol was followed using the following antibody combinations: mouse anti-ϕ3-tubulin KMX-1 [diluted 1:800], rabbit anti-Myc-Tag (clone 71D10, Cell Signaling Technology mAb #2278) [diluted 1:200], and rat anti-HA-Tag (clone 7C9, chromotek 7c9-100) [diluted 1:200] as primary antibodies; and Alexa Fluor™ 568-conjugated goat anti-mouse IgG (Invitrogen A-11031) [diluted 1:800], Alexa Fluor™ 488-conjugated donkey anti-rabbit IgG (Invitrogen A-21206) [diluted 1:200], and Alexa Fluor^TM^ 647-conjugated donkey anti-rat IgG (Invitrogen A-78947) as secondary antibodies.

### Microscopy and Image Analysis

Cells were examined by widefield fluorescence microscopy using a Zeiss Axio Observer7 microscope in z-stack mode (20 optical sections were captured per field). Images were processed using Fiji v.2.14.0/1.54f, employing the Microvolution blind deconvolution module. Maximum intensity projections were subsequently generated using selected z-stack layers retrieved with the “z-project” function in Fiji. Super-resolution structured illumination microscopy (SR-SIM) was performed on a Zeiss Elyra 7 system using the Lattice SIM modality. 3D acquisition in 4 colour was performed [DAPI Excitation 405nm Emission 420-480nm; AF488 Excitation 488nm Emission 490-560nm; AF568 Excitation 561nm Emission 570-630nm; AF647 Excitation 642nm Emission >640nm] with a z-stack of 50 slices captured at 0.091 µm intervals. Zeiss Zen Black v3.0 software was used to reconstruct the images by SIM^2^ processing using different pre-set processing parameter depending on the fluorescent signal intensities [fixed standard for β-tubulin_AF568 and NSP_AF647; fixed weak for MSK_AF488, Rab3GAP_AF647, RFGAP1_AF647, and LmxM.36.2560_AF647; and low contrast for DAPI]. To ensure accurate colour alignment, SR-SIM images were taken of TetraSpeck fluorescent microspheres with 200 nm diameter (Thermo Fisher Scientific). Then the Zen Black Channel Alignment module was used to calculate a correction matrix that was applied to the experimental images.

### Electron microscopy

Promastigote cells growing in the presence or absence of 10 µM 1NM-PP1 for 24 h were washed twice with PBS (1,400 g for 10 min at room temperature). Cells were then fixed for 1 h in 2.5% type II glutaraldehyde (Sigma, Missouri, USA) diluted in 0.1 M cacodylate buffer pH 7.2. For scanning electron microscopy (SEM), cells were subsequently adhered to coverslips precoated with 1 mg mL^-1^ poly-L-lysine, postfixed for 1 h with 1% osmium tetroxide diluted in cacodylate buffer, and dehydrated in a graded alcohol series (50%, 70%, 90%, and two exchanges of 100% ethanol for 10 min each step). Samples were critical-point dried in a Leica EM CPD030 apparatus (Leica, Wetzlar, Germany). Specimens were sputtered with gold in a Balzers FL9496 unit (Postfach 1000 FL-9496 Balzers Liechtenstein) and observed in an EVO 40 VP SEM (Zeiss, Germany). For transmission electron microscopy (TEM), cells were washed twice in cacodylate buffer and postfixed during 1h with 1% osmium tetroxide, 0.8% potassium ferrocyanide, 5 mM calcium chloride diluted in 0.1 M cacodylate buffer. Samples were then washed in cacodylate buffer, dehydrated in a graded series of acetone solutions (50%, 70%, 90%, and two exchanges of 100% acetone) for 10 min at each step, and embedded in Polybed resin. Ultrathin sections were stained with 5% uranyl acetate for 45 min and lead citrate for 5 min before observation in a Tecnai™ Spirit TEM transmission electron microscope (Zeiss, Oberkochen, Germany).

### Proximity Labelling

Proximity labelling was performed using *L. mexicana* cell lines engineered via CRISPR-Cas9 to express MSK_AS and KKT2_AS variants fused to 3xMyc epitope and the miniTurbo biotin ligase (3xMyc::mT::AS_MSK and KKT2_AS::mT::3xMyc) (Supplementary Fig. 9). Parasites were cultured to 5×10⁶ cells mL^-1^ and treated for 2 h at 25 °C with 10 µM 1NM-PP1 and 0.5 mM biotin to simultaneously inhibit kinase activity and induce biotinylation. Untreated cells served as controls. Each condition was performed in five biological replicates, following our previously described methodology^33^. After treatment, parasites were washed twice with PBS (15 min, 1,500 g) and resuspended to 7×10⁷ cells mL^-1^ in pre-warmed PBS. DSP crosslinker was added to 1 mM and *in vivo* cross-linking proceeded for 10 min at 37°C. After cross-linking, the reaction was quenched with 20 mM Tris-HCl (pH 7.5) for 5 min, parasites were harvested by 10 min centrifugation at 1,500 g and pellets were stored at -80°C until lysis. For each sample, 7×10^8^ cells were lysed in 500 μL ice-cold RIPA buffer (0.1% sodium dodecyl sulfate, 0.5% sodium deoxycholate, 1% IgePal-CA-630, 0.1 mM EDTA, 125 mM NaCl, 50 mM Tris pH 7.5) containing 0.1 mM PMSF, 1 μg mL^−1^ pepstatin A, 1 μM E-64, and 0.4 mM 1-10 phenanthroline. In addition, every 10 ml of RIPA was supplemented with 200 μL proteolytic protease inhibitor cocktail (Abcam, ab270055: w/v 2.16% AEBSF, 0.047% aprotinin, 0.156% bestatin, 0.049% E-64, 0.084% leupeptin, 0.093% pepstatin A), four tablets of complete protease inhibitor EDTA free (Roche, 05892791001) and one tablet PhosSTOP (Roche, 4906845001). Lysates were sonicated with a microtip sonicator on ice for three rounds of 10 s each at 25% amplitude. Digestion of nucleic acids for 50 mins on ice was achieved by adding 250 units of BaseMuncher endonuclease (Abcam, ab270049) and the lysates were then clarified by centrifugation at 10,000 g for 10 min at 4 °C.

For enrichment of biotinylated material, 1 mg of MagResyn^®^ Streptavidin beads (Resyn Bioscience, MR-STV005) was used for each affinity purification, which was carried out by end-over-end rotation at 4°C overnight. Beads were sequentially washed for 5 min in: RIPA buffer supplemented with protease and phosphatase inhibitors for four washes; 4 M urea for one wash; 6M urea for one wash; 1M KCl for one wash; and 50 mM TEAB pH 8.5 for one wash. Beads from each affinity purification were then resuspended in 200 μL 50 mM TEAB pH 8.5 containing 0.01% ProteaseMAX (Promega, V2071), 10 mM TCEP, 10 mM Iodoacetamide, 1 mM CaCl_2_ and 500 ng Trypsin Lys-C (Promega, V5071). The on-bead digest was carried out overnight at 37 °C while shaking at 800 rpm. Supernatant from digests was retained and beads were washed for 5 min in 50 μL water which was then added to the supernatant. Digests were acidified with trifluoroacetic acid (TFA) to a final concentration of 0.5% before centrifugation for 10 min at 17,000 g. The supernatant was desalted using in-house prepared C_18_ desalting tips, and eluted in 60 µL 80% acetonitrile with 0.1% TFA. 40% of the desalted peptides volume were dried down for MS analysis of the ‘Total’ proximal protein, whereas the remaining 60% were used to enrich for proximal phosphopeptides.

For proximal phosphopeptide enrichment to the remaining 36 µL of the desalted peptides 51.2 μL acetonitrile, 10 μL 1 M glycolic acid and 5 μL TFA were added. For each affinity purification 200 micrograms MagReSyn Ti-iMAC-HP beads (ReSyn Biosciences, MR-THP002), resuspended in 500 µL loading buffer (0.1 M glycolic acid, 80% acetonitrile (ACN), 5% TFA), were used. Peptides were added to the beads and incubated with shaking at 800 rpm for 40 min at room temperature. Beads were then washed for 2 min at 800 rpm in the following buffers: 100 μL loading buffer; 100 μL 80% ACN, 1% TFA; and 100 μL 10% CAN, 0.2% TFA. Phosphopeptides were eluted in 2 × 40 μL 1% NH_4_OH for 10 min shaking at 800 rpm. 4 μL TFA were added and phosphopeptides were dried down for MS analysis.

### Mass spectrometry data acquisition

For ‘Total’ proximal proteome analysis, peptides were loaded onto an mClass nanoflow UPLC system (Waters) equipped with a nanoEaze M/Z Symmetry 100 Å, C_18_, 5 μm trap column (180 μm × 20 mm, Waters) and a PepMap, 2 μm, 100 Å, C_18_ EasyNano nanocapillary column (75mm× 500 mm, Thermo). Separation used gradient elution of two solvents: solvent A, aqueous 0.1% (v:v) formic acid; solvent B, ACN containing 0.1% (v:v) formic acid. The linear multi-step gradient profile for ‘Total’ protein was: 3 – 10% B over 8 min, 10 – 35% B over 115 min, 35 – 99% B over 30 min and then proceeded to wash with 99% solvent B for 4 min. For phosphopeptide enriched samples, the following gradient profile was used: 3 – 10% B over 7 min, 10 – 35% B over 30 min, 35 – 99% B over 5 min and then proceeded to wash with 99% solvent B for 4 min. In all cases, the trap wash solvent was aqueous 0.05% (v:v) trifluoroacetic acid and the trapping flow rate was 15 μL min^-1^. The trap was washed for 5 min before switching flow to the capillary column. The flow rate for the capillary column was 300 nL min^-1^ and the column temperature was 40 °C. The column was returned to initial conditions and re-equilibrated for 15 min before subsequent injections.

The nanoLC system was interfaced with an Orbitrap Fusion hybrid mass spectrometer (Thermo) with an EasyNano ionisation source (Thermo). Positive ESI-MS and MS^2^ spectra were acquired using Xcalibur software (version 4.0, Thermo). Instrument source settings were: ion spray voltage, 1,900 V; sweep gas, 0 Arb; ion transfer tube temperature, 275 °C. MS spectra were acquired in the Orbitrap with 120,000 resolution, scan range: *m/z* 375 – 1500; AGC target, 4e5; max fill time, 100 ms. Data-dependent acquisition was performed in top speed mode using a fixed 1 s cycle, selecting the most intense precursors with charge states 2 – 5. Easy-IC was used for internal calibration. Dynamic exclusion was performed for 50 s post precursor selection and a minimum threshold for fragmentation was set at 5e3. MS^2^ spectra were acquired in the linear ion trap with: scan rate, turbo; quadrupole isolation, 1.6 *m/z*; activation type, HCD; activation energy: 32%; AGC target, 5e3; first mass, 110 m/z; max fill time, 100 ms. Acquisitions were arranged by Xcalibur to inject ions for all available parallelizable time.

### Mass spectrometry data analysis

Peak lists in .raw format were imported into Progenesis QI (Version 2.2., Waters) for peak picking and chromatographic alignment. A concatenated product ion peak list was exported in .mgf format for database searching against the *Leishmania mexicana* subset of the TriTrypDB (8,250 sequences; 5,180,224 residues) database, appended with common proteomic contaminants. Mascot Daemon (version 2.6.1, Matrix Science) was used to submit searches to a locally-running copy of the Mascot programme (Matrix Science Ltd., version 2.7.0.1). Search criteria specified: Enzyme, trypsin; Max missed cleavages, 2; Fixed modifications, Carbamidomethyl (C); Variable modifications, Oxidation (M), Phospho (STY), Acetyl (Protein N-term, K), Biotin (Protein N-term, K); Peptide tolerance, 3 ppm (# 13 C = 1); MS/MS tolerance, 0.5 Da; Instrument, ESI-TRAP. Peptide identifications were passed through the percolator algorithm to achieve a 1% false discovery rate as assessed empirically by reverse database search, and individual matches were filtered to require minimum expected scores of 0.05. The Mascot.XML results file was imported into Progenesis QI, and peptide identifications associated with precursor peak areas were mapped between acquisitions. Relative protein abundances were calculated using precursor ion areas from nonconflicting unique peptides. For ‘Total’ proteome data, only non-modified peptides were used for protein-level quantification. Normalisation was performed in Progenesis QI^71^. For phosphopeptide identifications, Mascot-derived site localisation probabilities were used. To associate quantification data with site localisation probabilities, Mascot search results, including raw spectral details and QI quantification data were exported separately in .csv format. Empty rows in the Mascot.csv were removed in R (3.6.1) before joining the data from the two .csv files using KNIME Analytics Platform (4.3.1 KNIME AG). The combined.csv was then stripped of non-quantified peptides.

For ‘Total’ proximal data, peptide ion quantification data were exported from Progenesis LFQ and proteins with ≥2 missing values across the five biological replicates were removed. Missing value imputation was performed for each sample group, drawing values from a left-shifted normal log2 intensity distribution to model low abundance proteins (mean = 6.5; sd = 1.5). For each protein, a mean intensity profile across all samples was calculated from peptide ion intensities. For each peptide ion, a Pearson correlation coefficient was calculated between the mean protein intensity profile and the individual peptide ion intensity profile. Peptide ions with a correlation coefficient >0.4 were then summed to calculate the label-free intensity of the parent protein. One peptide was required for protein quantification and one phosphopeptide for phosphosite quantification. Protein intensities were log2 transformed and proximal proteins were determined with the limma package^72^ using options trend = TRUE and robust = TRUE for the eBayes function. Protein intensities in the inhibited MSK samples were compared to those in the untreated MSK or KKT2 control sample to determine proximal proteins. Multiple testing correction was carried out according to Benjamini & Hochberg, the false discovery rate for identified proximals was 5%.

Label-free intensities for phosphosite proximal data were exported from Progenesis LFQ and phosphopeptides with ≥2 missing values across the 5 biological replicates were removed. Missing values were imputed by drawing values from a left-shifted normal log2 intensity distribution to model low abundance phosphopeptides (mean = 3.4; sd = 1.1). Phosphosites were aggregated by summing intensities which were then log2 transformed. Proximal phosphosites were determined with the limma package^72^ using options trend = TRUE and robust = TRUE for the eBayes function. Multiple testing correction was carried out according to Benjamini & Hochberg, false discovery rate for proximal phosphosites was 5%.

### Co-immunoprecipitation

MSK AS cell lines were engineered to express both candidate substrate proteins and MSK simultaneously tagged at the endogenous loci with 3xHA_puromycin and 3xMyc::mNG_blasticidin, respectively. For co-immunoprecipitation, 6.8x10^8^ mid-log phase promastigotes were harvested by centrifugation (1,200 g, 10 min, room temperature), washed twice in PBS, and resuspended in 10 mL pre-warmed PBS containing 1 mM DSP crosslinker. Crosslinking was performed for 10 min at 37°C, then quenched by addition of Tris-HCl (pH 7.5) to a final concentration of 20 mM for 5 min. Parasites were then harvested by 10 min centrifugation at 1,200 g and pellets were stored at -80°C until lysis. Cell pellets were lysed using 1 mL ice-cold RIPA buffer (Thermo, Cat. 89900) supplemented with 3x HALT Protease inhibitors (Thermo, Cat. 78430), PhosSTOP (Roche, Cat. 4906837001) and 10 µM E-64. After 30 minutes incubation on ice, samples were subjected to 3 cycles of sonication (30 s on, 30 s off) using a Bioruptor Pico (Diagenode). Nucleic acids were digested by adding 500 U BaseMuncher endonuclease (Abcam, ab270049) and incubating for 30 min on ice followed by 5 min at 37°C. Lysates were clarified by centrifugation (17,000 g, 15 min, 4 °C). HA-tagged proteins were immunoprecipitated with 40 µL anti-HA magnetic beads (Pierce, Cat. 88836) for 3 h at 4°C with end-over-end rotation. Beads were washed five times with ice-cold lysis buffer and bound proteins were eluted in 60 µL 1.5x Laemmli buffer containing 100 mM DTT, heated at 70°C for 15 min. Eluted proteins were fractionated by SDS-PAGE using 4 – 15% TGX Stain-Free™ Protein Gels (Biorad, Cat. 4568083). Western blotting was used to detect co-precipitated 3xMyc::mNG::MSK and HA-tagged bait proteins. Membranes were first probed with chicken anti-Myc tag (Bethil Laboratories, Cat. A190-103A) and DyLight^TM^800-conjugated anti-chicken (Invitrogen, Cat. SA5-10076) antibodies diluted at 1:4,000. After imaging, HA-tagged bait proteins were detected using rat anti-HA tag (chromotek, Cat. 7c9-100) and Alexa Fluor^TM^ 647-conjugated anti-rat (Invitrogen, Cat. A78947) antibodies diluted at 1:4,000.

### Statistics

Unless otherwise indicated, data were obtained from three independent biological replicates. Statistical analyses were performed using GraphPad Prism v10.4.1, with the specific tests detailed in the corresponding figure legends. Results are presented as mean ± standard error of the mean (SEM). Mass spectrometry data were analysed in RStudio v.2023.09.1+494 using the limma package^72^. Each condition included five independent replicates, except for the KKT2 total proteome dataset, which included four replicates.

### Reporting summary

Further information on research design is available in the Nature Research Reporting Summary linked to this article.

## Acknowledgements

This work was supported by funding from MRC GCRF (MR/P027989/1) and Wellcome Trust (223045/Z/21/Z). The funders had no role in study design, data collection, data analysis, interpretation, and writing of the manuscript. We thank our colleagues in the Bioscience Technology Facility of University of York who provided insight and expertise that greatly assisted our microscopy, flow cytometry, genomic data analysis, and mass spectrometry research. The York Centre of Excellence in Mass Spectrometry was created thanks to a major capital investment through Science City York, supported by Yorkshire Forward with funds from the Northern Way Initiative, and subsequent support from EPSRC (EP/K039660/1; EP/M028127/1).

## Author contributions

J.B.T.C., J.C.M. and A.J.W. conceived the project. J.C.M. and A.J.W. supervised the project. J.B.C.T., V.G., J.A.B., M.S., A.P.C.A.L. A.D. and M.C.M.M designed the experiments. J.B.T.C., V.G., J.A.B., M.C.M.M. A.D. and C.H. performed the experiments. J.B.T.C., V.G., J.A.B., M.S., A.P.C.A.L. A.D. and M.C.M.M. analysed experimental data. J.C.M. acquired funding. J.B.C.T and J.C.M. wrote the manuscript and all other authors revised and approved it.

## Additional information

Supplementary information: Supplementary Tables and Figures supporting the main text.

Supplementary Data 1: XL-BioID label-free protein quantification in MSK and KKT2 proximity biotinylation experiments, including limma statistical analysis and identification of proximal MSK proteins.

Supplementary Data 2: XL-BioID label-free phosphopeptides quantification in MSK and KKT2 proximity biotinylation experiments, including limma statistical analysis and identification of proximal MSK phosphosites.

Supplementary Data 3: Custom R and Python codes used to analyse the proximity labelling data.

## References

1 Cheeseman, I. M. & Desai, A. Molecular architecture of the kinetochore–microtubule interface. Nature Reviews Molecular Cell Biology 9, 33–46 (2008). 10.1038/nrm2310

2 Ishihara, T. et al. Dynamics of mitochondrial DNA nucleoids regulated by mitochondrial fission is essential for maintenance of homogeneously active mitochondria during neonatal heart development. Mol Cell Biol 35, 211–223 (2015). 10.1128/MCB.01054-14

3 Kopek, B. G., Shtengel, G., Xu, C. S., Clayton, D. A. & Hess, H. F. Correlative 3D superresolution fluorescence and electron microscopy reveal the relationship of mitochondrial nucleoids to membranes. Proc Natl Acad Sci U S A 109, 6136–6141 (2012). 10.1073/pnas.1121558109

4 Prosser, S. L. & Pelletier, L. Mitotic spindle assembly in animal cells: a fine balancing act. Nature Reviews Molecular Cell Biology 18, 187–201 (2017). 10.1038/nrm.2016.162

5 Musacchio, A. & Desai, A. A Molecular View of Kinetochore Assembly and Function. Biology 6, 5 (2017).

6 Milletti, G., Colicchia, V. & Cecconi, F. Cyclers’ kinases in cell division: from molecules to cancer therapy. Cell Death & Differentiation 30, 2035–2052 (2023). 10.1038/s41418-023-01196-z

7 Ong, J. Y. & Torres, J. Z. Dissecting the mechanisms of cell division. J Biol Chem 294, 11382–11390 (2019). 10.1074/jbc.AW119.008149

8 Legros, F., Malka, F., Frachon, P., Lombes, A. & Rojo, M. Organization and dynamics of human mitochondrial DNA. J Cell Sci 117, 2653–2662 (2004). 10.1242/jcs.01134

9 Meeusen, S. & Nunnari, J. Evidence for a two membrane-spanning autonomous mitochondrial DNA replisome. J Cell Biol 163, 503–510 (2003). 10.1083/jcb.200304040

10 Povelones, M. L. Beyond replication: division and segregation of mitochondrial DNA in kinetoplastids. Mol Biochem Parasitol 196, 53–60 (2014). 10.1016/j.molbiopara.2014.03.008

11 Wheeler, R. J., Gluenz, E. & Gull, K. The cell cycle of Leishmania: morphogenetic events and their implications for parasite biology. Mol Microbiol 79, 647–662 (2011). 10.1111/j.1365-2958.2010.07479.x

12 Hair, M. et al. Whole cell reconstructions of Leishmania mexicana through the cell cycle. PLoS Pathog 20, e1012054 (2024). 10.1371/journal.ppat.1012054

13 Ambit, A., Woods, K. L., Cull, B., Coombs, G. H. & Mottram, J. C. Morphological events during the cell cycle of Leishmania major. Eukaryot Cell 10, 1429–1438 (2011). 10.1128/EC.05118-11

14 Minocha, N., Kumar, D., Rajanala, K. & Saha, S. Kinetoplast morphology and segregation pattern as a marker for cell cycle progression in Leishmania donovani. J Eukaryot Microbiol 58, 249–253 (2011). 10.1111/j.1550-7408.2011.00539.x

15 da Silva, M. S. et al. Leishmania amazonensis promastigotes present two distinct modes of nucleus and kinetoplast segregation during cell cycle. PLoS One 8, e81397 (2013). 10.1371/journal.pone.0081397

16 Berriman, M. et al. The genome of the African trypanosome Trypanosoma brucei. Science 309, 416–422 (2005). 10.1126/science.1112642

17 Ivens, A. C. et al. The genome of the kinetoplastid parasite, Leishmania major. Science 309, 436–442 (2005). 10.1126/science.1112680

18 El-Sayed, N. M. et al. Comparative genomics of trypanosomatid parasitic protozoa. Science 309, 404–409 (2005). 10.1126/science.1112181

19 Akiyoshi, B. & Gull, K. Discovery of unconventional kinetochores in kinetoplastids. Cell 156, 1247–1258 (2014). 10.1016/j.cell.2014.01.049

20 Brusini, L., D’Archivio, S., McDonald, J. & Wickstead, B. Trypanosome KKIP1 Dynamically Links the Inner Kinetochore to a Kinetoplastid Outer Kinetochore Complex. Frontiers in Cellular and Infection Microbiology 11 (2021). 10.3389/fcimb.2021.641174

21 Nerusheva, O. O., Ludzia, P. & Akiyoshi, B. Identification of four unconventional kinetoplastid kinetochore proteins KKT22–25 in Trypanosoma brucei. Open Biology 9, 190236 (2019). doi:10.1098/rsob.190236

22 D’Archivio, S. & Wickstead, B. Trypanosome outer kinetochore proteins suggest conservation of chromosome segregation machinery across eukaryotes. J Cell Biol 216, 379–391 (2017). 10.1083/jcb.201608043

23 Ogbadoyi, E. O., Robinson, D. R. & Gull, K. A high-order trans-membrane structural linkage is responsible for mitochondrial genome positioning and segregation by flagellar basal bodies in trypanosomes. Mol Biol Cell 14, 1769–1779 (2003). 10.1091/mbc.e02-08-0525

24 Baker, N. et al. Systematic functional analysis of Leishmania protein kinases identifies regulators of differentiation or survival. Nat Commun 12, 1244 (2021). 10.1038/s41467-021-21360-8

25 Monnerat, S. et al. Identification and Functional Characterisation of CRK12:CYC9, a Novel Cyclin-Dependent Kinase (CDK)-Cyclin Complex in Trypanosoma brucei. PLoS One 8, e67327 (2013). 10.1371/journal.pone.0067327

26 Bishop, A. C. et al. A chemical switch for inhibitor-sensitive alleles of any protein kinase. Nature 407, 395–401 (2000). 10.1038/35030148

27 Carnielli, J. B. T. et al. Chemical genetics reveals Leishmania KKT2 and CRK9 kinase activity is required for cell cycle progression. bioRxiv, 2025.2009.2008.674917 (2025). 10.1101/2025.09.08.674917

28 Ogbadoyi, E., Ersfeld, K., Robinson, D., Sherwin, T. & Gull, K. Architecture of the Trypanosoma brucei nucleus during interphase and mitosis. Chromosoma 108, 501–513 (2000). 10.1007/s004120050402

29 Blum, M. et al. InterPro: the protein sequence classification resource in 2025. Nucleic Acids Res (2024). 10.1093/nar/gkae1082

30 Abramson, J. et al. Accurate structure prediction of biomolecular interactions with AlphaFold 3. Nature 630, 493–500 (2024). 10.1038/s41586-024-07487-w

31 Bishop, A. C. et al. Design of allele-specific inhibitors to probe protein kinase signaling. Curr Biol 8, 257–266 (1998). 10.1016/s0960-9822(98)70198-8

32 Bishop, A. C. et al. Generation of Monospecific Nanomolar Tyrosine Kinase Inhibitors via a Chemical Genetic Approach. Journal of the American Chemical Society 121, 627–631 (1999). 10.1021/ja983267v

33 Geoghegan, V. et al. CLK1/CLK2-driven signalling at the Leishmania kinetochore is captured by spatially referenced proximity phosphoproteomics. Commun Biol 5, 1305 (2022). 10.1038/s42003-022-04280-1

34 Oppermann, F. S. et al. Combination of chemical genetics and phosphoproteomics for kinase signaling analysis enables confident identification of cellular downstream targets. Mol Cell Proteomics 11, O111 012351 (2012). 10.1074/mcp.O111.012351

35 Branon, T. C. et al. Efficient proximity labeling in living cells and organisms with TurboID. Nat Biotechnol 36, 880–887 (2018). 10.1038/nbt.4201

36 Robinson, D. R., Sherwin, T., Ploubidou, A., Byard, E. H. & Gull, K. Microtubule polarity and dynamics in the control of organelle positioning, segregation, and cytokinesis in the trypanosome cell cycle. J Cell Biol 128, 1163–1172 (1995). 10.1083/jcb.128.6.1163

37 Ploubidou, A., Robinson, D. R., Docherty, R. C., Ogbadoyi, E. O. & Gull, K. Evidence for novel cell cycle checkpoints in trypanosomes: kinetoplast segregation and cytokinesis in the absence of mitosis. J Cell Sci 112 **(Pt** **24**), 4641–4650 (1999). 10.1242/jcs.112.24.4641

38 Hayashi, H. & Akiyoshi, B. Degradation of cyclin B is critical for nuclear division in Trypanosoma brucei. Biol Open 7 (2018). 10.1242/bio.031609

39 Sherwin, T. & Gull, K. The cell division cycle of Trypanosoma brucei brucei: timing of event markers and cytoskeletal modulations. Philos Trans R Soc Lond B Biol Sci 323, 573–588 (1989). 10.1098/rstb.1989.0037

40 Hammarton, T. C., Clark, J., Douglas, F., Boshart, M. & Mottram, J. C. Stage-specific differences in cell cycle control in Trypanosoma brucei revealed by RNA interference of a mitotic cyclin. J Biol Chem 278, 22877–22886 (2003). 10.1074/jbc.M300813200

41 Tu, X., Kumar, P., Li, Z. & Wang, C. C. An Aurora Kinase Homologue Is Involved in Regulating Both Mitosis and Cytokinesis in Trypanosoma brucei*. Journal of Biological Chemistry 281, 9677–9687 (2006). 10.1074/jbc.M511504200

42 Li, Z. et al. Identification of a novel chromosomal passenger complex and its unique localization during cytokinesis in Trypanosoma brucei. PLoS One 3, e2354 (2008). 10.1371/journal.pone.0002354

43 Black, J. A. et al. AUK3 is required for faithful nuclear segregation in the bloodstream form of Trypanosoma brucei. Mol Biochem Parasitol 261, 111664 (2025). 10.1016/j.molbiopara.2024.111664

44 Wiedeman, J., Harrison, R. & Etheridge, R. D. A limitation lifted: A conditional knockdown system reveals essential roles for Polo-like kinase and Aurora kinase 1 in Trypanosoma cruzi cell division. Proc Natl Acad Sci U S A 122, e2416009122 (2025). 10.1073/pnas.2416009122

45 Jones, N. G. et al. Regulators of Trypanosoma brucei cell cycle progression and differentiation identified using a kinome-wide RNAi screen. PLoS Pathog 10, e1003886 (2014). 10.1371/journal.ppat.1003886

46 Urbaniak, M. D. Casein kinase 1 isoform 2 is essential for bloodstream form Trypanosoma brucei. Mol Biochem Parasitol 166, 183–185 (2009). 10.1016/j.molbiopara.2009.03.001

47 Sullenberger, C., Hoffman, B., Wiedeman, J., Kumar, G. & Mensa-Wilmot, K. Casein kinase TbCK1.2 regulates division of kinetoplast DNA, and movement of basal bodies in the African trypanosome. PLoS One 16, e0249908 (2021). 10.1371/journal.pone.0249908

48 Kumar, G., Kajuluri, L. P., Gupta, C. M. & Sahasrabuddhe, A. A. A twinfilin-like protein coordinates karyokinesis by influencing mitotic spindle elongation and DNA replication in Leishmania. Mol Microbiol 100, 173–187 (2016). 10.1111/mmi.13310

49 Musacchio, A. & Salmon, E. D. The spindle-assembly checkpoint in space and time. Nat Rev Mol Cell Biol 8, 379–393 (2007). 10.1038/nrm2163

50 Aravind, L. & Koonin, E. V. The HORMA domain: a common structural denominator in mitotic checkpoints, chromosome synapsis and DNA repair. Trends Biochem Sci 23, 284–286 (1998). 10.1016/s0968-0004(98)01257-2

51 Akiyoshi, B. & Gull, K. Evolutionary cell biology of chromosome segregation: insights from trypanosomes. Open Biol 3, 130023 (2013). 10.1098/rsob.130023

52 Adams, I. R. & Kilmartin, J. V. Spindle pole body duplication: a model for centrosome duplication? Trends Cell Biol 10, 329–335 (2000). 10.1016/s0962-8924(00)01798-0

53 Tamm, T. et al. Brr6 drives the Schizosaccharomyces pombe spindle pole body nuclear envelope insertion/extrusion cycle. J Cell Biol 195, 467–484 (2011). 10.1083/jcb.201106076

54 Paoletti, A. et al. Fission yeast cdc31p is a component of the half-bridge and controls SPB duplication. Mol Biol Cell 14, 2793–2808 (2003). 10.1091/mbc.e02-10-0661

55 Selvapandiyan, A. et al. Centrin gene disruption impairs stage-specific basal body duplication and cell cycle progression in Leishmania. J Biol Chem 279, 25703–25710 (2004). 10.1074/jbc.M402794200

56 Maiato, H., Sampaio, P. & Sunkel, C. E. Microtubule-associated proteins and their essential roles during mitosis. Int Rev Cytol 241, 53–153 (2004). 10.1016/S0074-7696(04)41002-X

57 Miserey-Lenkei, S. et al. A role for the Rab6A’ GTPase in the inactivation of the Mad2-spindle checkpoint. EMBO J 25, 278–289 (2006). 10.1038/sj.emboj.7600929

58 Imada, K. & Nakamura, T. The exocytic Rabs Ypt3 and Ypt2 regulate the early step of biogenesis of the spore plasma membrane in fission yeast. Mol Biol Cell 27, 3317–3328 (2016). 10.1091/mbc.E16-03-0162

59 Poon, P. P. et al. Retrograde transport from the yeast Golgi is mediated by two ARF GAP proteins with overlapping function. EMBO J 18, 555–564 (1999). 10.1093/emboj/18.3.555

60 Strom, M., Vollmer, P., Tan, T. J. & Gallwitz, D. A yeast GTPase-activating protein that interacts specifically with a member of the Ypt/Rab family. Nature 361, 736–739 (1993). 10.1038/361736a0

61 Kraut-Cohen, J. & Gerst, J. E. Addressing mRNAs to the ER: cis sequences act up! Trends Biochem Sci 35, 459–469 (2010). 10.1016/j.tibs.2010.02.006

62 Lang, B. D., Li, A., Black-Brewster, H. D. & Fridovich-Keil, J. L. The brefeldin A resistance protein Bfr1p is a component of polyribosome-associated mRNP complexes in yeast. Nucleic Acids Res 29, 2567–2574 (2001). 10.1093/nar/29.12.2567

63 Xue, Z., Shan, X., Sinelnikov, A. & Melese, T. Yeast mutants that produce a novel type of ascus containing asci instead of spores. Genetics 144, 979–989 (1996). 10.1093/genetics/144.3.979

64 Jackson, C. L. & Kepes, F. BFR1, a multicopy suppressor of brefeldin A-induced lethality, is implicated in secretion and nuclear segregation in Saccharomyces cerevisiae. Genetics 137, 423–437 (1994). 10.1093/genetics/137.2.423

65 Bajak, K., Leiss, K., Clayton, C. E. & Erben, E. The endoplasmic reticulum-associated mRNA-binding proteins ERBP1 and ERBP2 interact in bloodstream-form Trypanosoma brucei. PeerJ 8, e8388 (2020). 10.7717/peerj.8388

66 Jiou, J. et al. Mechanism of RanGTP priming H2A-H2B release from Kap114 in an atypical RanGTP*Kap114*H2A-H2B complex. Proc Natl Acad Sci U S A 120, e2301199120 (2023). 10.1073/pnas.2301199120

67 Palacios, V., Kimble, G. C., Tootle, T. L. & Buszczak, M. Importin-9 regulates chromosome segregation and packaging in Drosophila germ cells. J Cell Sci 134 (2021). 10.1242/jcs.258391

68 Beneke, T. et al. Genome sequence of Leishmania mexicana MNYC/BZ/62/M379 expressing Cas9 and T7 RNA polymerase. Wellcome Open Res 7, 294 (2022). 10.12688/wellcomeopenres.18575.2

69 Beneke, T. et al. A CRISPR Cas9 high-throughput genome editing toolkit for kinetoplastids. R Soc Open Sci 4, 170095 (2017). 10.1098/rsos.170095

70 Weischenfeldt, J. & Porse, B. Bone Marrow-Derived Macrophages (BMM): Isolation and Applications. CSH Protoc 2008, pdb prot5080 (2008). 10.1101/pdb.prot5080

71 Valikangas, T., Suomi, T. & Elo, L. L. A systematic evaluation of normalization methods in quantitative label-free proteomics. Brief Bioinform 19, 1–11 (2018). 10.1093/bib/bbw095

72 Ritchie, M. E. et al. limma powers differential expression analyses for RNA-sequencing and microarray studies. Nucleic Acids Res 43, e47 (2015). 10.1093/nar/gkv007

