## Supplementary information for "The mitotic spindle kinase MSK co-ordinates segregation of the nucleus and kinetoplast in *Leishmania mexicana*"

**Supplementary Table 1. Sequence of oligonucleotides used to generate sgRNA and DNA repair template for CRISPR-Cas9 edited lines.**

| Oligo ID | Engineered cell line | Sequence (5’ → 3’) | Description |
|---|---|---|---|
| OL6137 (R) | N/A | AAAAGCACCGACTCGGTGCCACTTTTTCAAGTTGATAACGGACTAGCCTTATTTTAACTTGCTATTTCTAGCTCTAAAAC | Reverse oligo to generate 5’sgRNA and 3’sgRNA |
| OL11595 (F) | AS MSK$^{M192G/A}$ | GAAATTAATACGACTCACTATAGGTCCACAGATTGTTCGATTTCGTTTTAGAGCTAGAAATAGC | 5’sgRNA |
| OL11596 (F) | | GAAATTAATACGACTCACTATAGGGAGTTGATGGACAGCGACGTGTTTTAGAGCTAGAAATAGC | 3’sgRNA |
| OL6696 (F) | 3xMyc::mNG::MSK | GAAATTAATACGACTCACTATAGGGGTGAAAAGCTAAACAGTAAGTTTTAGAGCTAGAAATAGC | 5’sgRNA |
| OL6694 (F) | 3xMyc::mNG::MSK_AS | ATATAGATTGCAGAGAAGACTTGTATACCAGTATAATGCAGACCTGCTGC | Upstream forward to generate repair template |
| OL6695 (R) | 3xMyc::mT::MSK_AS | ATGGGTTTGAAGAATACCTGATGACTTCATACTACCCGATCCTGATCCAG | Upstream reverse to generate repair template |
| OL13560 (F) | *Δrab3gap*<br>3xHA::Rab3GAP | GAAATTAATACGACTCACTATAGGAGGTGATGCTGGTGTTCTTGGTTTTAGAGCTAGAAATAGC | 5’sgRNA |
| OL13564 (F) | | GAAATTAATACGACTCACTATAGGGAGAGATGGGGGAGGGCAGGGTTTTAGAGCTAGAAATAGC | 3’sgRNA |
| OL13557 (F) | | CCGCCCGTGTCATCTTGTGCACCGCCACCAGTATAATGCAGACCTGCTGC | Upstream forward to generate repair template |
| OL13559 (R) | | CGACTCGCTGCGCGCATACGTCGTCTCCATACCAGAACCAGAACCAGAACCAGAACCAGCGTAATCTG | Upstream reverse to generate repair template |
| OL13563 (R) | | GGGGCCGATGATGCGAGCGAGTGAGCCCCGCCAATTTGAGAGACCTGTGC | Downstream reverse to generate repair template |
| OL13568 (F) | *Δnsp*<br>NSP::3xHA | GAAATTAATACGACTCACTATAGGGATCCCCACGTGACTTCTCTGTTTTAGAGCTAGAAATAGC | 5’sgRNA |
| OL13572 (F) | | GAAATTAATACGACTCACTATAGGTATGGTTACCGGGAGCGTAGGTTTTAGAGCTAGAAATAGC | 3’sgRNA |
| OL13565 (F) | | CGGCATTGCTTTTTTGCGGCACGGTCGATTGTATAATGCAGACCTGCTGC | Upstream forward to generate repair template |
| OL13570 (F) | | CCGGGGACGAAGGCGACTGAGGAGGCTGCCGGTAGTGGATCGGGTAGTGGATCGGGTAGTATGTACCCATACG | Downstream forward to generate repair template |
| OL13571 (R) | | TCCAAACTAAGGCCACAAGACTCAATGAACCCAATTTGAGAGACCTGTGC | Downstream reverse to generate repair template |
| OL13576 (F) | *Δarfgap1*<br>ARFGAP1::3xHA | GAAATTAATACGACTCACTATAGGTGCGTATGCTCTGCGTTCCAGTTTTAGAGCTAGAAATAGC | 5’sgRNA |
| OL13580 (F) | | GAAATTAATACGACTCACTATAGGACCCCTGCGAGTCATGAAAAGTTTTAGAGCTAGAAATAGC | 3’sgRNA |
| OL13573 (F) | | TTCTCCTCCACCGTTGGCGTGCACCCACCGGTATAATGCAGACCTGCTGC | Upstream forward to generate repair template |
| OL13578 (F) | | GACGAGTGGGCTTGGGATGACGAGAACATCGGTAGTGGATCGGGTAGTGGATCGGGTAGTATGTACCCATACG | Downstream forward to generate repair template |
| OL13579 (R) | | CAAGCAACACGCGGCGAAGCTTCTCGTCCCCCAATTTGAGAGACCTGTGC | Downstream reverse to generate repair template |
| OL13552 (F) | *Δlmxm.36.2560*<br>LmxM.36.2560::3xHA | GAAATTAATACGACTCACTATAGGATTACTAGGTAGAGGAGGAGGTTTTAGAGCTAGAAATAGC | 5’sgRNA |
| OL13556 (F) | | GAAATTAATACGACTCACTATAGGGTCACGAAGCCCTTCCTTCTGTTTTAGAGCTAGAAATAGC | 3’sgRNA |
| OL13549 (F) | | GTGGGGCCAGTCATCTGCCTCTGCCTTCCTGTATAATGCAGACCTGCTGC | Upstream forward to generate repair template |
| OL13554 (F) | | CGCCCCCAGCTCGTGGTCCTCGCGGCTCTCGGTAGTGGATCGGGTAGTGGATCGGGTAGTATGTACCCATACG | Downstream forward to generate repair template |
| OL13555 (R) | | ATCAAACAACAGCCACACACACCCAACGCACCAATTTGAGAGACCTGTGC | Downstream reverse to generate repair template |
| OL14380 (F) | PM Rab3GAP$^{S804S/A}$ | GAAATTAATACGACTCACTATAGGGCTCCTGCGCTACGCACAAGGTTTTAGAGCTAGAAATAGC | 5’sgRNA |
| OL14381 (F) | | GAAATTAATACGACTCACTATAGGGGACAACGGTTCGCCGACTAGTTTTAGAGCTAGAAATAGC | 3’sgRNA |

| Oligo ID | Engineered cell line | Sequence (5' → 3') | Description |
|---|---|---|---|
| OL14382 (F) | PM NSP$^{T484S/A}$ | GAAATTAATACGACTCACTATAGGATCCTCTGGAGCCTTTGGCGGTTTTAGAGCTAGAAATAGC | 5'sgRNA |
| OL13572 (F) | | GAAATTAATACGACTCACTATAGGTATGGTTACCGGGAGCGTAGGTTTTAGAGCTAGAAATAGC | 3'sgRNA |
| OL14383 (F) | PM ARFGAP1$^{S155S/A/E}$ | GAAATTAATACGACTCACTATAGGAGTCGCGTGTACCAGGCTGGGTTTTAGAGCTAGAAATAGC | 5'sgRNA |
| OL14384 (F) | | GAAATTAATACGACTCACTATAGGAAGCGGCTTGGGTCCGTTTGGTTTTAGAGCTAGAAATAGC | 3'sgRNA |
| OL14377 (F) | PM LmxM.36.2560$^{S552S/A}$ | GAAATTAATACGACTCACTATAGGGACGATCCGGCGGTTGGTGGGTTTTAGAGCTAGAAATAGC | 5'sgRNA |
| OL14378 (F) | | GAAATTAATACGACTCACTATAGGACCTGCGCGCGCGTCTCGTCGTTTTAGAGCTAGAAATAGC | 3'sgRNA |

R, reverse oligo; F, forward oligo; N/A, not applicable; AS, analog-sensitive kinase; PM, phosphosite mutation; Δ, knockout target gene.

Rab3GAP [Rab3 GTPase-activating protein catalytic subunit, LmxM.25.1490]; NSP [nuclear segregation protein, LmxM.31.0020]; ARFGAP1 [ADP-ribosylation factor GTPase activating protein 1, LmxM.31.1230]; hypothetical protein, conserved [LmxM.36.2560].

The R sgRNA oligo OL6137 holds the Cas9 scaffold (black) and the complementary sequence to the forward primer (blue). The F oligo sequences to generate sgRNA was coloured as follow: T7 promoter in black; sgRNA target sites in red; complementary sequence to sgRNA scaffold in blue.

The upstream and downstream oligo sequences to generate DNA repair template was coloured as follow: 30 bp for homology direct recombination in red; complementary sequence to the template plasmid in blue.

**Supplementary Table 2. Sequence of DNA repair templates used to engineer analog-sensitive kinases or phosphosite mutants in *Leishmania*.**

| Engineered cell line | Sequence (ssDRT or dsDRT) |
|---|---|
| AS MSK$^{M192G}$ | TTATCTTGGGCCTCATCCACAGATTGTGCGCTTCCTCGGCAGCTACACAACGTCGAAGAACACGAGCTTTTTTACA**GGC**GAACTTATGGACAGTGATGTTGGACGGGAGTTGAGGGAAGG (ssDRT) |
| AS MSK$^{M192A}$ | TTATCTTGGGCCTCATCCACAGATTGTGCGCTTCCTCGGCAGCTACACAACGTCGAAGAACACGAGCTTTTTTACA**GCG**GAACTTATGGACAGTGATGTTGGACGGGAGTTGAGGGAAGG (ssDRT) |
| PM Rab3GAP$^{S804A}$ | TTGGGCATGCCTCCAGAAGACCCCCCCGTCGACCTTGGTCGGCCGCTGCTGACGCAGAAGCTGCAGCTCCTGCGGTATGCTCAAGCCGCTCTCCTGCGTTCATTGGAGGACAACGGT**GCG**CCCACAACCGCAACGCCCGC<br>GGCGAGCACGAAGGCGGAAGCGGACGGGGCAGCTACCTCGCTGACCACGAGCCCCGTCCG (dsDRT)<br>OL14435 (F) TTGGGCATGCCTCCAGAAGACCCCCCCGTCGACCTTGGTCGGCCGCTGCTGACGCAGAAGCTGCAGCTCCTGCGGTATGCTCAAGCCGCTCTCCTGCGTTCATTGGAGGACAACGG<br>OL14436 (R) CGGACGGGGCTCGTGGTCAGCGAGGTAGCTGCCCCGTCCGCTTCCGCCTTCGTGCTCGCCGCGGGCGTTGCGGTTGTGGGCGCACCGTTGTCCTCCAATGAACG |
| PM Rab3GAP$^{S804S}$ | TTGGGCATGCCTCCAGAAGACCCCCCCGTCGACCTTGGTCGGCCGCTGCTGACGCAGAAGCTGCAGCTCCTGCGGTATGCTCAAGCCGCTCTCCTGCGTTCATTGGAGGACAACGGT**TCC**CCCACAACCGCAACGCCCGC<br>GGCGAGCACGAAGGCGGAAGCGGACGGGGCAGCTACCTCGCTGACCACGAGCCCCGTCCG (dsDRT)<br>OL14435 (F) TTGGGCATGCCTCCAGAAGACCCCCCCGTCGACCTTGGTCGGCCGCTGCTGACGCAGAAGCTGCAGCTCCTGCGGTATGCTCAAGCCGCTCTCCTGCGTTCATTGGAGGACAACGG<br>OL14437 (R) CGGACGGGGCTCGTGGTCAGCGAGGTAGCTGCCCCGTCCGCTTCCGCCTTCGTGCTCGCCGCGGGCGTTGCGGTTGTGGGGGAACCGTTGTCCTCCAATGAACG |
| PM NSP$^{T484A}$ | CTCTCTAGTGGCGACGACGAAGACGAGGAGGCGCCGGACGACGGGGGTGACACGCCATCCCCTAAAGCACCAGAGGATGCGGAG**GCG**CCGGGGACGAAGGCGACTGAGGAGGCTGCCTAGCCACTTCGGTCGCGCTAACC<br>ATAGTTCATTGAGTCTTGTGGCCTTAGTTTGGAAAGAGTTGGTGAGGGAGAAGAGAGACG (dsDRT)<br>OL14440 (F) CTCTCTAGTGGCGACGACGAAGACGAGGAGGCGCCGGACGACGGGGGTGACACGCCATCCCCTAAAGCACCAGAGGATGCGGAGGCGCCGGGGACGAAGGCGACTGAGGAGGCTGCCTAG<br>OL14438 (R) CGTCTCTCTTCTCCCTCACCAACTCTTTCCAAACTAAGGCCACAAGACTCAATGAACTATGGTTAGCGCGACCGAAGTGGCTAGGCAGCCTCCTCAGTC |
| PM NSP$^{T484S}$ | CTCTCTAGTGGCGACGACGAAGACGAGGAGGCGCCGGACGACGGGGGTGACACGCCATCCCCTAAAGCACCAGAGGATGCGGAG**ACC**CCGGGGACGAAGGCGACTGAGGAGGCTGCCTAGCCACTTCGGTCGCGCTAACC<br>ATAGTTCATTGAGTCTTGTGGCCTTAGTTTGGAAAGAGTTGGTGAGGGAGAAGAGAGACG (dsDRT)<br>OL14439 (F) CTCTCTAGTGGCGACGACGAAGACGAGGAGGCGCCGGACGACGGGGGTGACACGCCATCCCCTAAAGCACCAGAGGATGCGGAGACCCCGGGGACGAAGGCGACTGAGGAGGCTGCCTAG<br>OL14438 (R) CGTCTCTCTTCTCCCTCACCAACTCTTTCCAAACTAAGGCCACAAGACTCAATGAACTATGGTTAGCGCGACCGAAGTGGCTAGGCAGCCTCCTCAGTC |
| PM ARFGAP1$^{S155A}$ | CTCGAGTCCGAGGCGCTTGGCAGGCCTTTCAACGAGGCGTCCTGGCAGCCACCGGCGTGGTATACCCGACTCAAAGCCGCGGCGAGCCTCTCAGGGCCG**GCG**CCAACGTCATCGTACCCGCAGACCGATCCTAGTCGCTT<br>CGCCGGTGTTGGCTCGAATGGGCATCCGCACGTGATGCCGGGCAGCGGGGGCGGCGACAG (dsDRT)<br>OL14442 (F) CTCGAGTCCGAGGCGCTTGGCAGGCCTTTCAACGAGGCGTCCTGGCAGCCACCGGCGTGGTATACCCGACTCAAAGCCGCGGCGAGCCTCTCAGGGCCGGCGCCAACGTCATCGTACCCG<br>OL14441 (R) CTGTCGCCGCCCCCGCTGCCCGGCATCACGTGCGGATGCCCATTCGAGCCAACACCGGCGAAGCGACTAGGATCGGTCTGCGGGTACGATGACGTTGG |
| PM ARFGAP1$^{S155E}$ | CTCGAGTCCGAGGCGCTTGGCAGGCCTTTCAACGAGGCGTCCTGGCAGCCACCGGCGTGGTATACCCGACTCAAAGCCGCGGCGAGCCTCTCAGGGCCG**GAG**CCAACGTCATCGTACCCGCAGACCGATCCTAGTCGCTT<br>CGCCGGTGTTGGCTCGAATGGGCATCCGCACGTGATGCCGGGCAGCGGGGGCGGCGACAG (dsDRT)<br>OL14576 (F) CTCGAGTCCGAGGCGCTTGGCAGGCCTTTCAACGAGGCGTCCTGGCAGCCACCGGCGTGGTATACCCGACTCAAAGCCGCGGCGAGCCTCTCAGGGCCGGAGCCAACGTCATCGTACCCG<br>OL14441 (R) CTGTCGCCGCCCCCGCTGCCCGGCATCACGTGCGGATGCCCATTCGAGCCAACACCGGCGAAGCGACTAGGATCGGTCTGCGGGTACGATGACGTTGG |
| PM ARFGAP1$^{S155S}$ | CTCGAGTCCGAGGCGCTTGGCAGGCCTTTCAACGAGGCGTCCTGGCAGCCACCGGCGTGGTATACCCGACTCAAAGCCGCGGCGAGCCTCTCAGGGCCG**TCG**CCAACGTCATCGTACCCGCAGACCGATCCTAGTCGCTT<br>CGCCGGTGTTGGCTCGAATGGGCATCCGCACGTGATGCCGGGCAGCGGGGGCGGCGACAG (dsDRT)<br>OL14443 (F) CTCGAGTCCGAGGCGCTTGGCAGGCCTTTCAACGAGGCGTCCTGGCAGCCACCGGCGTGGTATACCCGACTCAAAGCCGCGGCGAGCCTCTCAGGGCCGTCGCCAACGTCATCGTACCCG<br>OL14441 (R) CTGTCGCCGCCCCCGCTGCCCGGCATCACGTGCGGATGCCCATTCGAGCCAACACCGGCGAAGCGACTAGGATCGGTCTGCGGGTACGATGACGTTGG |
| PM LmxM.36.2560$^{S552A}$ | TGGCGAAGGCGCAAACGTACGTGGCCACGTTCGTGGTGCCGGTGATGTGGTGGTTTCTGACGATCCGGCGGTTAGTCGCCGAAGTCAGAGAAAAGATGGCGCCG**GCG**CCGACGACGGGTGCCCCCGATGAAACCCGCGCG<br>CAGGTAGAGGATTTAGTGCGACCGTACGCCACCCTAGAGCAGATCCTGTTCGCGTCGCAG (dsDRT)<br>OL14432 (F) TGGCGAAGGCGCAAACGTACGTGGCCACGTTCGTGGTGCCGGTGATGTGGTGGTTTCTGACGATCCGGCGGTTAGTCGCCGAAGTCAGAGAAAAGATGGC<br>OL14433 (R) CTGCGACGCGAACAGGATCTGCTCTAGGGTGGCGTACGGTCGCACTAAATCCTCTACCTGCGCGCGGGTTTCATCGGGGGCACCCGTCGTCGGCGCCGGCGCCATCTTTTCTCTGACTTC |
| PM LmxM.36.2560$^{S552S}$ | TGGCGAAGGCGCAAACGTACGTGGCCACGTTCGTGGTGCCGGTGATGTGGTGGTTTCTGACGATCCGGCGGTTAGTCGCCGAAGTCAGAGAAAAGATGGCGCCG**TCC**CCGACGACGGGTGCCCCCGATGAAACCCGCGCG<br>CAGGTAGAGGATTTAGTGCGACCGTACGCCACCCTAGAGCAGATCCTGTTCGCGTCGCAG (dsDRT)<br>OL14432 (F) TGGCGAAGGCGCAAACGTACGTGGCCACGTTCGTGGTGCCGGTGATGTGGTGGTTTCTGACGATCCGGCGGTTAGTCGCCGAAGTCAGAGAAAAGATGGC<br>OL14434 (R) CTGCGACGCGAACAGGATCTGCTCTAGGGTGGCGTACGGTCGCACTAAATCCTCTACCTGCGCGCGGGTTTCATCGGGGGCACCCGTCGTCGGGGACGGCGCCATCTTTTCTCTGACTTC |

AS, analog-sensitive kinase; PM, phosphosite mutation; ssDRT, single-stranded DNA repair template; dsDRT, double-stranded DNA repair template.

Rab3GAP [Rab3 GTPase-activating protein catalytic subunit, LmxM.25.1490]; NSP [nuclear segregation protein, LmxM.31.0020]; ARFGAP1 [ADP-ribosylation factor GTPase activating protein 1, LmxM.31.1230]; hypothetical protein, conserved [LmxM.36.2560].

Repair template sequence was coloured as follow: homology arm in black; recoded codons in blue; and mutated target codon in red.

Sequences in grey below each repair template correspond to the oligonucleotides (OL) used to generate dsDRT by PCR.

**Supplementary Table 3. Sequence of oligonucleotides used to screen the CRISPR-Cas9 engineered *L. mexicana* cell lines.**

| Oligo ID | Engineered cell line | Sequence | Description |
|---|---|---|---|
| OL11599 (F) | AS MSK$^{M192G/A}$ | TTTGGGAAAAGGTGGAAGTG | PCR followed by restriction site digestion to screen analog sensitive mutant – Sanger sequencing |
| OL11600 (R) | | CCGAGAAATGCGACTAAAGC | PCR followed by restriction site digestion to screen analog sensitive mutant – Sanger sequencing |
| OL12754 (F) | 3xMyc::mNG::MSK | CGGGCGATTCAGCAAGAAAA | CRISPR-Cas9 endogenous tagged cell line screening |
| OL12755 (R) | 3xMyc::mNG::MSK_AS | GCAGCAGGTCTGCATTATAC | CRISPR-Cas9 endogenous tagged cell line screening |
| OL12756 (R) | 3xMyc::mT::MSK | TCCCTGTAGACCCGTTAGCA | CRISPR-Cas9 endogenous tagged cell line screening |
| OL13599 (F) | | TCCGTTCACACACACAGTCC | CRISPR-Cas9 Knocked out and endogenous tagged cell lines screening |
| OL13600 (R) | Δ*rab3gap* | GACTGCCAGGTACTGCTCAA | CRISPR-Cas9 endogenous tagged cell line screening |
| OL13602 (R) | 3xHA::Rab3GAP | CGATGTTTGAGGCACGGCA | CRISPR-Cas9 Knocked out cell line screening |
| OL14179 (F) | PM Rab3GAP$^{S804S/A}$ | GCCATCTCGATCCCTTCCAG | CRISPR-Cas9 Knocked out cell line screening<br>PCR followed by restriction site digestion to screen analog sensitive mutant – Sanger sequencing |
| OL14180 (F) | | GTGCCACTGCACAAAGTCAG | CRISPR-Cas9 Knocked out cell line screening<br>PCR followed by restriction site digestion to screen analog sensitive mutant – Sanger sequencing |
| OL13603 (F) | | CCTGCTTGGGTGGAATGACT | CRISPR-Cas9 Knocked out cell line screening |
| OL13605 (F) | | GTTGATCCGGTACTGTCGCT | CRISPR-Cas9 endogenous tagged cell line screening |
| OL13606 (R) | Δ*nsp*<br>NSP::3xHA | CATGCACCCAGGGAATGACA | CRISPR-Cas9 Knocked out and endogenous tagged cell lines screening<br>PCR followed by restriction site digestion to screen analog sensitive mutant – Sanger sequencing |
| OL14181 (F) | PM NSP$^{T484S/A}$ | CGCTCAGGACGCTGAAATTG | CRISPR-Cas9 Knocked out cell line screening<br>PCR followed by restriction site digestion to screen analog sensitive mutant – Sanger sequencing |
| OL14183 (F) | | ATCCTGAACCGGGAGTACGA | Sanger sequencing |
| OL14184 (R) | | GCACCTCAATGTGGCGAATC | CRISPR-Cas9 Knocked out cell line screening |
| OL13607 (F) | | AGGGGCTCTCTGTGCATCAT | CRISPR-Cas9 Knocked out cell line screening<br>PCR followed by restriction site digestion to screen analog sensitive mutant – Sanger sequencing |
| OL13609 (F) | Δ*arfgap1* | ACCGCACTTCCTCAAATCCA | CRISPR-Cas9 endogenous tagged cell line screening |
| OL13610 (R) | ARFGAP1::3xHA | ACACCGACACAGAGGTAGGA | CRISPR-Cas9 Knocked out and endogenous tagged cell lines screening |
| OL14185 (F) | PM ARFGAP1$^{S155S/A/E}$ | ATGGACGGCTGGATGAAGTG | CRISPR-Cas9 Knocked out cell line screening – Sanger sequencing |
| OL14186 (R) | | CCTTGCGCTATTCGCTGTTG | CRISPR-Cas9 Knocked out cell line screening – Sanger sequencing |
| OL14188 (R) | | TTGAGGAAGTGCGGTACTGG | PCR followed by restriction site digestion to screen analog sensitive mutant – Sanger sequencing |
| OL13595 (F) | | TTAACAGCCCATCTCCTCCT | CRISPR-Cas9 Knocked out cell line screening |
| OL13597 (F) | Δ*lmxm.36.2560* | AACGAGTGCTGGGATGACTG | CRISPR-Cas9 endogenous tagged cell line screening |
| OL13598 (R) | LmxM.36.2560::3xHA | CTAAGCGCAACGACATGCAA | CRISPR-Cas9 Knocked out and endogenous tagged cell lines screening<br>PCR followed by restriction site digestion to screen analog sensitive mutant – Sanger sequencing |
| OL14174 (R) | PM LmxM.36.2560$^{S552S/A}$ | TCAGGATGCCGTTCGTATCG | CRISPR-Cas9 Knocked out cell line screening |
| OL14175 (F) | | TCCGTTTCTAGCGTGCTTGT | CRISPR-Cas9 Knocked out cell line screening |
| OL14379 (F) | | CGATACGAACGGCATCCTGA | PCR followed by restriction site digestion to screen analog sensitive mutant – Sanger sequencing |

| Oligo ID | Engineered cell line | Sequence | Description |
|---|---|---|---|
| OL9369 (R) | N/A | GCAGCAGGTCTGCATTATAC | Integration of the DNA repair template in the CRISPR-Cas9 knocked-out cell lines |
| OL12716 (R) | N/A | TCTCGCGTAGCAAGACGAACCGTTGG | Integration of the DNA repair template in the CRISPR-Cas9 N/C-terminus endogenous tagged cell lines with puromycin resistant marker |
| OL12757 (F) | N/A | GCACAGGTCTCTCAAATTGG | Integration of the DNA repair template in the CRISPR-Cas9 C-terminus endogenous tagged cell lines |

R, reverse oligo; F, forward oligo; N/A, not applicable; AS, analog-sensitive kinase; PM, phosphosite mutation; Δ, knockout target gene.

Rab3GAP [Rab3 GTPase-activating protein catalytic subunit, LmxM.25.1490]; NSP [nuclear segregation protein, LmxM.31.0020]; ARFGAP1 [ADP-ribosylation factor GTPase activating protein 1, LmxM.31.1230]; hypothetical protein, conserved [LmxM.36.2560].

**MSK**

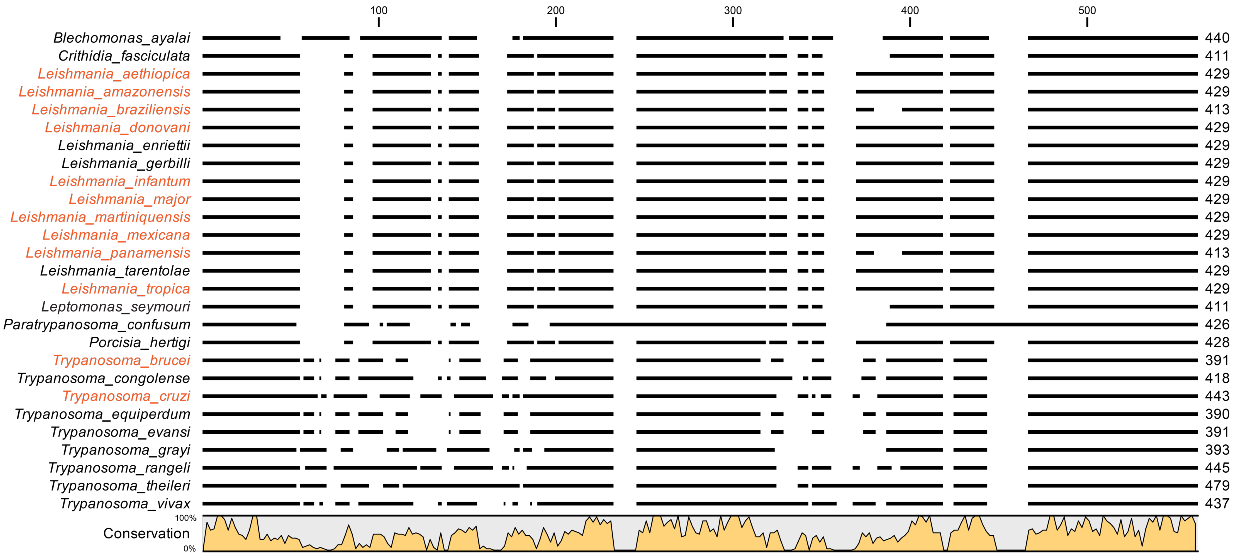

**Supplementary Fig. 1 MSK is conserved in Trypanosomatida lineage.** *L. mexicana* MSK (LmxM.24.0670) protein kinase domain sequence and its orthologues from trypanosomatid reference strains available on TriTrypDB were aligned in CLC Genomics Workbench v.22 using the Clustal Omega algorithm. The consensus line graph is shown below the corresponding alignment. The trypanosomatids species that are human pathogens are highlighted in orange. The upstream ATG start codon in-frame with the annotated CDS on TriTrypDB for *L. mexicana* MSK, previously reported by Fiebig et al., 2015[1], were manually added to the MSK DNA sequence. This extended CDS comprises the N-terminus of MSK kinase domain. By DNA sequence analysis, this missing sequence was also identified and manually inserted for: *Leishmania braziliensis* MHOM/BR/75/M2904; *Leishmania donovani* BPK282A1; *Leishmania enriettii* strain LEM3045; *Leishmania infantum* JPCM5; *Leishmania tropica* L590; and *Leptomonas seymouri* ATCC 30220.

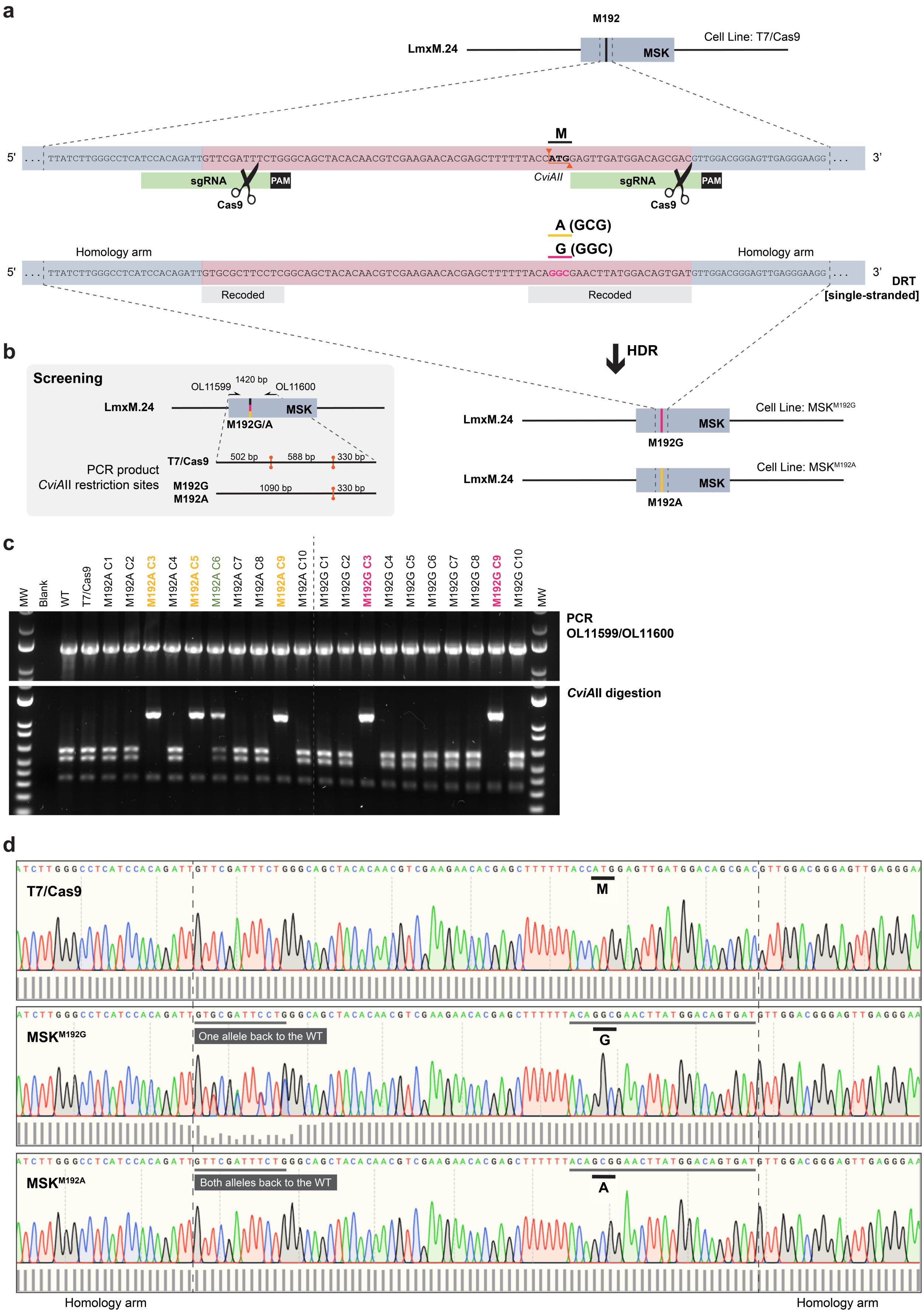

**Supplementary Fig. 2 CRISPR-Cas9–mediated precision genome editing of MSK to engineer an analog-sensitive protein kinase. a** Strategy to precisely edit the *Leishmania* genome using CRISPR-Cas9. MSK gatekeeper

(M) was replaced by Glycine (G) or Alanine (A), using two guide RNAs (sgRNA) and a 120 bp DNA repair template (DRT) containing silent mutations (Recoded) and the gatekeeper codon mutation, which removed a *CviAII* restriction site from the parental cell line T7/Cas9 allowing the genotype screening. PAM, protospacer adjacent motif; HDR, homology-directed repair; OL, oligonucleotide as listed in Supplementary Table 3. **b** Screening strategy used to genotype the engineered AS kinases. **c** Screening genotype, as described in section b, of ten clones (C1 – C10) for each mutation attempted. The genotypes are highlighted as follow: Black, wild-type; Magenta, MSK$^{M192G}$; Yellow, MSK$^{M192A}$; Green, MSK$^{M192A/M}$ or a non-clonal of MSK$^{M192A}$ and wild-type. **d** Sanger sequencing of MSK confirming the replacement of the methionine gatekeeper in T7/Cas9 progenitor for glycine or alanine in MSK$^{M192G}$ and MSK$^{M192A}$ mutants, respectively. Part of the recoded region in the DRT was not present in one or both alleles of the mutants (dark grey boxes), which retained the wild-type sequence. However, this did not alter the amino acid sequence. Sequencing files were opened on SnapGene v.7.2 and the bars on the bottom indicates the sequence quality per base.

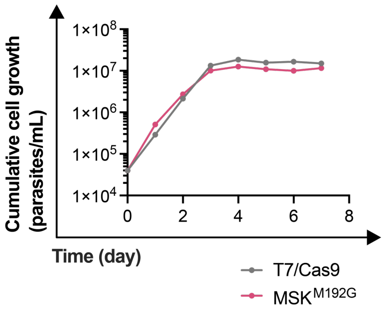

**Supplementary Fig. 3 *L. mexicana* MSK analog-sensitive and progenitor line growth curves.** The growth curve for *L. mexicana* promastigote, grown in HOMEM medium supplemented with 10% hi-FBS, was set at 4x10$^{4}$ cell mL$^{-1}$ and the cumulative cell growth was measured daily by cell counting in Neubauer chamber. The growth rate was calculated in the logarithmic area of the growth curve (0 – 96 h) and depicted as mean ± SEM: T7/Cas9, 1.53 ± 0.007; and MSK$^{M192G}$, 1.44 ± 0.011. Difference in growth rate was calculated using the non-parametric Kruskal–Wallis' test, comparing the mutants with the progenitor line T7/Cas9: no significance difference was found.

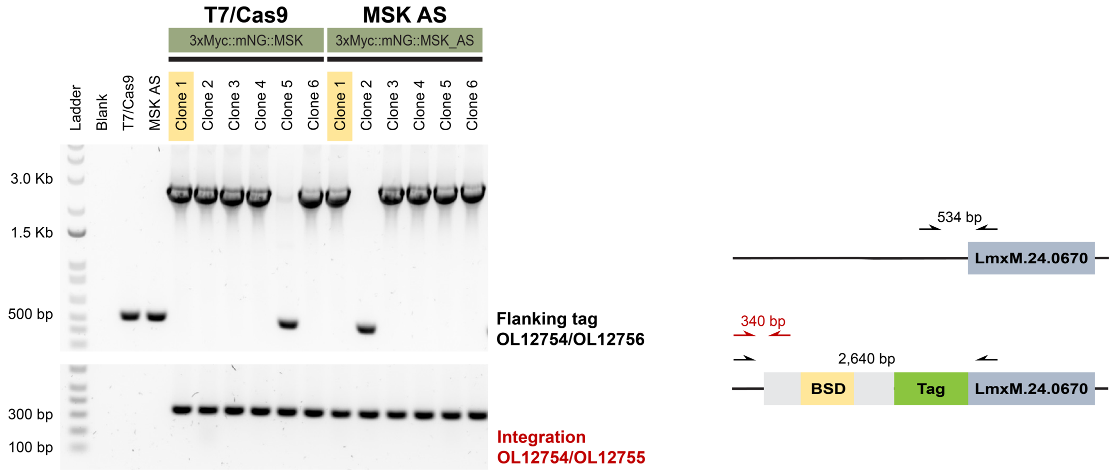

**Supplementary Fig. 4 PCR screening of the cell lines with 3xMyc::mNG fused to MSK.** Using CRISPR Cas9, the T7/Cas9 and MSK AS lines had both alleles of the MSK endogenously tagged with 3xMyc::mNeonGreen (mNG) to generate the 3xMyc::mNG::MSK and 3xMyc::mNG::MSK_AS cell lines. Repair template containing blasticidin drug resistance marker and linear DNA fragment for *in vivo* transcription of the single guide RNA were transfected into the parental line. After the overnight recovery time, transfected cells were selected with blasticidin and cloned by serial dilution. PCR amplification was used to determine the integration of the repair template in the right position in both alleles. PCR strategies and DNA product size expected for each PCR are shown in the diagram on the right. OL, oligonucleotide as listed in Supplementary Table 3. The clones highlighted in yellow were selected to perform the further experiments.

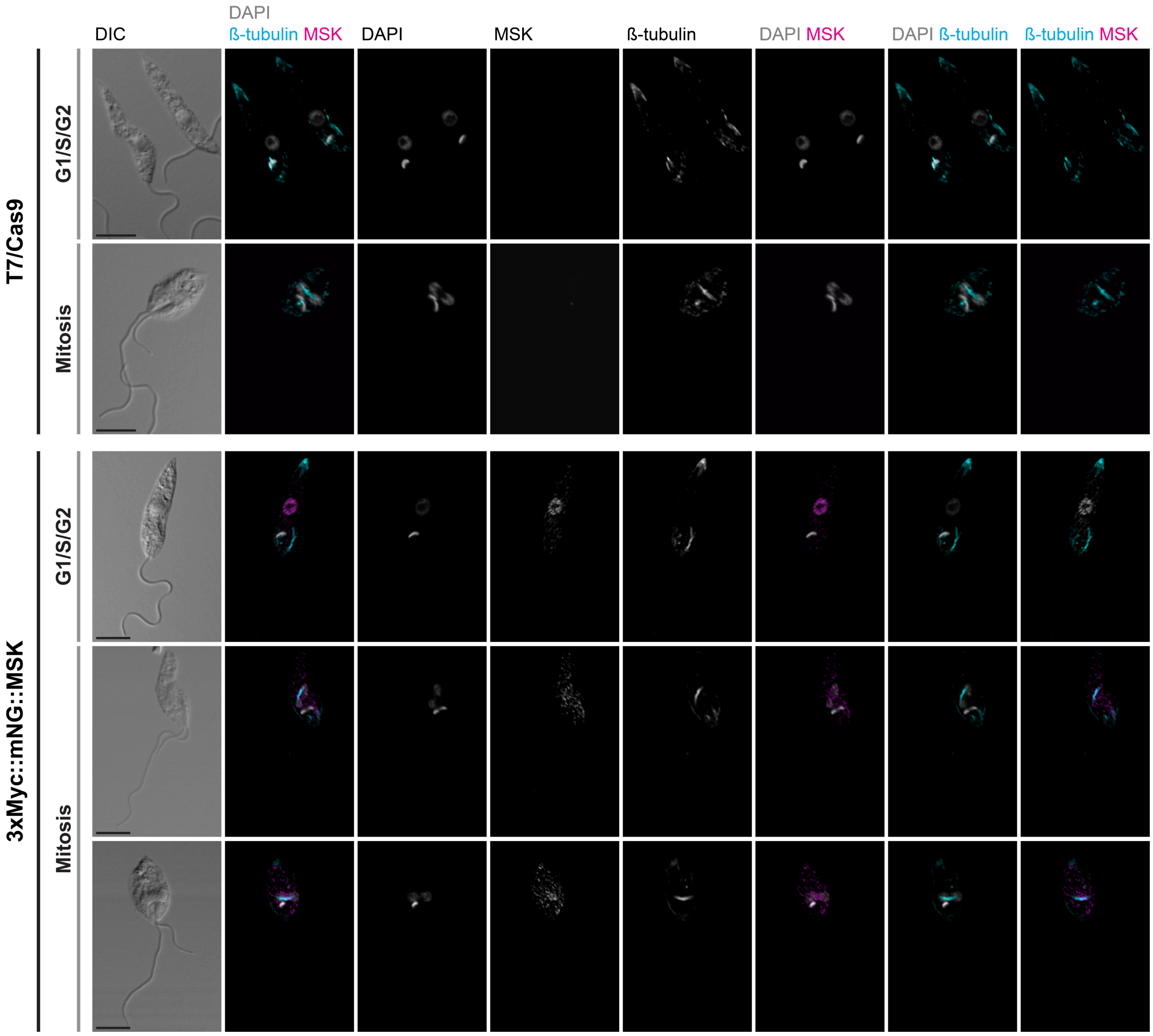

**Supplementary Fig. 5 MSK localization in *L. mexicana* promastigotes.** Promastigotes stained with KMX-1 antibody to recognise β-tubulin, anti-Myc to recognise the endogenously tagged 3xMyc::mNG::MSK, and counterstained with DAPI to visualise DNA. Representative confocal fluorescence micrographs showing the parental control T7/Cas9 and the MSK endogenously tagged parasites in different cell cycle stages. The channel and the colours used for each marking are indicated on the top. DIC, the Nomarsky differential interference contrast. Scale bars, 5 µm.

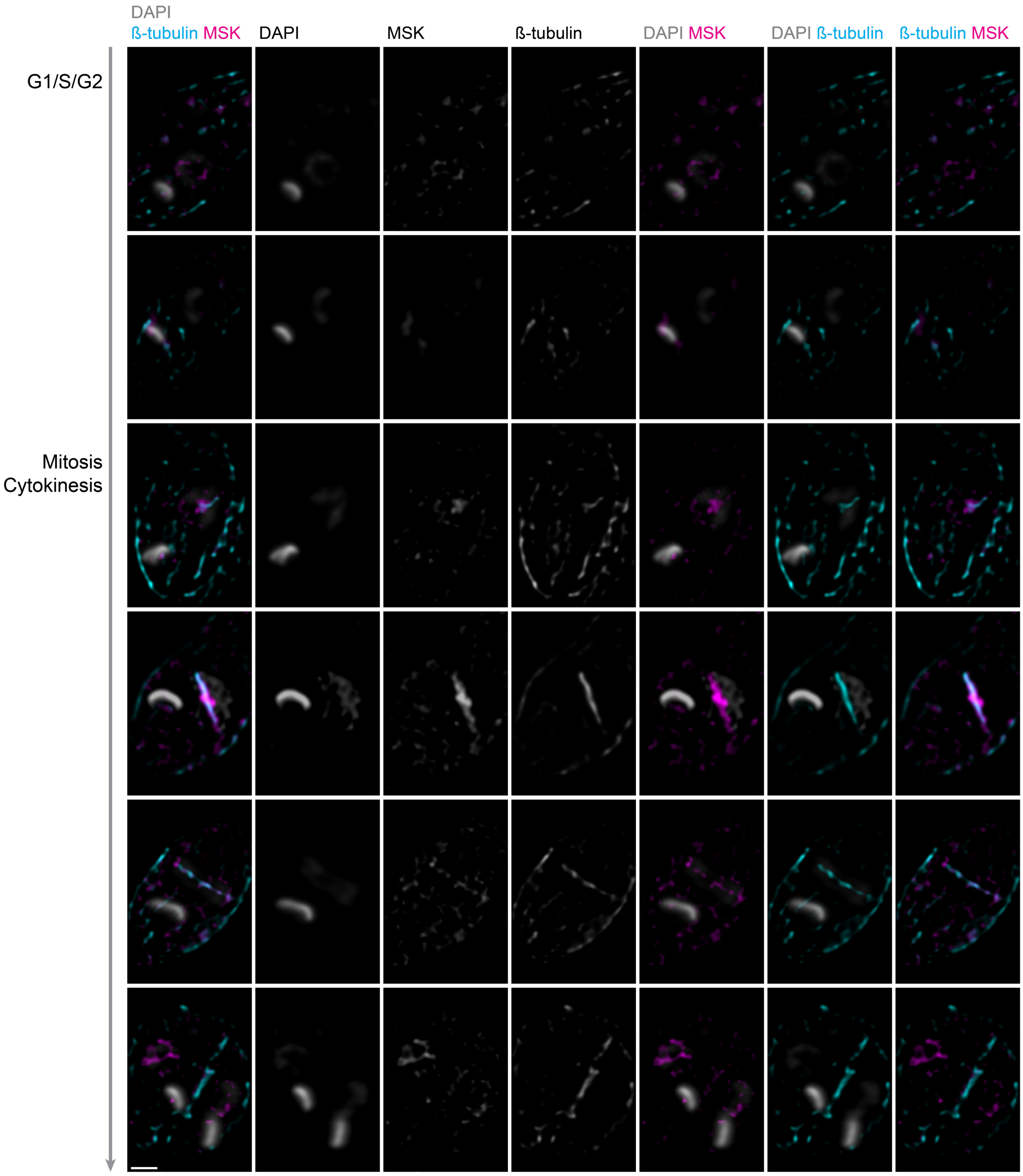

**Supplementary Fig. 6 High resolution microscopy of *L. mexicana* promastigotes endogenously expressing 3xMyc::mNG::MSK_AS.** High resolution microscopy of promastigotes stained with KMX-1 antibody to recognise β-tubulin, anti-Myc to recognise the endogenously tagged 3xMyc::mNG::MSK_AS, and counterstained with DAPI to visualise DNA. Representative fluorescence micrographs showing parasites in different cell cycle stages. The channel and the colours used for each marking are indicated on the top. Scale bars, 5 µm.

**a**

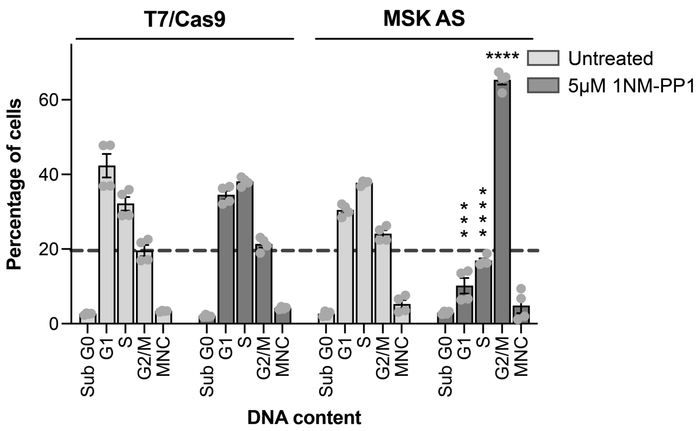

**b**

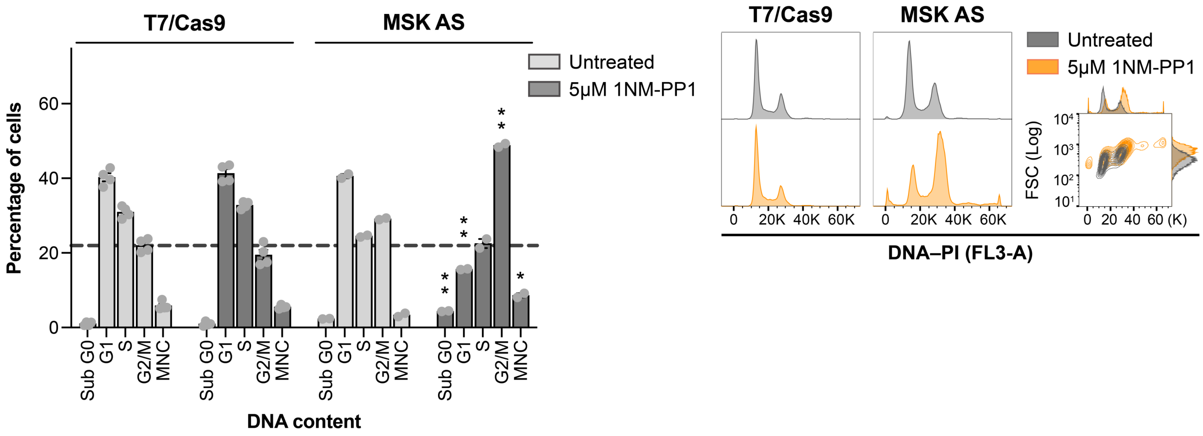

**c**

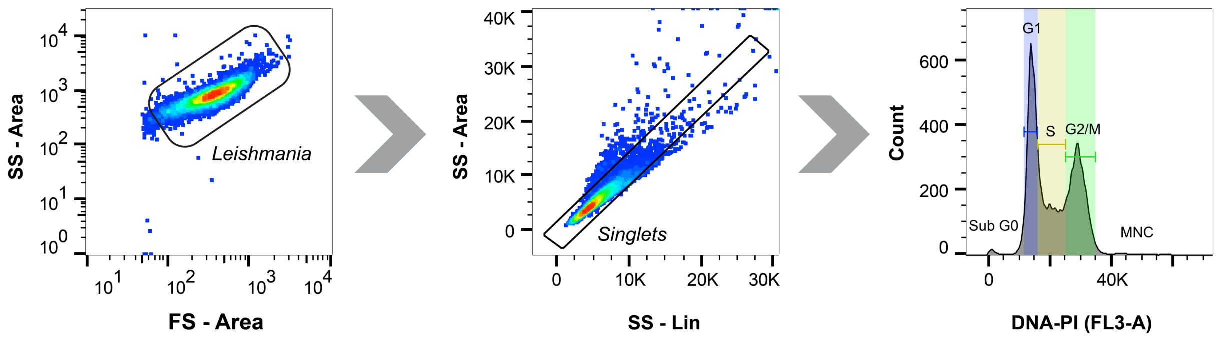

**Supplementary Fig. 7 Cell cycle-profile of MSK AS after 1NM-PP1 treatment. a** Cell cycle analysis of cells stained with propidium iodide (PI) after 6 h of treatment with 5µM of 1NM-PP1. Plot of the percentage of cells in each cell cycle stage. **b** Cell cycle analysis of cells stained with PI after 24 h the treatment course with 5µM of 1NM-PP1. Panel on the left, plot of the percentage of cell in each cell cycle stage. Panel on the right, representative cell cycle-profile histogram. The parental line T7/Cas9 and MSK AS untreated parasites cultured for the same time were used as controls. The dotted line represents the basal G2/M untreated average. MNC, multinucleated cells. P values were calculated using two-tailed Student's t-tests comparing the percentage of cells between treated and untreated. * p-value <0.1; ** p-value <0.01; *** p-value <0.001; **** p-value <0.0001. Data are mean ± SEM of four biological replicates (except for MSK AS treated with 1NM-PP1 during 24h, where only two biological replicates were analysed). **c** Representative gating strategy for cell cycle analysis. *Leishmania* cells were initially gated based on side scatter (SS) versus forward scatter (FS) area parameters to exclude debris. Single cells (*Singlets*) were then selected by applying a ~ 45° diagonal gate on a side scatter linear (SS - Lin) versus side scatter area (SS - Area) plot to remove doublets. The resulting population was analysed by plotting a DNA content histogram based on PI fluorescence. The FlowJo v.10.10.0 cell cycle algorithm Watson model was used to measure the percentage of cells in each cell cycle stage.

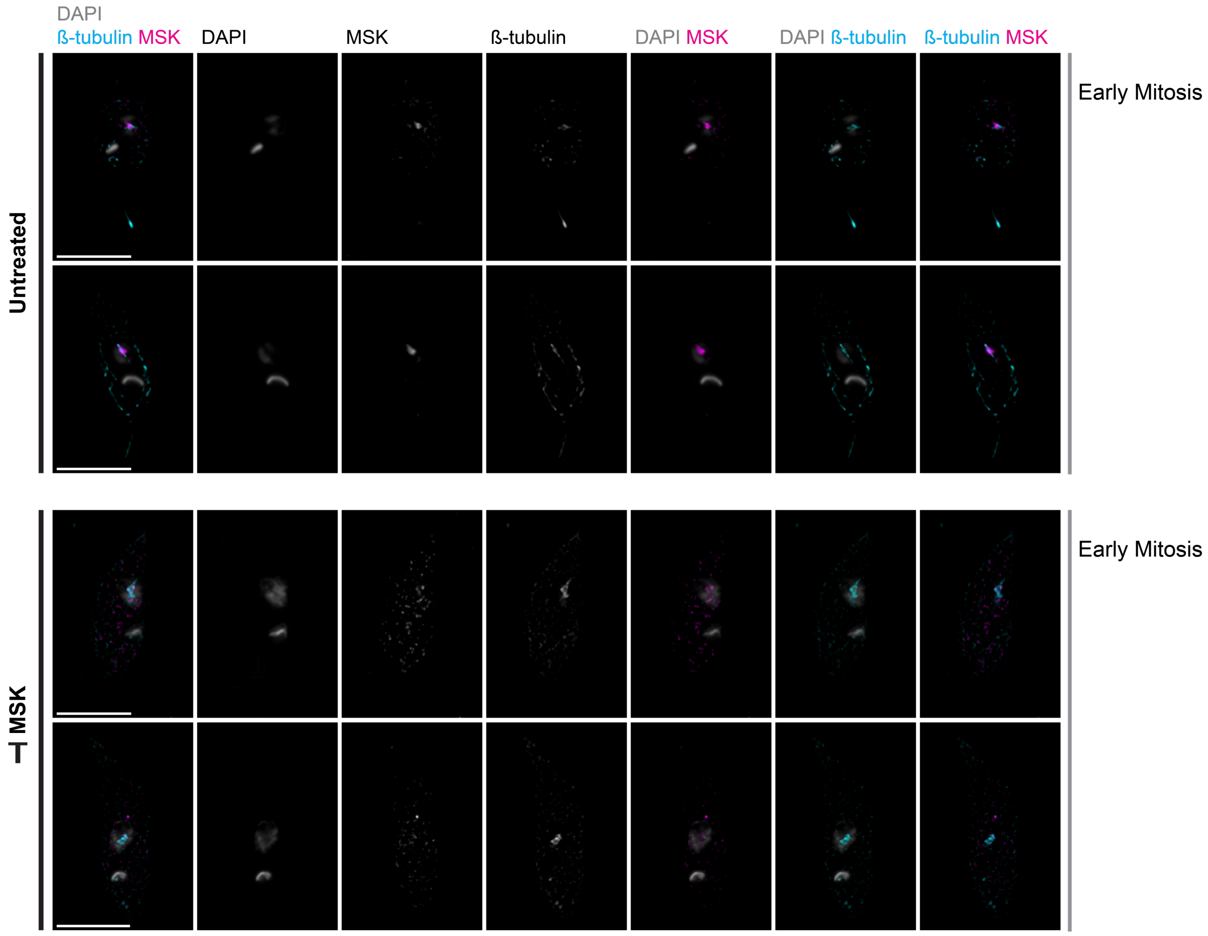

**Supplementary Fig. 8 High resolution microscopy of *L. mexicana* promastigotes endogenously expressing 3xMyc::mNG::MSK_AS.** High resolution microscopy of promastigotes, after 6 h treatment with 10 µM 1NM-PP1, stained with KMX-1 antibody to recognise β-tubulin, anti-Myc to recognise the endogenously tagged 3xMyc::mNG::MSK_AS, and counterstained with DAPI to visualise DNA. Representative fluorescence micrographs showing parasites in early mitosis in the presence and absence of the BKI. The channel and the colours used for each marking are indicated on the top. Scale bars, 5 µm.

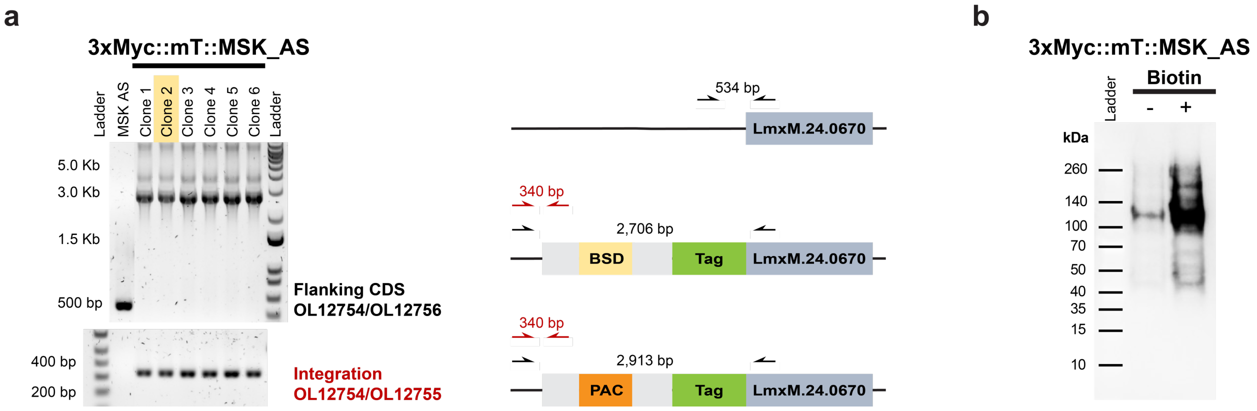

**Supplementary Fig. 9 Generation of cell lines expressing miniTurbo fused to MSK. a** Using CRISPR Cas9, the MSK AS line had both alleles of the MSK endogenously tagged with 3xMyc::mT to generate the 3xMyc::mT::MSK_AS cell lines. Repair templates containing blasticidin or puromycin drug resistance markers and linear DNA fragment for *in vivo* transcription of the single guide RNA were transfected into the parental line. After the overnight recovery time, transfected cells were selected with the blasticidin and puromycin, and then cloned by serial dilution. PCR amplification was used to determine the integration of the repair template in the correct position in both alleles. PCR strategies and DNA product size expected for each PCR are shown in the diagram on the right. OL, oligonucleotide as listed in Supplementary Table 3. The clone highlighted in yellow was selected to perform the further experiments. **b** Biotinylation of miniTurbo endogenously tagged MSK used for proximity labelling analysis. 3xMyc::mT::MSK_AS promastigotes were cultured with (+) or without (-) 500 µM biotin for 2h to facilitate biotinylation and then processed according to proximity labelling workflow to enrich biotinylated material. Enrichment of the biotinylated bait protein was assessed by western blotting using anti-Myc tag antibody. Predicted molecular weight for 3xMyc::mT::MSK is 173 kDa.

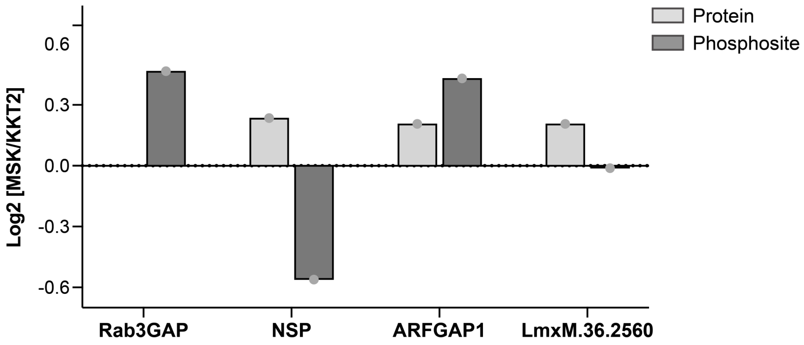

**Supplementary Fig. 10 Profile of the MSK substrates in the proteome and phosphoproteome MSK/KKT2 proximal maps.** Log2 fold change of the MSK substrates in the proteome and phosphoproteome comparing MSK and the spatial reference control KKT2. None of the protein or phosphosite were assigned as proximal to MSK or KKT2. p-values: 0.578896239 for the Rab3GAP phosphosite; 0.497028403 and 0.270212537 for the NSP protein and phosphosite, respectively; 0.732632028 and 0.228603919 for the ARFGAP1 protein and phosphosite, respectively; 0.905997756 and 0.981441948 for the hypothetical LmxM.36.2560 protein and phosphosite, respectively.

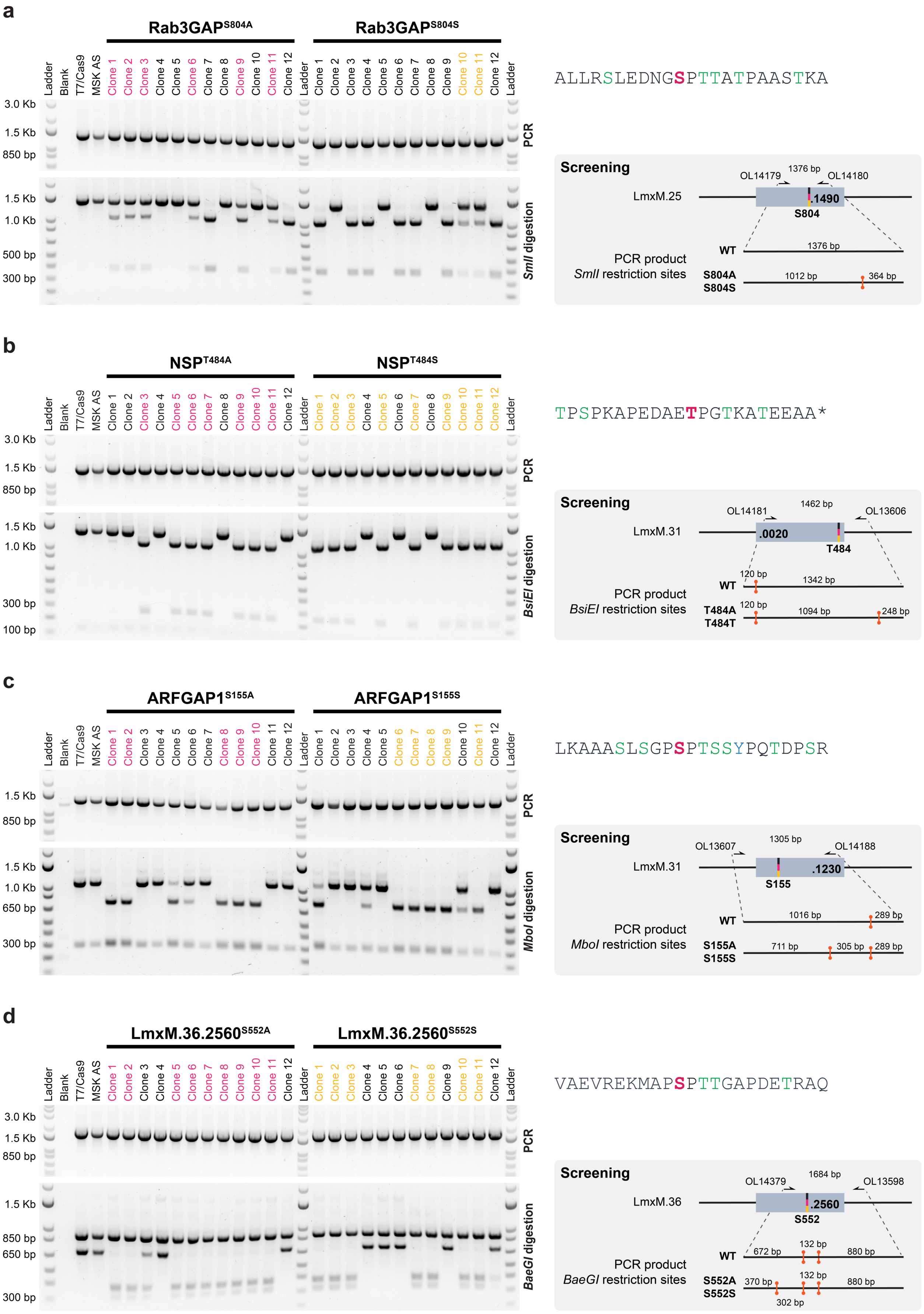

**Supplementary Fig. 11 Phosphosite mutants engineered by CRISPR-Cas9–mediated precision genome editing.** Using CRISPR Cas9 without drug resistance markers, the MSK substrates had both alleles of the phosphosite mutated to an Alanine in the MSK AS line. 200 bp double strand DNA, containing the codon mutation, was used as repair templates, which was transfected with the linear DNA fragment for *in vivo* transcription of the single guide RNA into the parental line. A repair template containing a silent mutation for the target phosphosite was used as control. After the overnight recovery time, transfected cells were cloned. PCR amplification followed by restriction digestion was used to genotype 12 clones from each transfection. The genotypes are highlighted as follow: Black, wild-type; Magenta, Rab3GAP$^{S804A}$ or NSP$^{T484A}$ or ARFGA1$^{S155A}$ or LmxM.36.2560$^{S552A}$; Yellow, silent mutation for Rab3GAP$^{S804S}$ or NSP$^{T484T}$ or ARFGA1$^{S155A}$ or LmxM.36.2560$^{S552S}$. Screening strategies used to genotype the target phosphosite are shown in the grey box on the right. Amino acid sequence surrounding the target phosphosite for each MSK interactor is shown above the screening box: magenta, target phosphosite; green other amino acid residue that can be phosphorylated. OL, oligonucleotide as listed in Supplementary Table 3. **a** Mutation of S804 in Rab3GAP (LmxM.25.1490). **b** Mutation of T484 in NSP (LmxM.31.0020). **c** Mutation of S115 in ARFGAP1 (LmxM.31.1230). **d** Mutation of S552 in LmxM.36.2560.

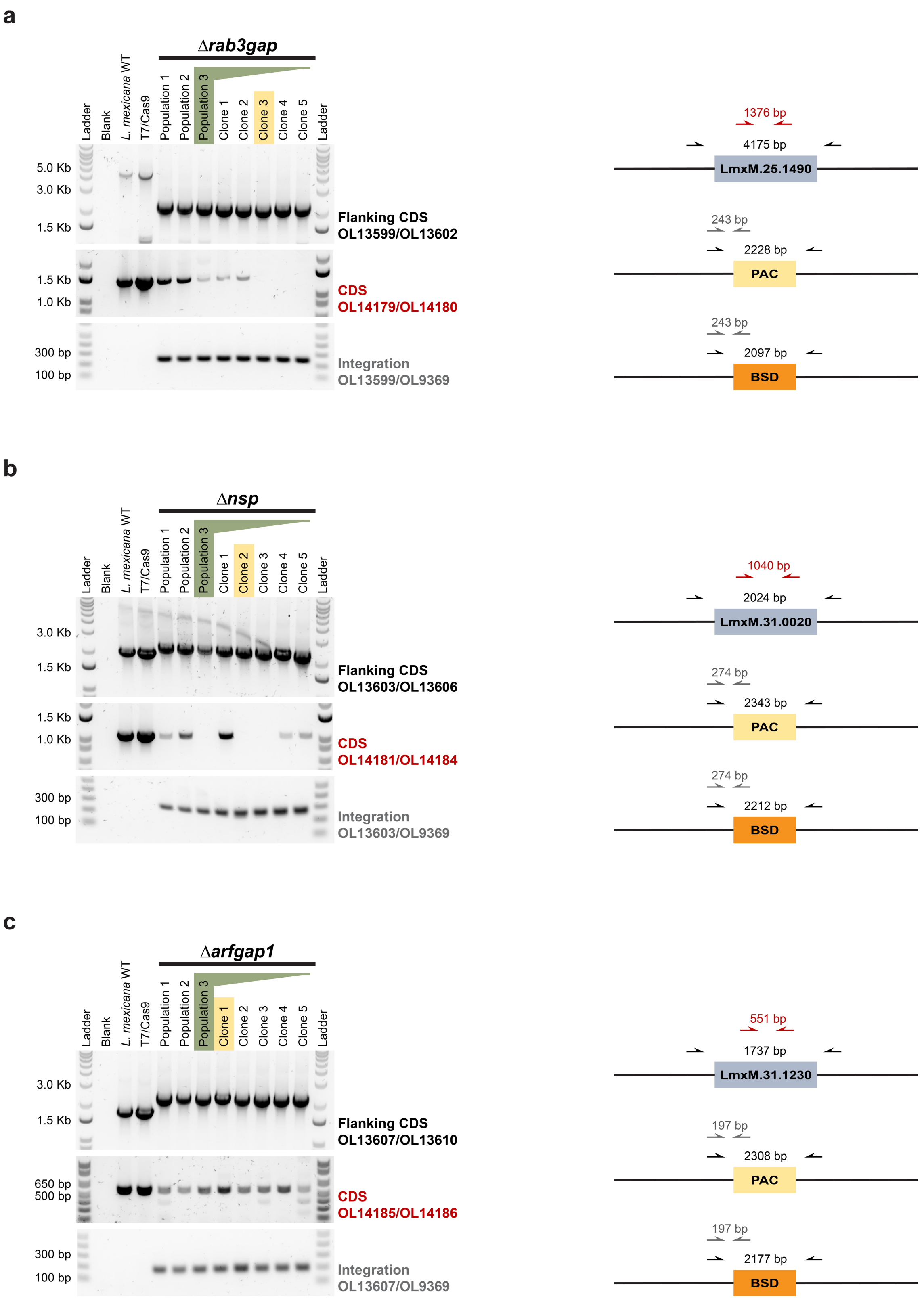

**Supplementary Fig. 12 Generation of Rab3GAP, NSP, ARFGAP1 and LmxM.36.2560 gene deletion mutants.**

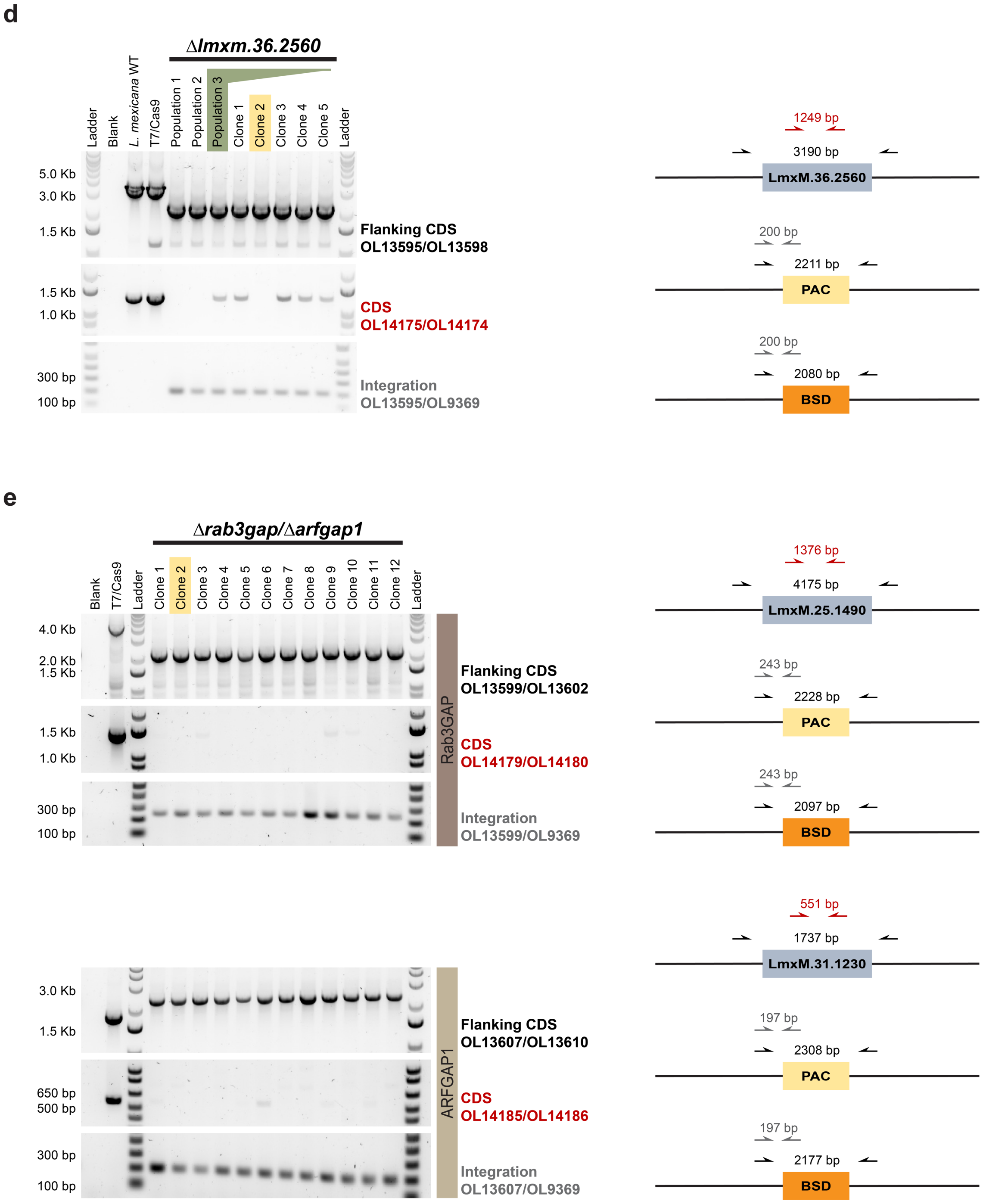

**Supplementary Fig. 12 Generation of *Rab3GAP*, *NSP*, *ARFGAP1* and *LmxM.36.2560* gene deletion mutants.** Three independent transfections were performed using DNA repair templates containing drug resistance markers (blasticidin or puromycin) and linear DNA fragment for *in vivo* transcription of the single guide RNA to individually delete the *Rab3GAP* (LmxM.25.1490), *NSP* (LmxM.31.0020), *ARFGAP1* (LmxM.31.1230) and the hypothetical *LmxM.2560* gene, or to simultaneously delete the two GTPase-activating proteins, Rab3GAP and ARFGAP1. All three transfections yielded populations capable of proliferating under drug selection. Clonal lines were derived from the third transfection. PCR amplification was used to assess the presence of the coding sequences (CDSs) and to confirm integration of the drug resistance cassettes. PCR strategies and DNA product size expected for each PCR are shown in the diagram on the right. OL, oligonucleotide as listed in Supplementary Table 3. The clones highlighted in yellow has their whole genome sequenced by Illumina. **a** Δ*Rab3gap*. **b** Δ*nsp*. **c** Δarfgap1. **d** Δ*LmxM.36.2560*. **e** Δ*Rab3gap*/*Δarfgp1*.

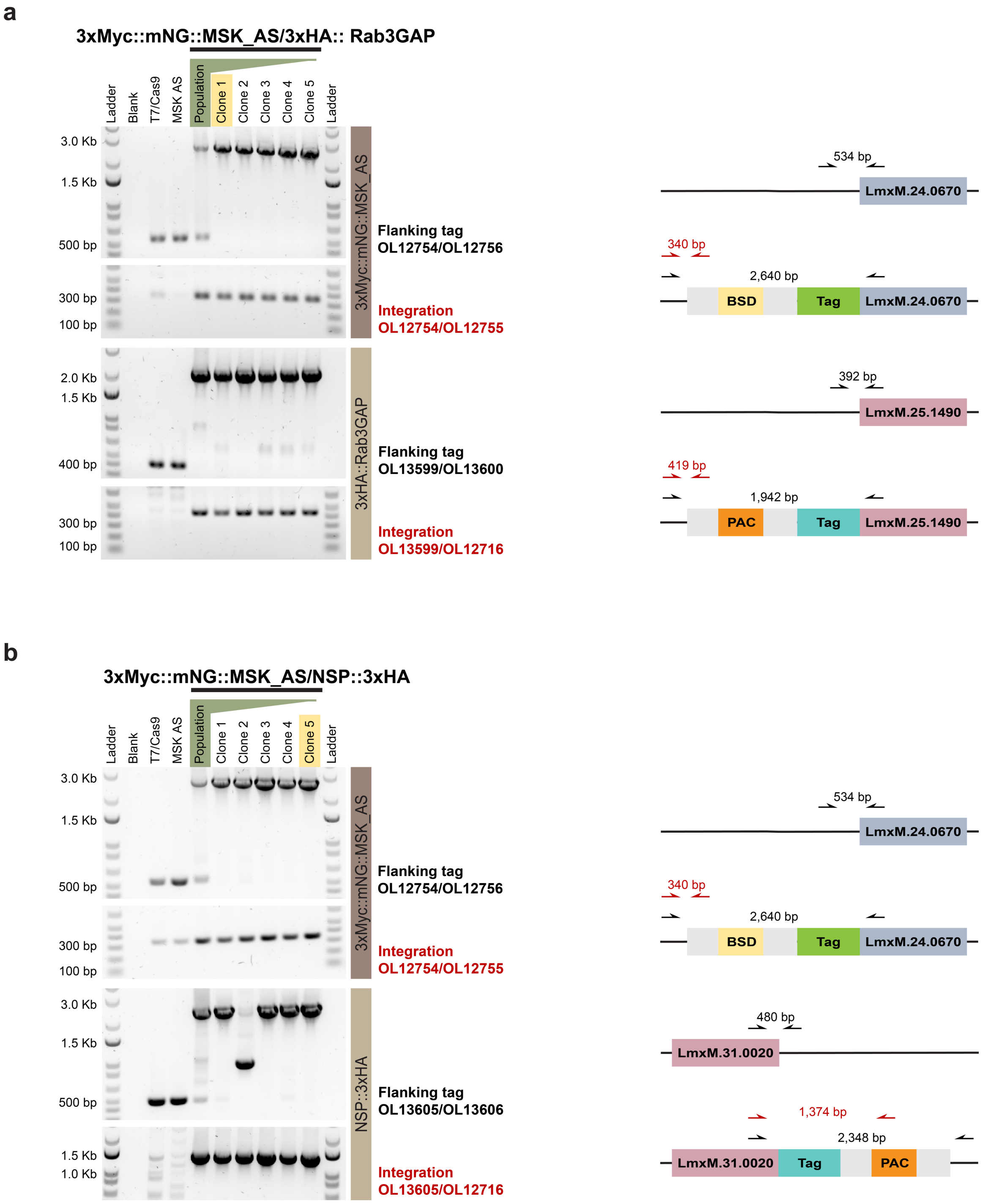

**Supplementary Fig. 13 Endogenous tagging of MSK and its substrates.**

**c**

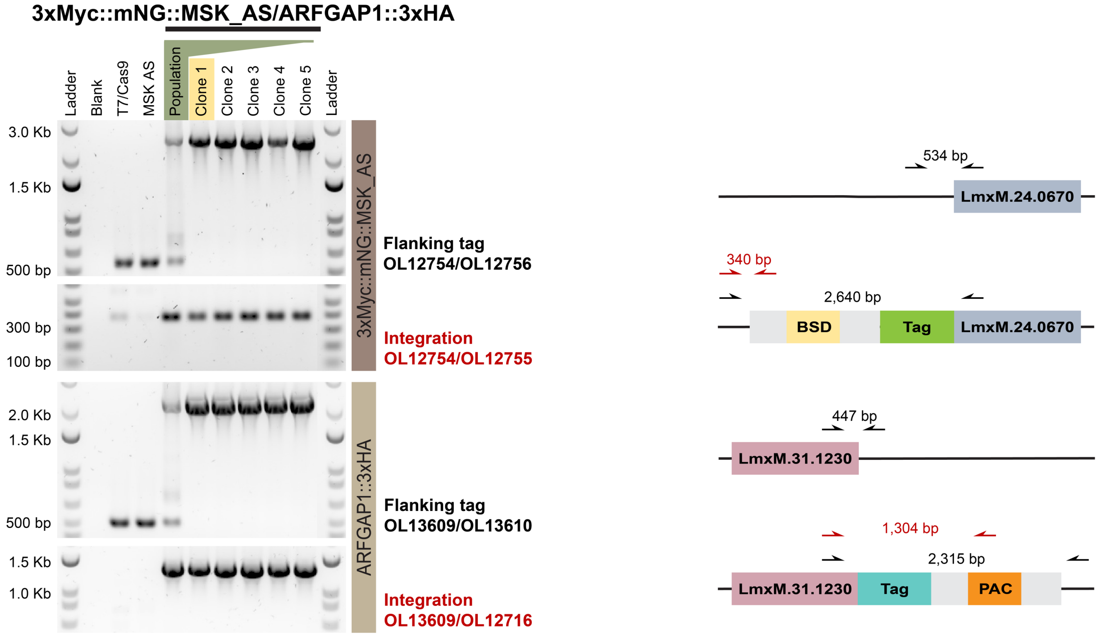

**d**

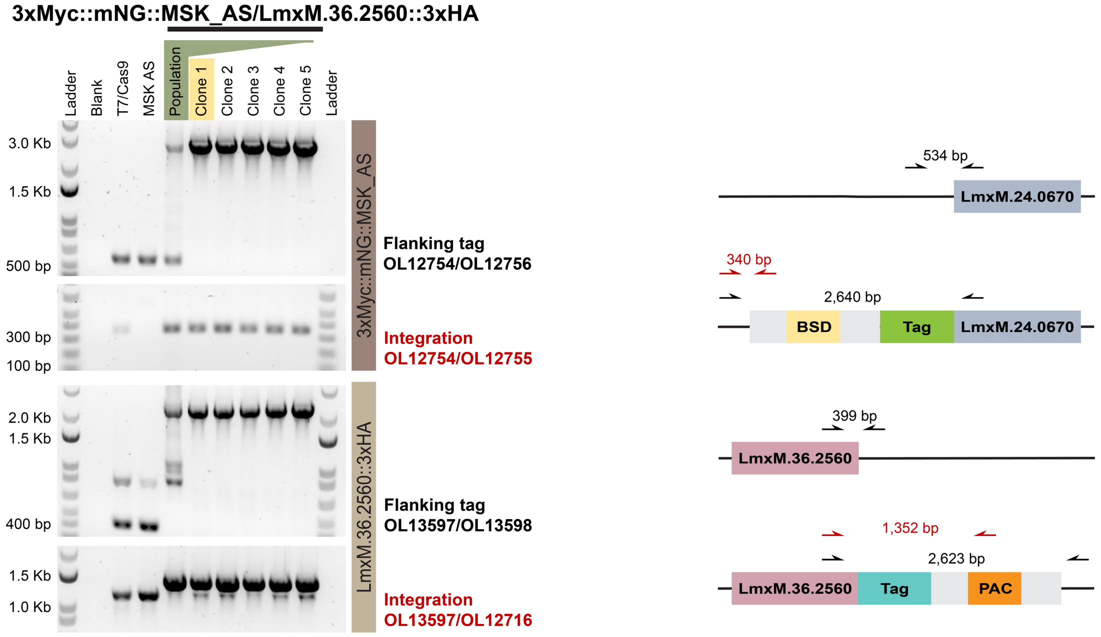

**Supplementary Fig. 13 Endogenous tagging of MSK and its substrates.** Using CRISPR-Cas9 genome editing, the MSK AS line was endogenously dual tagged: MSK was tagged with 3xMyc::mNeonGreen (mNG), and one of its substrates was simultaneously tagged with 3xHA. Repair templates containing blasticidin and puromycin as drug resistance marker, were used to tag MSK and its substrates, respectively. Together with their respectively linear DNA fragment for *in vivo* transcription of the single guide RNA, the repair templates were transfected into the parental line. After the overnight recovery time, transfected cells were selected with the blasticidin and puromycin, and then cloned by serial dilution. PCR amplification was used to confirm integration of the drug resistance cassettes. PCR strategies and DNA product size expected for each PCR are shown in the diagram on the right. OL, oligonucleotide as listed in Supplementary Table 3. The clones highlighted in yellow were selected to perform the further experiments. **a** 3xMyc::mNG::MSK_AS/3xHA:: Rab3GAP. **b** 3xMyc::mNG::MSK_AS/NSP::3xHA. **c** 3xMyc::mNG::MSK_AS/ARFGAP1::3xHA. **d** 3xMyc::mNG::MSK_AS/LmxM.36.2560::3xHA.

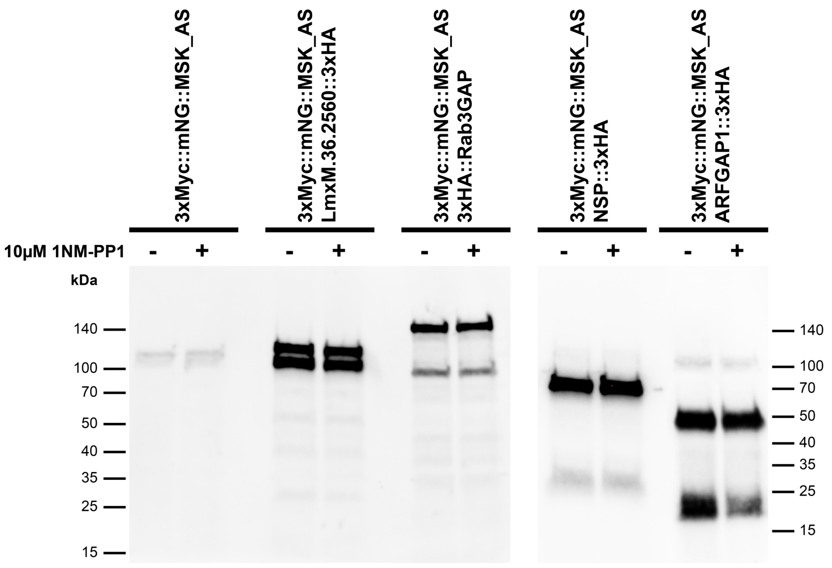

**Supplementary Fig. 14 Western blotting of MSK substrates.** Promastigote parasites were cultured with (+) or without (-) 10 µM 1NM-PP1 µM for 6 h. Predicted molecular weight for 3xMyc::mT::MSK is 171.24 kDa. Western blotting for the HA epitope, using mouse anti-HA antibody and anti-Mouse HRP conjugated antibody diluted at 1:5,000. Predicted molecular weight: 3xHA::Rab3GAP=138.82 kDa; NSP::3xHA=60.64 kDa; ARFGAP1::3xHA=47.09 kDa; LmxM.36.2560::3xHA=104.96 kDa.

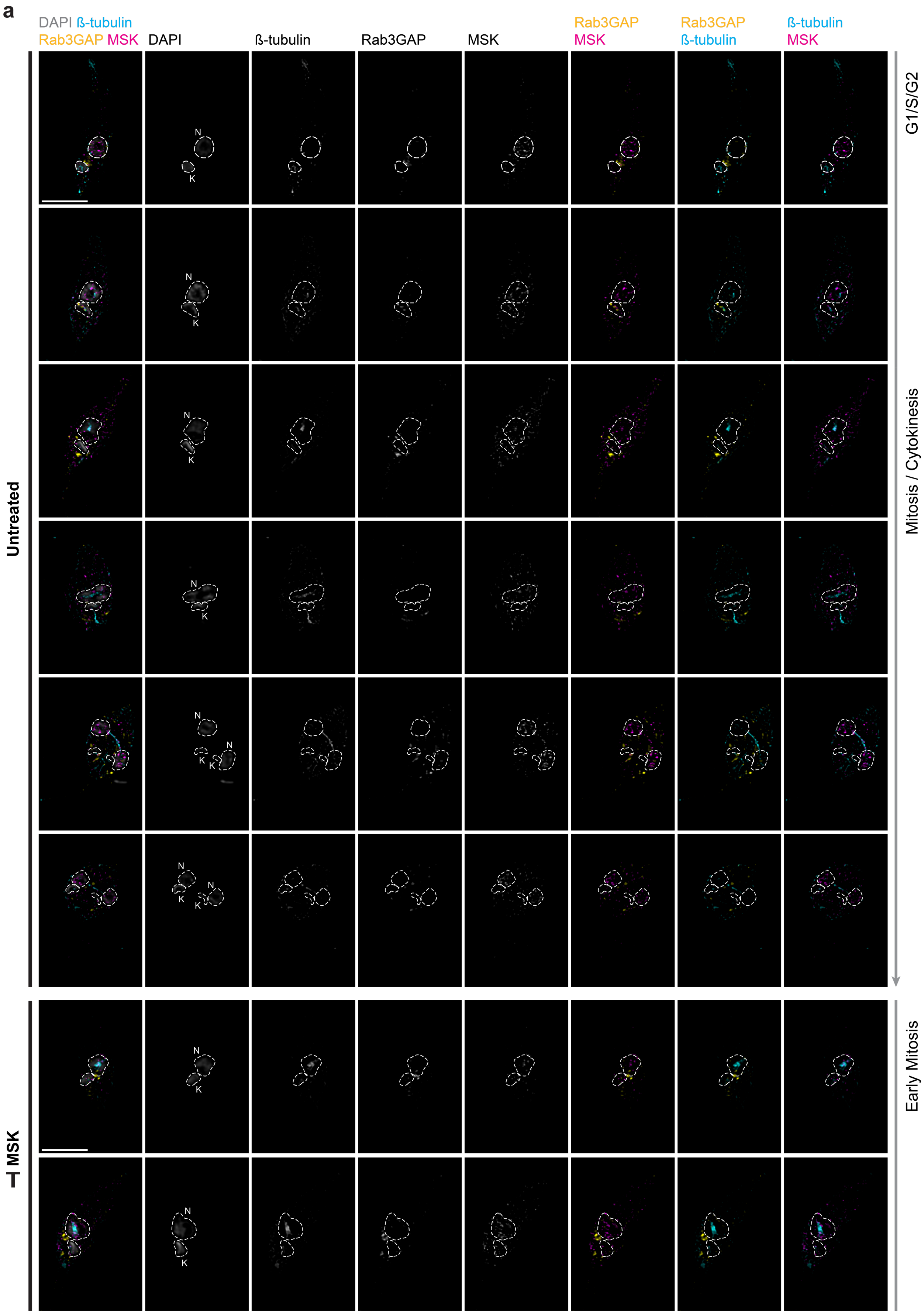

**Supplementary Fig. 15 The distribution of MSK and its substrates in *L. mexicana* promastigotes during the cell cycle.**

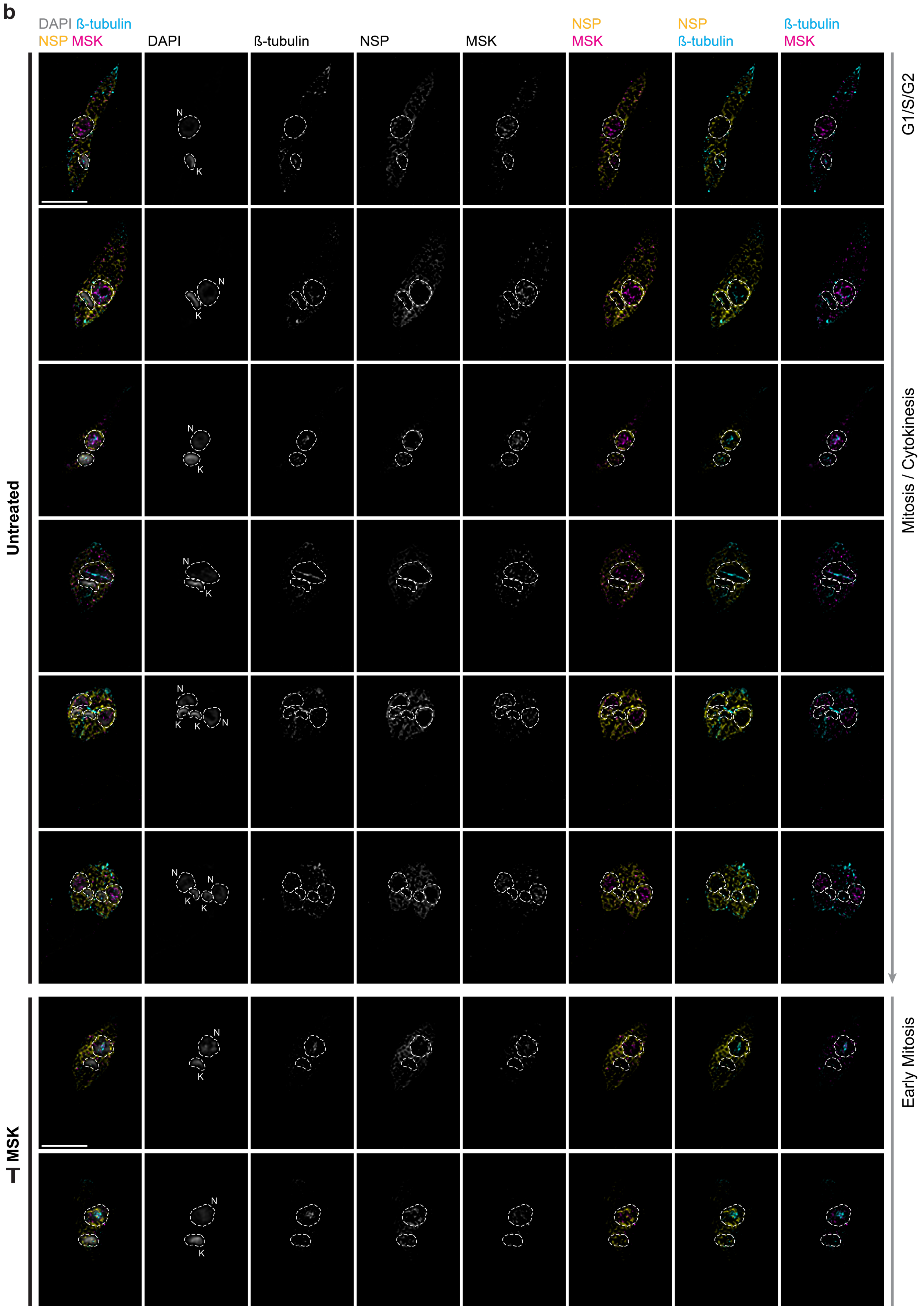

**Supplementary Fig. 15 The distribution of MSK and its substrates in *L. mexicana* promastigotes during the cell cycle.**

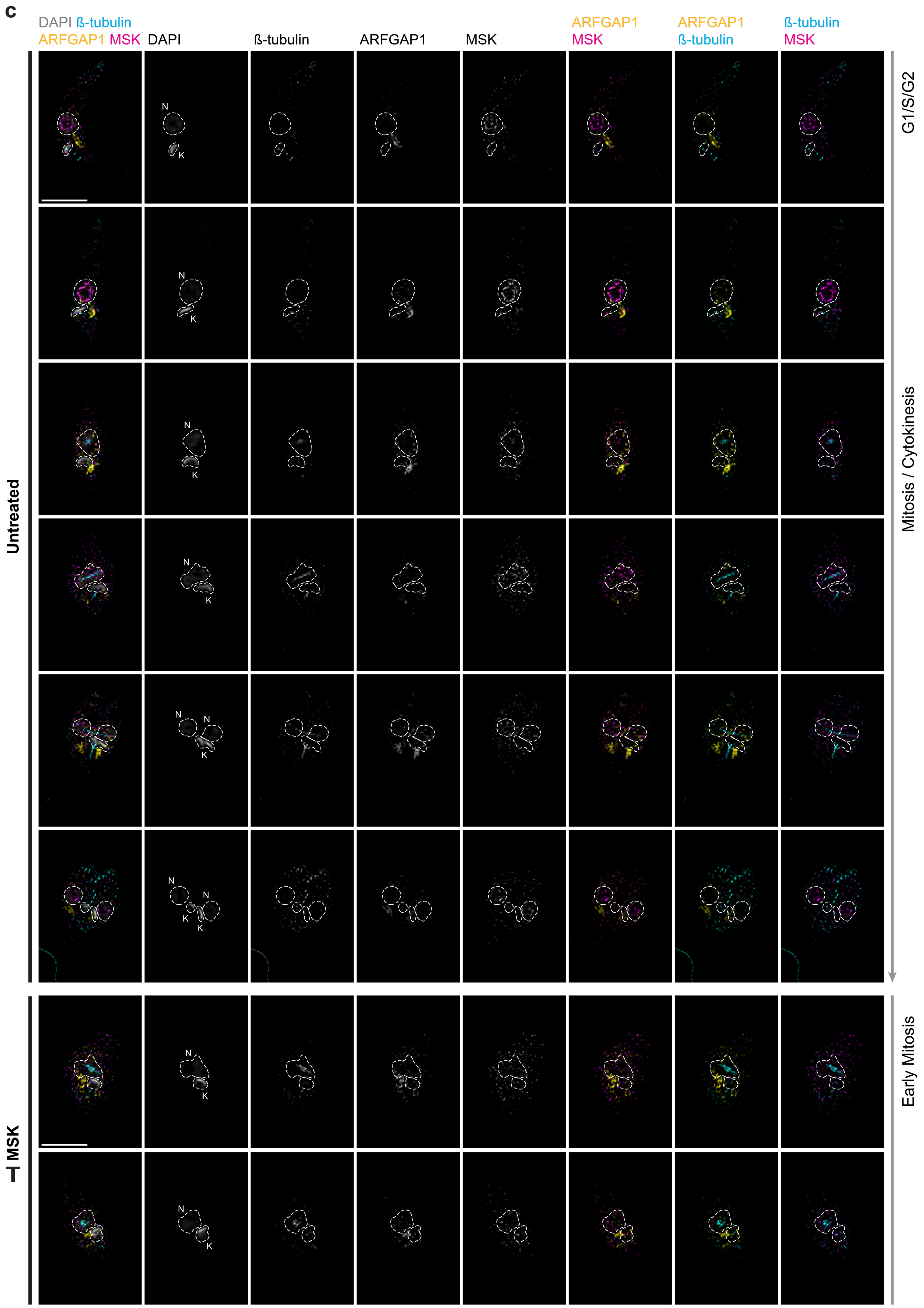

**Supplementary Fig. 15 The distribution of MSK and its substrates in *L. mexicana* promastigotes during the cell cycle.**

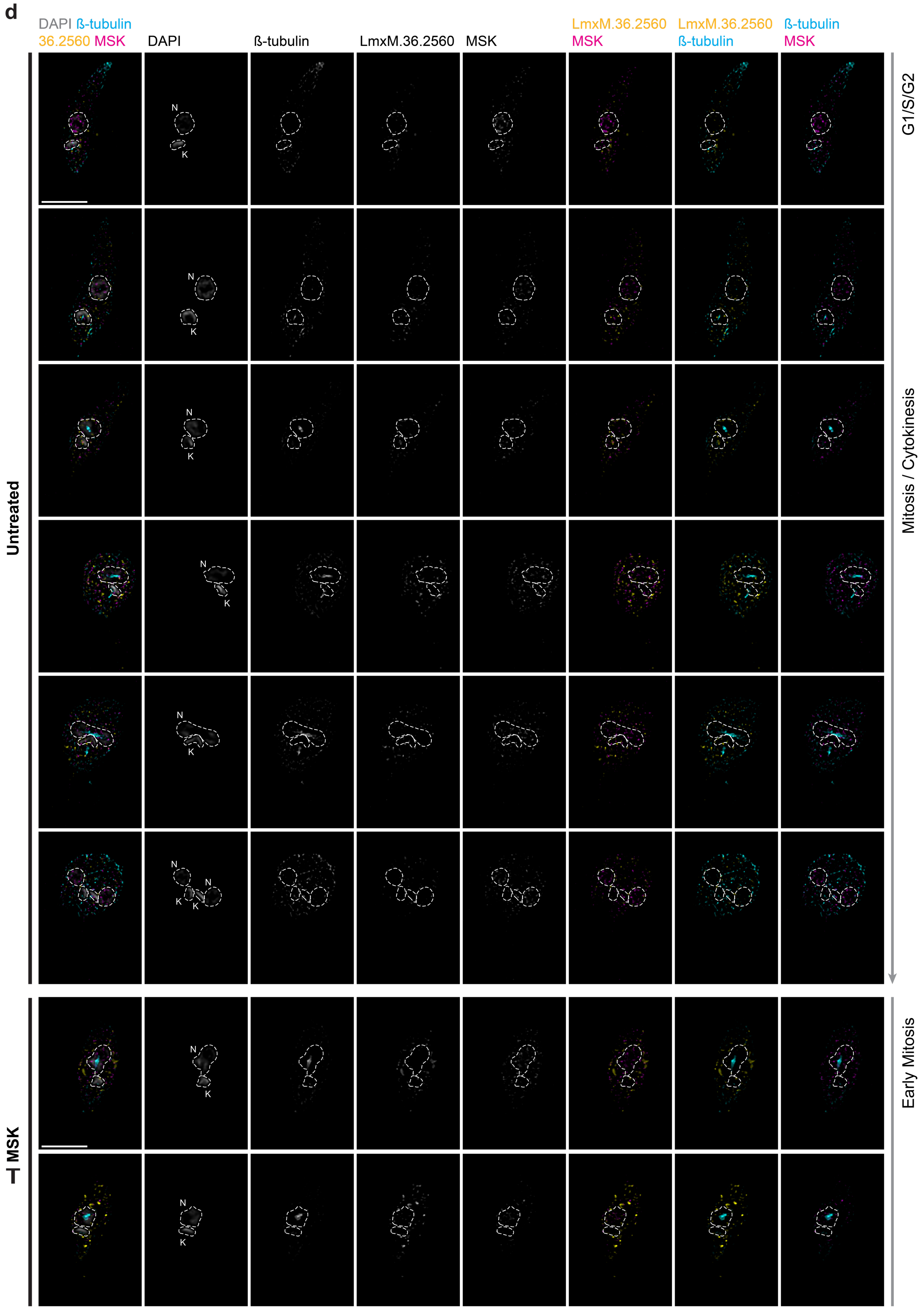

**Supplementary Fig. 15 The distribution of MSK and its substrates in *L. mexicana* promastigotes during the cell cycle.** High resolution microscopy of promastigotes after 6 h of treatment with or without 10 µM 1NM-PP1, stained with KMX-1 antibody to recognise β-tubulin, anti-Myc to recognise the endogenously tagged 3xMyc::mNG::MSK_AS, anti-HA to recognise MSK substrates endogenously tagged with 3xHA (3xHA::Rab3GAP; or NSP::3xHA; or 3xHA::ARFGAP1; or 3xHA::LmxM.36.2560) and counterstained with DAPI to visualise cell DNA. Representative fluorescence micrographs showing parasites in different cell cycle stages. The channel and the colours used for each marking are indicated on the top. Scale bars, 5 µm. **a** Microscopy of MSK and Rab3GAP proteins. **b** Microscopy of MSK and NSP proteins. **c** Microscopy of MSK and ARFGAP1 proteins. **d** Microscopy of MSK and the hypothetical protein LmxM.36.2560.

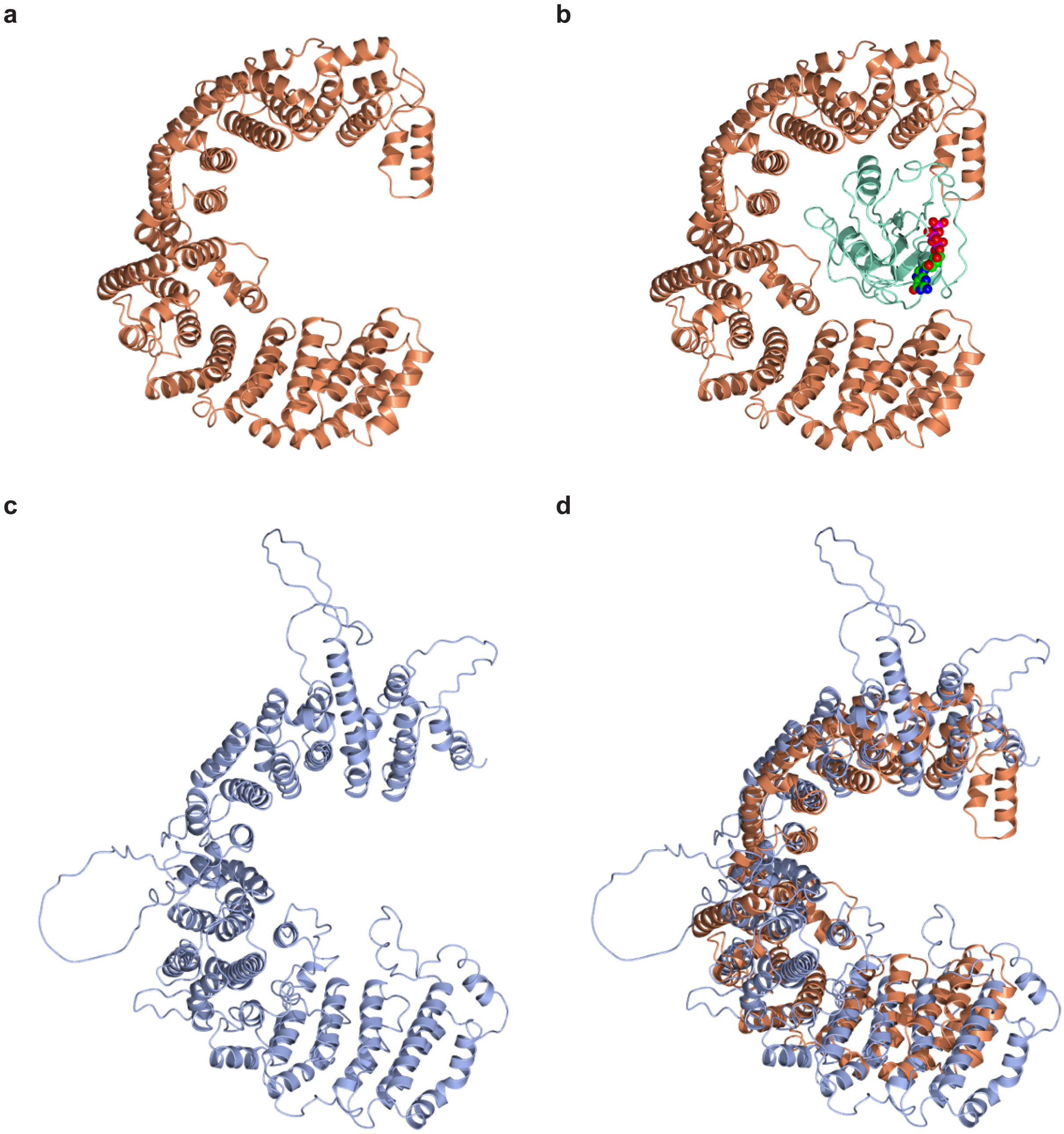

**Supplementary Fig. 16 Structural homology between the *Leishmania mexicana* hypothetical protein LmxM.36.2560 and yeast importin Kap114. a-b** Cartoon representation of yeast importin Kap114 (PDB code 8F7A, N-terminal domain, colored coral) in complex with Ran (green) bound to GTP (spheres). **c** AlphaFold 3 model of LmxM.36.2560 (blue). **d** Structural superposition of LmxM.36.2560 and Kap114. The proteins share an overall architecture. Moreover the surface of the horseshoe interior is rich in Asp/Glu residues to give an electronegative environment which is a common feature of importins.

### References

1 Fiebig, M., Kelly, S. & Gluenz, E. Comparative Life Cycle Transcriptomics Revises Leishmania mexicana Genome Annotation and Links a Chromosome Duplication with Parasitism of Vertebrates. *PLoS Pathog* **11**, e1005186 (2015). https://doi.org/10.1371/journal.ppat.1005186
