## Supplementary Data 3 for "The mitotic spindle kinase MSK co-ordinates segregation of the nucleus and kinetoplast in *Leishmania mexicana*"

**Code Availability**

**CUSTOM R CODE TO ANALYSE PROXIMITY LABELLING EXPERIMENT**

**1. ‘Total’ proximal protein**

**1.1. Code to analyse MSK against the spatial reference KKT2**

```
library('ggplot2')
library('dplyr')
library('reshape2')
library('RColorBrewer')
library('coin')
library('ggrepel')
library('limma')
library('statmod')
library('grid')
library('gridExtra')
set.seed(42)
options(ggrepel.max.overlaps = Inf)


#-----------------------------Extracting dataset that will be compared---------------------------

total <- read.csv('D548_QI_Total_Proteome_Results.csv', header=TRUE, skip=4)
head(total)
nrow(total)

#extracting the selected data (only the groups that will be compared)
total_s <- (total[,c('MSK_A', 'MSK_B', 'MSK_C', 'MSK_D', 'MSK_E',
                     'KKT2_A', 'KKT2_B', 'KKT2_C', 'KKT2_D')])

rownames(total_s) <- total$Accession
head(total_s)
class(total_s)

write.csv(total_s, 'D548_QI_Total_Proteome_Results_Sel.csv')

#Proteins with ≥2 missing values across the five biological replicates were removed
#Use =COUNTIF (range, "criterion") in excel to retain proteins with non-zero intensity in at least 3 of the 5 biological replicates.
#Save file as 'D548_QI_Total_Proteome_Results_Sel_n0.csv'


#--------------------------------------Imputing 0 values-----------------------------------------
total <- read.csv('D548_QI_Total_Proteome_Results_Sel_n0.csv', header=TRUE)
head(total)
nrow(total)

total_m <- melt(total[,c('MSK_A', 'MSK_B', 'MSK_C', 'MSK_D', 'MSK_E',
                     'KKT2_A', 'KKT2_B', 'KKT2_C', 'KKT2_D')])

total_m$Sample_Group <- sapply(strsplit(as.character(total_m$variable), "(?<=_)[12345]", perl=TRUE), function(x) x[[1]])

sd_intensities <- sd(log2(total_m$value[which(total_m$value>0)]))
sd_intensities#3.93
mean_intensities <- mean(log2(total_m$value[which(total_m$value>0)]))
mean_intensities#16.06

n5000 <- data.frame(value=(rnorm(500, mean=6.5, sd=1.5)),variable='norm')
head(n5000)

p <- ggplot(total_m, aes(x=log2(value), fill=Sample_Group))+
  geom_histogram(binwidth=0.3)+
  geom_histogram(data=n5000,aes(x=value), fill='yellow', alpha=0.5,binwidth=0.3)+
  #geom_density()
  facet_wrap(~Sample_Group, scales='free_y')+xlab('log2 protein intensity')

p

#set seed for generating random numbers from rnorm
#value are for peptide level imputation
set.seed(2021-06-01)
impute <- function(x){
  #make sure input is numeric vector
  b <- as.numeric(x)
  #count zeros in group
  n.zeros <- length(b[b==0])
  if(n.zeros > 0){
    ndx.zeros <- which(b %in% c(0))
    #generate random number from normal distribution which
    #models low abundance peptides
    imputed_value <- rnorm(n.zeros,mean=6.5,sd=1.5)
    #transform imputed values from log space
    imputed_values_intensity <- 2^(imputed_value)
    b[ndx.zeros] <- imputed_values_intensity
    return(c(b))
  }else{
    return(c(b))
  }
}

#impute missing values per sample group randomly taking number from normal distribution created
#model the low intensity peptides
total_imputed <- apply(total[,c('MSK_A', 'MSK_B', 'MSK_C', 'MSK_D', 'MSK_E',
                     'KKT2_A', 'KKT2_B', 'KKT2_C', 'KKT2_D')],1 ,function(x) impute(x))

#apply returns vector, so need to convert to matrix, filling by row
total_imputed <- matrix(total_imputed, ncol=9, byrow=TRUE)
colnames(total_imputed) <- c('MSK_A', 'MSK_B', 'MSK_C', 'MSK_D', 'MSK_E',
                     'KKT2_A', 'KKT2_B', 'KKT2_C', 'KKT2_D')

rownames(total_imputed) <- total$Accession
head(total_imputed)
nrow(total_imputed)#3570
write.csv(total_imputed, 'Total_imputed_zero_replaced.csv')


#------------------------------------Statistical analysis----------------------------------------
#limma analysis
#Before run the next command label column A as "Accession"

design <- read.csv('limma_input_Total_MSKvsKKT2_model.csv', row.names=1, header=TRUE)
design <- as.matrix(design)
head(design)

contrasts <- read.csv('limma_input_Total_MSKvsKKT2_contrasts.csv', header=TRUE, row.names=1)
head(contrasts)

#get data for this comparison
total_MSK_vs_KKT2 <- as.matrix(total_imputed[,c('MSK_A’, 'MSK_B', 'MSK_C', 'MSK_D', 'MSK_E',
                     			'KKT2_A', 'KKT2_B', 'KKT2_C', 'KKT2_D')])

rownames(total_MSK_vs_KKT2) <- total$Accession

total_MSK_vs_KKT2<- na.omit(total_MSK_vs_KKT2)
total_MSK_vs_KKT2<- apply(total_MSK_vs_KKT2, c(1,2), function(x) log2(x))

#try eBayes options#seems to give good results
fit <- lmFit(total_MSK_vs_KKT2, design)
fit_contrast <- contrasts.fit(fit, contrasts)
fit_contrast <- eBayes(fit_contrast, trend=TRUE, robust=TRUE)
#fit_contrast <- eBayes(fit_contrast, robust=TRUE)
table_ total_MSK_vs_KKT2<- topTable(fit_contrast, number=Inf)
head(table_ total_MSK_vs_KKT2)
nrow(table_ total_MSK_vs_KKT2)#3570
write.csv(table_ total_MSK_vs_KKT2, 'limma_output_ total_MSK_vs_KKT2.csv')

#adjust p values with benjamini hochberg
t_test_results <- table_ total_MSK_vs_KKT2[,c('P.Value')]
ndx <- order(t_test_results)
ttest_order <- t_test_results[ndx]
length(ttest_order)
length(ttest_order[!is.na(ttest_order)])
ttest_BH <- data.frame(pvalues=ttest_order, rank=rep(1:3570))
#set FDR to 5%
ttest_BH$BH_critical <- (ttest_BH$rank/3570)*0.05
tail(ttest_BH)
cutoff <- ttest_BH[which(ttest_BH$pvalues < ttest_BH$BH_critical),]
head(cutoff)
tail(cutoff)
#5% FDR: 143 proteins


#-------------------------------------------Radial Plot------------------------------------------
#As the analysis was performed with KKT2 against MSK, the log2FC (KKT2/MSK) was transformed to lof2FC (MSK/KKT2), using Microsoft Excel.

#Data set
data_total <- read.csv("Total_MSK.proximals_vs_KKT2.csv", header=TRUE, row.names=1)
head(data_total)

proximals <- data_total[which(data_total$adj.P.Val <= 0.05 & data_total$log2FC > 0),]
nrow(proximals)#78 proximal proteins for MSK

to_plot <- data_total[,c('log2FC','adj.P.Val','Radial.plot.label','Protein.function.process’)]
to_plot_sig <- to_plot[which(to_plot$adj.P.Val <0.05),]
nrow(to_plot_sig)#143 significantly changing proteins
to_plot_sig
head(to_plot)

display.brewer.all()
colours <- brewer.pal(n=12, name='Paired')
display.brewer.pal(n = 12, name = 'Paired')
brewer.pal(n = 12, name = 'Paired')
colours

#select colours to represent the highlights
cols <- c('Protein kinase'=colours[4],
          'Phosphatase'=colours[7],
          'Kinetochore'=colours[10],
          'Histone'=colours[8],
          'RNA binding / Processing / Translation'=colours[6],
          'DNA binding / Processing / Repair / Replication'=colours[1],
          'Protein Folding'=colours[9],
          'Microtubule; Proteins that interact with microtubules; Cell motility'=colours[2],
          'Nuclear pore'=colours[3],
          'Nucleotide Metabolism / Binding'=colours[11],
          'Hypothetical'=colours[5],
          'Other'=colours[12])

p <- ggplot(to_plot[which(to_plot$log2FC > 0),],
            aes(x=log2FC, y=log10(adj.P.Val),
                color= Protein.function.process,
                label=ifelse(adj.P.Val <= 0.05, as.character(paste(Radial.plot.label), size=4),'')))+

  geom_point(colour='grey40', size=4, alpha=0.5)+
  scale_color_manual(values=cols)+

  geom_point(data=to_plot[which(to_plot$log2FC>0 & to_plot$adj.P.Val <0.05),],
             aes(x=log2FC, y=log10(adj.P.Val),
                 color=Protein.function.process), size=4, alpha=0.5)+

  geom_text_repel(data=to_plot[which(to_plot$adj.P.Val < 0.05),], aes(label=Radial.plot.label), size=4,
                  min.segment.length = 0,
                  seed = 42,
                  box.padding = unit(0.5, "lines"),
                  point.padding = unit(0, "lines"),
                  nudge_y = 3.5,
                  segment.alpha = 0.5,
                  force = 200,
                  force_pull = 50,
                  max.time = 1000,
                  show.legend = FALSE,
                  max.overlaps = Inf
  ) +

  coord_polar(clip = 'off')+

  scale_x_continuous(
    limits=c(0, 7.5),
    breaks = c(0, 1.5, 3, 4.5, 6))+

  scale_y_continuous(
    limits = c(-5, 0),
    breaks = c(-2, -3, -4, -5)) +

  #adding the cut-off line for p-value <0.05
  geom_hline(yintercept = -1.301029995663981, colour="grey30", size=0.5, linetype='dashed')+

  annotate(
    x = 6.1,
    y = -1.7,
    label = "-2",
    geom = "text",
    color = "gray12",
    size = 3,
  ) +
  annotate(
    x = 6.1,
    y = -2.7,
    label = "-3",
    geom = "text",
    color = "gray12",
    size = 3,
  ) +
  annotate(
    x = 6.15,
    y = -3.7,
    label = "-4",
    geom = "text",
    color = "gray12",
    size = 3,
  ) +
  annotate(
    x = 6.25,
    y =-4.7,
    label = "-5",
    geom = "text",
    color = "gray12",
    size = 3,
  ) +

  #removing the original label axis and axis
  labs(y = NULL)+
  labs(x = NULL)+

  #selecting the graph theme
  theme_bw(base_size = 14)+
  theme(
    axis.text.y = element_blank(),
    axis.ticks = element_blank(),
    panel.border = element_blank(),
  )

p

ggsave('Total_Radial Total_MSK_Proximal_vs_KKT2', dpi=1000)
ggsave('Total_Radial Total_MSK_Proximal_vs_KKT2.pdf', p, width=13, height=6)
```

**1.2. Code to analyse MSK against inhibited MSK**

```
library('ggplot2')
library('dplyr')
library('reshape2')
library('RColorBrewer')
library('coin')
library('ggrepel')
library('limma')
library('statmod')
library('grid')
library('gridExtra')
set.seed(42)
options(ggrepel.max.overlaps = Inf)


#-----------------------------Extracting dataset that will be compared---------------------------

total <- read.csv('D548_QI_Total_Proteome_Results.csv', header=TRUE, skip=4)
head(total)
nrow(total)

#extracting the selected data (only the groups that will be compared)
total_s <- (total[,c('MSK_A', 'MSK_B', 'MSK_C', 'MSK_D', 'MSK_E',
                     'MSK_BKI_A', 'MSK_BKI_B', 'MSK_BKI_C', 'MSK_BKI_D’, 'MSK_BKI_E')])

rownames(total_s) <- total$Accession
head(total_s)
class(total_s)

write.csv(total_s, 'D548_QI_Total_Proteome_Results_Sel.csv')

#Removing protein with ≥2 missing values across the five biological replicates were removed
#Use =COUNTIF (range, "criterion") in excel to retain proteins with non-zero intensity in at least 3 of the 5 biological replicates.
#Save file as 'D548_QI_Total_Proteome_Results_Sel_n0.csv'


#--------------------------------------Imputing 0 values-----------------------------------------
total <- read.csv('D548_QI_Total_Proteome_Results_Sel_n0.csv', header=TRUE)
head(total)
nrow(total)

total_m <- melt(total[,c('MSK_A', 'MSK_B', 'MSK_C', 'MSK_D', 'MSK_E',
                     'MSK_BKI_A', 'MSK_BKI_B', 'MSK_BKI_C', 'MSK_BKI_D’, 'MSK_BKI_E')])

total_m$Sample_Group <- sapply(strsplit(as.character(total_m$variable), "(?<=_)[12345]", perl=TRUE), function(x) x[[1]])

sd_intensities <- sd(log2(total_m$value[which(total_m$value>0)]))
sd_intensities#3.99
mean_intensities <- mean(log2(total_m$value[which(total_m$value>0)]))
mean_intensities#16.02

n5000 <- data.frame(value=(rnorm(500, mean=6.5, sd=1.5)),variable='norm')
head(n5000)

p <- ggplot(total_m, aes(x=log2(value), fill=Sample_Group))+
  geom_histogram(binwidth=0.3)+
  geom_histogram(data=n5000,aes(x=value), fill='yellow', alpha=0.5,binwidth=0.3)+
  #geom_density()
  facet_wrap(~Sample_Group, scales='free_y')+xlab('log2 protein intensity')

p

#set seed for generating random numbers from rnorm
#value are for peptide level imputation
set.seed(2021-06-01)
impute <- function(x){
  #make sure input is numeric vector
  b <- as.numeric(x)
  #count zeros in group
  n.zeros <- length(b[b==0])
  if(n.zeros > 0){
    ndx.zeros <- which(b %in% c(0))
    #generate random number from normal distribution which
    #models low abundance peptides
    imputed_value <- rnorm(n.zeros,mean=6.5,sd=1.5)
    #transform imputed values from log space
    imputed_values_intensity <- 2^(imputed_value)
    b[ndx.zeros] <- imputed_values_intensity
    return(c(b))
  }else{
    return(c(b))
  }
}

#impute missing values per sample group randomly taking number from normal distribution created
#model the low intensity peptides
total_imputed <- apply(total[,c('MSK_A', 'MSK_B', 'MSK_C', 'MSK_D', 'MSK_E',
   'MSK_BKI_A', 'MSK_BKI_B', 'MSK_BKI_C', 'MSK_BKI_D’, 'MSK_BKI_E')],1 ,function(x) impute(x))

#apply returns vector, so need to convert to matrix, filling by row
total_imputed <- matrix(total_imputed, ncol=9, byrow=TRUE)
colnames(total_imputed) <- c('MSK_A', 'MSK_B', 'MSK_C', 'MSK_D', 'MSK_E',
                     'MSK_BKI_A', 'MSK_BKI_B', 'MSK_BKI_C', 'MSK_BKI_D’, 'MSK_BKI_E')

rownames(total_imputed) <- total$Accession
head(total_imputed)
nrow(total_imputed)#3541
write.csv(total_imputed, 'Total_imputed_zero_replaced.csv')


#------------------------------------Statistical analysis----------------------------------------
#limma analysis
#Before run the next command label column A as "Accession"

design <- read.csv('limma_input_Total_MSKvsMSK_BKI_model.csv', row.names=1, header=TRUE)
design <- as.matrix(design)
head(design)

contrasts <- read.csv('limma_input_Total_MSKvsMSK_BKI_contrasts.csv', header=TRUE, row.names=1)
head(contrasts)

#get data for this comparison
total_MSK_vs_MSK_BKI <- as.matrix(total_imputed[,c('MSK_A', 'MSK_B', 'MSK_C', 'MSK_D', 'MSK_E',
                     'MSK_BKI_A', 'MSK_BKI_B', 'MSK_BKI_C', 'MSK_BKI_D’, 'MSK_BKI_E')])

rownames(total_MSK_vs_MSK_BKI) <- total$Accession

total_MSK_vs_MSK_BKI<- na.omit(total_MSK_vs_MSK_BKI)
total_MSK_vs_MSK_BKI<- apply(total_MSK_vs_MSK_BKI, c(1,2), function(x) log2(x))

#try eBayes options#seems to give good results
fit <- lmFit(total_MSK_vs_MSK_BKI, design)
fit_contrast <- contrasts.fit(fit, contrasts)
fit_contrast <- eBayes(fit_contrast, trend=TRUE, robust=TRUE)
#fit_contrast <- eBayes(fit_contrast, robust=TRUE)
table_ total_MSK_vs_MSK_BKI<- topTable(fit_contrast, number=Inf)
head(table_ total_MSK_vs_MSK_BKI)
nrow(table_ total_MSK_vs_MSK_BKI)#3541
write.csv(table_ total_MSK_vs_MSK_BKI, 'limma_output_ total_MSK_vs_MSK_BKI.csv')

#adjust p values with benjamini hochberg
t_test_results <- table_ total_MSK_vs_MSK_BKI[,c('P.Value')]
ndx <- order(t_test_results)
ttest_order <- t_test_results[ndx]
length(ttest_order)
length(ttest_order[!is.na(ttest_order)])
ttest_BH <- data.frame(pvalues=ttest_order, rank=rep(1:3541))
#set FDR to 5%
ttest_BH$BH_critical <- (ttest_BH$rank/3541)*0.05
tail(ttest_BH)
cutoff <- ttest_BH[which(ttest_BH$pvalues < ttest_BH$BH_critical),]
head(cutoff)
tail(cutoff)
#5% FDR: 0 protein


#-------------------------------------------Radial Plot------------------------------------------

#Data set
data_total <- read.csv("Total_MSK_vs_MSK_BKI.csv", header=TRUE, row.names=1)
head(data_total)

proximals <- data_total[which(data_total$adj.P.Val <= 0.05),]
nrow(proximals)#0 potential substrates

to_plot <- data_total[,c('log2FC','adj.P.Val',' Radial.plot.label',' Protein.function.process')]
to_plot_sig <- to_plot[which(to_plot$adj.P.Val <0.05),]
nrow(to_plot_sig)#0 significantly changing phosphopeptides
to_plot_sig
head(to_plot)

display.brewer.all()
colours <- brewer.pal(n=12, name='Paired')
display.brewer.pal(n = 12, name = 'Paired')
brewer.pal(n = 12, name = 'Paired')
colours

p <- ggplot(to_plot, aes(x=log2FC, y=log10(adj.P.Val),
                color=Protein.function.process,
                label=ifelse(adj.P.Val <= 0.05, as.character(paste(Radial.plot.label), size=4),'')))+

  geom_point(colour='grey40', size=4, alpha=0.5)+

  geom_point(data=to_plot[which(to_plot$adj.P.Val <0.05),],
             aes(x=log2FC, y=log10(adj.P.Val),
                 color=Protein.function.process), size=4, alpha=0.5)+

  geom_text_repel(data=to_plot[which(to_plot$adj.P.Val < 0.05),], aes(label=Radial.plot.label), size=4,
                  min.segment.length = 0,
                  seed = 42,
                  box.padding = unit(0.5, "lines"),
                  point.padding = unit(0, "lines"),
                  nudge_y = 3.5,
                  segment.alpha = 0.5,
                  force = 200,
                  force_pull = 50,
                  max.time = 1000,
                  show.legend = FALSE,
                  max.overlaps = Inf
  ) +

  coord_polar(theta = "x", start = 0, direction = 1, clip = 'on')+

  scale_x_continuous(
    limits=c(-6, 6),
    breaks = c(6, 4, 2, 0, -2, -4, -6))+

  scale_y_continuous(
    limits = c(-5, 0),
    breaks = c(-2, -3, -4)) +

  #adding the cut-off line for p-value <0.05
  geom_hline(yintercept = -1.301029995663981, colour="grey30", size=0.5, linetype='dashed')+

annotate(
  x = 4.55,
  y = -1.7,
  label = "-2",
  geom = "text",
  color = "gray12",
  size = 3,
) +
  annotate(
    x = 4.6,
    y = -2.7,
    label = "-3",
    geom = "text",
    color = "gray12",
    size = 3,
  ) +
  annotate(
    x = 4.65,
    y =-3.7,
    label = "-4",
    geom = "text",
    color = "gray12",
    size = 3,
  ) +

  #removing the original label axis and axis
  labs(y = NULL)+
  labs(x = NULL)+

  #selecting the graph theme
  theme_bw(base_size = 14)+
  theme(
    axis.text.y = element_blank(),
    axis.ticks = element_blank(),
    panel.border = element_blank()
  )

p

ggsave('Total_Radial Total_MSK_vs_MSK_BKI', dpi=1000)
ggsave('Total_Radial Total_MSK_ vs_MSK_BKI.pdf', p, width=13, height=6)
```

**2. Proximal phosphopeptides**

**2.1. Code to analyse MSK against the spatial reference KKT2**

```
library('ggplot2')
library('dplyr')
library('reshape2')
library('RColorBrewer')
library('coin')
library('ggrepel')
library('limma')
library('statmod')
library('grid')
library('gridExtra')
set.seed(42)
options(ggrepel.max.overlaps = Inf)


#-----------------------------Extracting dataset that will be compared---------------------------

Phospho <- read.csv('D548_Phos_QI_Results.csv', header=TRUE, skip=4)
head(total)
nrow(total)

#extracting the selected data (only the groups that will be compared)
Phospho_s <- (Phospho[,c('MSK_A', 'MSK_B', 'MSK_C', 'MSK_D', 'MSK_E',
                     'KKT2_A', 'KKT2_B', 'KKT2_C', 'KKT2_D', 'KKT2_E')])

rownames(phospho_s) <- phospho$Accession
head(phospho_s)
class(phospho_s)

write.csv(phospho_s, 'D548_Phos_QI_Results_Sel_n0.csv')

#Phosphopeptides with ≥2 missing values across the five biological replicates were removed
#Use =COUNTIF (range, "criterion") in excel to retain phosphopeptides with non-zero intensity in at least 3 of the 5 biological replicates.
#Save file as 'D548_Phos_QI_Results_Sel_n0.csv'


#--------------------------------------Imputing 0 values-----------------------------------------

phospho <- read.csv('D548_Phos_QI_Results_Sel_n0.csv', header=TRUE)
head(phospho)
colnames(phospho)

phospho_m <- melt(phospho[,c("MSK_A","MSK_B","MSK_C","MSK_D","MSK_E",
                             "KKT2_A","KKT2_B","KKT2_C","KKT2_D","KKT2_E")])

phospho_m$Sample_Group <- sapply(strsplit(as.character(phospho_m$variable), "(?<=_)[12345]", perl=TRUE), function(x) x[[1]])

sd_intensities <- sd(log2(phospho_m$value[which(phospho_m$value>0)]))
sd_intensities#3.12
mean_intensities <- mean(log2(phospho_m$value[which(phospho_m$value>0)]))
mean_intensities#10.42

p <- ggplot(phospho_m, aes(x=log2(value), fill=Sample_Group))+
  geom_histogram(binwidth=0.3)+
  #geom_density()
  facet_wrap(~Sample_Group, scales='free_y')+xlab('log2 peptide intensity')

p

#make a gaussian distribution with same mean and sd as the distribution of peptide intensities
n5000 <- data.frame(value=(rnorm(500, mean=3.4, sd=1.1)),variable='norm')
head(n5000)

p <- ggplot(phospho_m, aes(x=log2(value), fill=Sample_Group))+
  geom_histogram(binwidth=0.3)+
  geom_histogram(data=n5000,aes(x=value), fill='yellow', alpha=0.5,binwidth=0.3)+
  #geom_density()
  facet_wrap(~Sample_Group, scales='free_y')+xlab('log2 peptide intensity')

p

#set seed for generating random numbers from rnorm
set.seed(2021-06-13)
impute <- function(x){
  #make sure input is numeric vector
  b <- as.numeric(x)
  #count zeros in group
  n.zeros <- length(b[b==0])
  if(n.zeros > 0){
    ndx.zeros <- which(b %in% c(0))
    #generate random number from normal distribution which
    #models low abundance peptides
    imputed_value <- rnorm(n.zeros,mean=3.4,sd=1.1)
    #transform imputed values from log space
    imputed_values_intensity <- 2^(imputed_value)
    b[ndx.zeros] <- imputed_values_intensity
    return(c(b))
  }else{
    return(c(b))
  }
}

#impute missing values per sample group randomly taking number from normal distribution created to
#model the low intensity peptides
phospho_imputed <- apply(phospho[,c("MSK_A","MSK_B","MSK_C","MSK_D","MSK_E",
                         "KKT2_A","KKT2_B","KKT2_C","KKT2_D","KKT2_E")],1 ,function(x) impute(x))

#apply returns vector, so need to convert to matrix, filling by row
phospho_imputed <- matrix(phospho_imputed, ncol=10, byrow=TRUE)
head(phospho_imputed)
colnames(phospho_imputed) <- c("MSK_A","MSK_B","MSK_C","MSK_D","MSK_E",
                             "KKT2_A","KKT2_B","KKT2_C","KKT2_D","KKT2_E")
phospho_imputed <- as.data.frame(phospho_imputed)
phospho_imputed <- cbind(phospho_imputed, phospho[,c("Accession","Description","Sequence","Modifications",
                                            "pep_expect","pep_var_mod_conf",
                                            "pep_delta")])

write.csv(phospho_imputed, 'D548 imputed.csv')

#----------------------------phosphosite IDs generated by python script--------------------------
#See python code for this step


data_phosphosites <- read.csv('D548 imputed positions.csv',row.names=1, header=TRUE,stringsAsFactors = FALSE)

head(data_phosphosites)
nrow(data_phosphosites)
colnames(data_phosphosites)

#make column for phosphosite identifier, use this to aggregate
data_phosphosites$phosphosite_ID <- apply(data_phosphosites[,c('Accession','phos_position','Description')],
                                          1, function(x) paste(c(x[1], x[2], x[3]), collapse='_'))

#aggregate phosphosites by sum, this recovers the 'split' quantifications
data_phosphosites_agg <- data_phosphosites[,c("phosphosite_ID",
                             "MSK_A","MSK_B","MSK_C","MSK_D","MSK_E",
                             "KKT2_A","KKT2_B","KKT2_C","KKT2_D","KKT2_E")] %>%
  group_by(phosphosite_ID) %>%
  #sum only those sample columns
  summarise(across(.cols=contains("P_"), .fns=list(sum=sum), .names='{col}_{fn}'))
nrow(data_phosphosites_agg)#1912
head(data_phosphosites_agg)
colnames(data_phosphosites_agg) <- sapply(colnames(data_phosphosites_agg), function(x) strsplit(x, '_sum')[[1]])

#write.csv
write.csv(data_phosphosites_agg, 'D548 imputed positions aggregated.csv')


#------------------------------------Statistical analysis----------------------------------------
#limma analysis

data_phosphosites_agg <- read.csv('D548 imputed positions aggregated.csv', header=TRUE, row.names=1)

MSK_vs_KKT2 <- as.matrix(data_phosphosites_agg[,c("MSK_A","MSK_B","MSK_C","MSK_D","MSK_E",
                             "KKT2_A","KKT2_B","KKT2_C","KKT2_D","KKT2_E")])

design <- read.csv('limma_input_Phospho_MSK_vs_KKT2_model.csv', row.names=1, header=TRUE)
design <- as.matrix(design)
head(design)

contrasts <- read.csv('limma_input_Phospho_ MSK_vs_KKT2_contrasts.csv', header=TRUE, row.names=1)
head(contrasts)

rownames(MSK_vs_KKT2) <- data_phosphosites_agg$phosphosite_ID
head(MSK_vs_KKT2)
MSK_vs_KKT2 <- apply(MSK_vs_KKT2, c(1,2), function(x) log2(x))

#try eBayes options#seems to give good results
fit <- lmFit(MSK_vs_KKT2, design)
fit_contrast <- contrasts.fit(fit, contrasts)
fit_contrast <- eBayes(fit_contrast, trend=TRUE, robust=TRUE)
table_ MSK_vs_KKT2 <- topTable(fit_contrast, number=Inf)
head(table_ MSK_vs_KKT2)

#adjust p values with benjamini hochberg
t_test_results <- table_ MSK_vs_KKT2 [,c('P.Value')]
ndx <- order(t_test_results)
ttest_order <- t_test_results[ndx]
length(ttest_order)
length(ttest_order[!is.na(ttest_order)])
ttest_BH <- data.frame(pvalues=ttest_order, rank=rep(1:1912))
#set FDR to 5%
ttest_BH$BH_critical <- (ttest_BH$rank/1912)*0.05
head(ttest_BH)
cutoff <- ttest_BH[which(ttest_BH$pvalues < ttest_BH$BH_critical),]
head(cutoff)
tail(cutoff)
# 5% FDR: 233 phosphosites

write.csv(table_ MSK_vs_KKT2, 'limma_output_Phospho_ MSK_vs_KKT2.csv')


#-------------------------------------------Radial Plot------------------------------------------
#As the analysis was performed with KKT2 against MSK, the log2FC (KKT2/MSK) was transformed to lof2FC (MSK/KKT2), using Microsoft Excel.

#Data set
data_phos <- read.csv("Phospho_MSK.proximals_vs_KKT2.csv", header=TRUE, row.names=1)
head(data_phos)

proximals <- data_phos[which(data_phos$adj.P.Val <= 0.05 & data_phosl$log2FC > 0),]
nrow(proximals)#117 proximal phosphosite for MSK

to_plot <- data_phos[,c('log2FC','adj.P.Val','Radial.plot.label','Protein.function.process’,Phosphosite’)]
to_plot_sig <- to_plot[which(to_plot$adj.P.Val <0.05),]
nrow(to_plot_sig)#233 significantly changing phosphosites
to_plot_sig
head(to_plot)

display.brewer.all()
colours <- brewer.pal(n=12, name='Paired')
display.brewer.pal(n = 12, name = 'Paired')
brewer.pal(n = 12, name = 'Paired')
colours

#select colours to represent the highlights
cols <- c('Protein kinase'=colours[4],
          'Phosphatase'=colours[7],
          'Kinetochore'=colours[10],
          'Cell division'=colours[8],
          'RNA binding / Processing'=colours[6],
          'DNA binding / Processing / Repair / Replication'=colours[1],
          'Posttranscriptional regulation of gene expression=colours[9],
          'Microtubule; Proteins that interact with microtubules; Cell motility'=colours[2],
          'Nuclear pore'=colours[3],
          'Kinetoplast'=colours[11],
          'Hypothetical'=colours[5],
          'Other'=colours[12])

p <- ggplot(to_plot[which(to_plot$log2FC > 0),],
            aes(x=log2FC, y=log10(adj.P.Val),
                color= Protein.function.process,
                label=ifelse(adj.P.Val <= 0.05, as.character(paste(Radial.plot.label), size=4),'')))+

  geom_point(colour='grey40', size=4, alpha=0.5)+
  scale_color_manual(values=cols)+

  geom_point(data=to_plot[which(to_plot$log2FC>0 & to_plot$adj.P.Val <0.05),],
             aes(x=log2FC, y=log10(adj.P.Val),
                 color=Protein.function.process), size=4, alpha=0.5)+

  geom_text_repel(data=to_plot[which(to_plot$adj.P.Val < 0.05),], aes(label=Radial.plot.label), size=4,
                  min.segment.length = 0,
                  seed = 42,
                  box.padding = unit(0.5, "lines"),
                  point.padding = unit(0, "lines"),
                  nudge_y = 0.1,
                  segment.alpha = 0.5,
                  force = 200,
                  force_pull = 50,
                  max.time = 1000,
                  show.legend = FALSE,
                  max.overlaps = Inf
  ) +

  coord_polar(clip = 'off')+

  scale_x_continuous(
    limits=c(0, 10),
    breaks = c(0, 2, 4, 6, 8))+

  scale_y_continuous(
    limits = c(-8, 0),
    breaks = c(-2, -4, -6, -8)) +

  #adding the cut-off line for p-value <0.05
  geom_hline(yintercept = -1.301029995663981, colour="grey30", size=0.5, linetype='dashed')+

  annotate(
    x = 9.1,
    y = -1.7,
    label = "-2",
    geom = "text",
    color = "gray12",
    size = 3,
  ) +
  annotate(
    x = 9.15,
    y = -3.7,
    label = "-4",
    geom = "text",
    color = "gray12",
    size = 3,
  ) +
  annotate(
    x = 9.25,
    y = -5.7,
    label = "-6",
    geom = "text",
    color = "gray12",
    size = 3,
  ) +

  #removing the original label axis and axis
  labs(y = NULL)+
  labs(x = NULL)+

  #selecting the graph theme
  theme_bw(base_size = 14)+
  theme(
    axis.text.y = element_blank(),
    axis.ticks = element_blank(),
    panel.border = element_blank(),
    legend.position = "none" # to remove the legend
  )

p

ggsave('Phospho_Radial Phospho_MSK_Proximal_vs_KKT2', dpi=1000)
ggsave('Phospho_Radial Phospho_MSK_Proximal_vs_KKT2.pdf', p, width=13, height=6)
```

**2.2. Code to analyse MSK against inhibited MSK**

```
library('ggplot2')
library('dplyr')
library('reshape2')
library('RColorBrewer')
library('coin')
library('ggrepel')
library('limma')
library('statmod')
library('grid')
library('gridExtra')
set.seed(42)
options(ggrepel.max.overlaps = Inf)


#-----------------------------Extracting dataset that will be compared---------------------------

Phospho <- read.csv('D548_Phos_QI_Results.csv', header=TRUE, skip=4)
head(total)
nrow(total)

#extracting the selected data (only the groups that will be compared)
Phospho_s <- (Phospho[,c('MSK_A', 'MSK_B', 'MSK_C', 'MSK_D', 'MSK_E',
                     'MSK_BKI_A', 'MSK_BKI_B', 'MSK_BKI_C', 'MSK_BKI_D', 'MSK_BKI_E')])

rownames(phospho_s) <- phospho$Accession
head(phospho_s)
class(phospho_s)

write.csv(phospho_s, 'D548_Phos_QI_Results_Sel_n0.csv')

#Phosphopeptides with ≥2 missing values across the five biological replicates were removed
#Use =COUNTIF (range, "criterion") in excel to retain phosphopeptides with non-zero intensity in at least 3 of the 5 biological replicates.
#Save file as 'D548_Phos_QI_Results_Sel_n0.csv'


#--------------------------------------Imputing 0 values-----------------------------------------

# dataset
phospho <- read.csv('D548_Phos_QI_Results_Sel_n0.csv', header=TRUE)
head(phospho)
colnames(phospho)

phospho_m <- melt(phospho[,c('MSK_A', 'MSK_B', 'MSK_C', 'MSK_D', 'MSK_E',
                     'MSK_BKI_A', 'MSK_BKI_B', 'MSK_BKI_C', 'MSK_BKI_D', 'MSK_BKI_E')])

phospho_m$Sample_Group <- sapply(strsplit(as.character(phospho_m$variable), "(?<=_)[12345]", perl=TRUE), function(x) x[[1]])

sd_intensities <- sd(log2(phospho_m$value[which(phospho_m$value>0)]))
sd_intensities#3.11
mean_intensities <- mean(log2(phospho_m$value[which(phospho_m$value>0)]))
mean_intensities#10.40

p <- ggplot(phospho_m, aes(x=log2(value), fill=Sample_Group))+
  geom_histogram(binwidth=0.3)+
  #geom_density()
  facet_wrap(~Sample_Group, scales='free_y')+xlab('log2 peptide intensity')

p

#make a gaussian distribution with same mean and sd as the distribution of peptide intensities
n5000 <- data.frame(value=(rnorm(500, mean=3.4, sd=1.1)),variable='norm')
head(n5000)

p <- ggplot(phospho_m, aes(x=log2(value), fill=Sample_Group))+
  geom_histogram(binwidth=0.3)+
  geom_histogram(data=n5000,aes(x=value), fill='yellow', alpha=0.5,binwidth=0.3)+
  #geom_density()
  facet_wrap(~Sample_Group, scales='free_y')+xlab('log2 peptide intensity')

p

#set seed for generating random numbers from rnorm
set.seed(2021-06-13)
impute <- function(x){
  #make sure input is numeric vector
  b <- as.numeric(x)
  #count zeros in group
  n.zeros <- length(b[b==0])
  if(n.zeros > 0){
    ndx.zeros <- which(b %in% c(0))
    #generate random number from normal distribution which
    #models low abundance peptides
    imputed_value <- rnorm(n.zeros,mean=3.4,sd=1.1)
    #transform imputed values from log space
    imputed_values_intensity <- 2^(imputed_value)
    b[ndx.zeros] <- imputed_values_intensity
    return(c(b))
  }else{
    return(c(b))
  }
}

#impute missing values per sample group randomly taking number from normal distribution created to
#model the low intensity peptides
phospho_imputed <- apply(phospho[,c('MSK_A', 'MSK_B', 'MSK_C', 'MSK_D', 'MSK_E',
                     'MSK_BKI_A', 'MSK_BKI_B', 'MSK_BKI_C', 'MSK_BKI_D', 'MSK_BKI_E')],1 ,function(x) impute(x))

#apply returns vector, so need to convert to matrix, filling by row
phospho_imputed <- matrix(phospho_imputed, ncol=10, byrow=TRUE)
head(phospho_imputed)
colnames(phospho_imputed) <- c('MSK_A', 'MSK_B', 'MSK_C', 'MSK_D', 'MSK_E',
                     'MSK_BKI_A', 'MSK_BKI_B', 'MSK_BKI_C', 'MSK_BKI_D', 'MSK_BKI_E')
phospho_imputed <- as.data.frame(phospho_imputed)
phospho_imputed <- cbind(phospho_imputed, phospho[,c("Accession","Description","Sequence","Modifications",
                                            "pep_expect","pep_var_mod_conf",
                                            "pep_delta")])

write.csv(phospho_imputed, 'D548 imputed.csv')


#----------------------------phosphosite IDs generated by python script--------------------------
#See python code for this step

data_phosphosites <- read.csv('D548 imputed positions.csv',row.names=1, header=TRUE,stringsAsFactors = FALSE)
head(data_phosphosites)
nrow(data_phosphosites)
colnames(data_phosphosites)

#make column for phosphosite identifier, use this to aggregate
data_phosphosites$phosphosite_ID <- apply(data_phosphosites[,c('Accession','phos_position','Description')],
                                          1, function(x) paste(c(x[1], x[2], x[3]), collapse='_'))

#aggregate phosphosites by sum, this recovers the 'split' quantifications
data_phosphosites_agg <- data_phosphosites[,c("phosphosite_ID",
                     'MSK_A', 'MSK_B', 'MSK_C', 'MSK_D', 'MSK_E',
                     'MSK_BKI_A', 'MSK_BKI_B', 'MSK_BKI_C', 'MSK_BKI_D', 'MSK_BKI_E')] %>%
  group_by(phosphosite_ID) %>%
  #sum only those sample columns
  summarise(across(.cols=contains("P_"), .fns=list(sum=sum), .names='{col}_{fn}'))
nrow(data_phosphosites_agg)#1874
head(data_phosphosites_agg)
colnames(data_phosphosites_agg) <- sapply(colnames(data_phosphosites_agg), function(x) strsplit(x, '_sum')[[1]])

#write.csv
write.csv(data_phosphosites_agg, 'D548 imputed positions aggregated.csv')


#------------------------------------Statistical analysis----------------------------------------
#limma analysis

data_phosphosites_agg <- read.csv('D548 imputed positions aggregated.csv', header=TRUE, row.names=1)

MSK_vs_MSK_BKI <- as.matrix(data_phosphosites_agg[,c('MSK_A', 'MSK_B', 'MSK_C', 'MSK_D', 'MSK_E',
                     'MSK_BKI_A', 'MSK_BKI_B', 'MSK_BKI_C', 'MSK_BKI_D', 'MSK_BKI_E')])

design <- read.csv('limma_input_Phospho_MSK_vs_MSK_BKI_model.csv', row.names=1, header=TRUE)
design <- as.matrix(design)
head(design)

contrasts <- read.csv('limma_input_Phospho_ MSK_vs_MSK_BKI_contrasts.csv', header=TRUE, row.names=1)
head(contrasts)

rownames(MSK_vs_MSK_BKI) <- data_phosphosites_agg$phosphosite_ID
head(MSK_vs_MSK_BKI)
MSK_vs_MSK_BKI <- apply(MSK_vs_MSK_BKI, c(1,2), function(x) log2(x))

#try eBayes options#seems to give good results
fit <- lmFit(MSK_vs_MSK_BKI, design)
fit_contrast <- contrasts.fit(fit, contrasts)
fit_contrast <- eBayes(fit_contrast, trend=TRUE, robust=TRUE)
table_MSK_vs_MSK_BKI <- topTable(fit_contrast, number=Inf)
head(table_MSK_vs_MSK_BKI)

#adjust p values with benjamini hochberg
t_test_results <- table_MSK_vs_MSK_BKI[,c('P.Value')]
ndx <- order(t_test_results)
ttest_order <- t_test_results[ndx]
length(ttest_order)
length(ttest_order[!is.na(ttest_order)])
ttest_BH <- data.frame(pvalues=ttest_order, rank=rep(1:1874))
#set FDR to 5%
ttest_BH$BH_critical <- (ttest_BH$rank/1874)*0.05
head(ttest_BH)
cutoff <- ttest_BH[which(ttest_BH$pvalues < ttest_BH$BH_critical),]
head(cutoff)
tail(cutoff)
##5% FDR: 4 phosphosites

write.csv(table_MSK_vs_MSK_BKI, 'limma_output_Phospho_MSK_vs_MSK_BKI.csv')


#-------------------------------------------Radial Plot------------------------------------------

#Data set
data_phos <- read.csv("Phospho_MSK_vs_MSK_BKI.csv", header=TRUE, row.names=1)
head(data_phos)

proximals <- data_phos[which(data_phos$adj.P.Val <= 0.05),]
nrow(proximals)#4 potential substrates

to_plot <- data_phos[,c('log2FC','adj.P.Val',' Radial.plot.label',' Protein.function.process')]
to_plot_sig <- to_plot[which(to_plot$adj.P.Val <0.05),]
nrow(to_plot_sig)#4 significantly changing phosphopeptides
to_plot_sig
head(to_plot)

display.brewer.all()
colours <- brewer.pal(n=12, name='Paired')
display.brewer.pal(n = 12, name = 'Paired')
brewer.pal(n = 12, name = 'Paired')
colours

#select colours to represent the highlights
cols <- c('ARFGAP1 _ S155'=colours[8],
          'Rab3GAP _ S804'=colours[5],
          'LmxM.36.2560 _ S552'=colours[4],
          'NSP _ T484'=colours[2])

p <- ggplot(to_plot, aes(x=log2FC, y=log10(adj.P.Val),
                color=Protein.function.process,
                label=ifelse(adj.P.Val <= 0.05, as.character(paste(Radial.plot.label), size=4),'')))+

  geom_point(colour='grey40', size=4, alpha=0.5)+
  scale_color_manual(values=cols)+


  geom_point(data=to_plot[which(to_plot$adj.P.Val <0.05),],
             aes(x=log2FC, y=log10(adj.P.Val),
                 color=Protein.function.process), size=4, alpha=0.5)+

  geom_text_repel(data=to_plot[which(to_plot$adj.P.Val < 0.05),], aes(label=Radial.plot.label), size=4,
                  min.segment.length = 0,
                  seed = 42,
                  box.padding = unit(0.5, "lines"),
                  point.padding = unit(0, "lines"),
                  nudge_y = 0.1,
                  segment.alpha = 0.5,
                  force = 200,
                  force_pull = 50,
                  max.time = 1000,
                  show.legend = FALSE,
                  max.overlaps = Inf
  ) +

  coord_polar(theta = "x", start = 0, direction = 1, clip = 'on')+

  scale_x_continuous(
    limits=c(-5, 5),
    breaks = c(4, 2, 0, -2, -4))+

  scale_y_continuous(
    limits = c(-5, 0),
    breaks = c(-2, -3, -4)) +

  #adding the cut-off line for p-value <0.05
  geom_hline(yintercept = -1.301029995663981, colour="grey30", size=0.5, linetype='dashed')+

annotate(
  x = 4.55,
  y = -1.7,
  label = "-2",
  geom = "text",
  color = "gray12",
  size = 3,
) +
  annotate(
    x = 4.6,
    y = -2.7,
    label = "-3",
    geom = "text",
    color = "gray12",
    size = 3,
  ) +
  annotate(
    x = 4.65,
    y =-3.7,
    label = "-4",
    geom = "text",
    color = "gray12",
    size = 3,
  ) +

  #removing the original label axis and axis
  labs(y = NULL)+
  labs(x = NULL)+

  #selecting the graph theme
  theme_bw(base_size = 14)+
  theme(
    axis.text.y = element_blank(),
    axis.ticks = element_blank(),
    panel.border = element_blank()
  )

p

ggsave('Phospho_Radial Phospho_MSK_vs_MSK_BKI', dpi=1000)
ggsave('Phospho_Radial Phospho_MSK_ vs_MSK_BKI.pdf', p, width=13, height=6)
```

**CUSTOM PYTHON CODE TO GENERATE PHOSPHOSITES ID**

```
#to get phosphosite or phosphorylated region from list of accession numbers
#used proteome from TriTrypDB version 54.

import csv
import re

#open the Lmex fasta file and make it into a dictionary, key is accession, value is protein sequence
fasta = open('/Users/JulianaCarnielli/TriTrypDB-54_LmexicanaMHOMGT2001U1103_AnnotatedProteins.fasta','r')
acc_seq = {}
sequence = ''
gene_id = ''
for line in fasta:
    header_search = re.search('>L', line)
    #found gene header
    if header_search != None:
        #dump previous gene id and sequence into dictionary
        if gene_id != '':
            acc_seq[gene_id] = sequence
            sequence = ''
            gene_id = ''
        #find the gene accession
        split_line = line.split('|')
        for item in split_line:
            gene_search = re.search('gene=', item)
            if gene_search != None:
                item_split = item.split('=')
                gene_id = item_split[1].rstrip()
            else:
                pass
    #not header so part of protein sequence
    else:
        seq_chunk = line.rstrip()
        #add sequence to previous chunk
        sequence = sequence + seq_chunk
#the last entry
acc_seq[gene_id] = sequence

fasta.close()

#open the imputed phospho data set
file = open('/Users/JulianaCarnielli/D548 imputed.csv','r')

#output
file_out = open('/Users/JulianaCarnielli/D548 imputed positions.csv','w',newline='')
output = csv.writer(file_out)
def getModPosition(accession, peptide, mod_pos, mod_pos_conf):
    #get the protein sequence for accession
    try:
        protein = acc_seq[accession]
    except KeyError:
        protein = 'NA'
    #get position of first AA in peptide within protein
    pep_start = protein.find(peptide)
    #only get position of confident localisations
    if int(mod_pos_conf) >=75:
        mod_position_num = pep_start + int(mod_pos)
        residue = peptide[int(mod_pos) - 1]
        mod_position = residue + str(mod_position_num)
    #or get list of possible modified residues
    else:
        mod_residues = []
        res_count = 0
        for residue in peptide:
            res_count = res_count + 1
            if residue == 'S' or residue == 'T':
                position = res_count + pep_start
                #denote uncertain mods with asterisk
                mod_residues.append(residue + str(position) + '*')
        mod_position = ';'.join(mod_residues)

    return mod_position

data = csv.reader(file)
#get accessions and put in a list
accessions = []
row_count = 0
acc_index = 0
for row in data:
    if row_count == 0:
        acc_index = row.index('Accession')
        mod_index = row.index("Modifications")
        conf_index = row.index("pep_var_mod_conf")
        seq_index = row.index("Sequence")
        output.writerow(row+['phos_position'])
        row_count = row_count + 1
    else:
        #accession = row[acc_index]
        split_accession = row[acc_index].split(';')
        #if there are multiple accessions for a protein id, sort them and take first one
        accessions = sorted(split_accession)
        accession = accessions[0]
        mods = row[mod_index].split('|')
        positions = []
        for mod in mods:
            phos_search = re.search('Phospho', mod)
            if phos_search != None:
                mod_position_pep = mod.split('[')[-1].split(']')[0]
                if row[conf_index] == '':
                    #no confidence value so only one possible position
                    conf = 100
                else:
                    conf = int(row[conf_index].split('%')[0])
                mod_position = getModPosition(accession, row[seq_index], mod_position_pep, conf)
                positions.append(mod_position)
        #convert list to set to get only unique positions
        positions_s = set(positions)
        positions_str = ';'.join(positions_s)
        output.writerow(row + [positions_str])
        row_count = row_count + 1

file.close()
file_out.close()
```
